# Systematic engineering and machine learning analysis of intrinsic terminators reveal crucial nucleotides directly upstream of the terminator hairpin

**DOI:** 10.64898/2026.07.06.736697

**Authors:** Charlotte C. Koster, Barbara Terlouw, Thijs Nieuwkoop, Sjoerd C.A. Creutzburg, Maria Martin-Pascual, Miguel Paredes-Barrada, Panagiotis Kopsiaftis, Hans G.H.J Heilig, Theo van Laar, John van der Oost, Nico J. Claassens

**Affiliations:** Laboratory of Microbiology, Wageningen University & Research, Wageningen, the Netherlands; Bioinformatics Group, Wageningen University & Research, Wageningen, the Netherlands; Institute for Complex Molecular Systems (ICMS), Department of Biomedical Engineering, Eindhoven University of Technology, Eindhoven, The Netherlands; Systems and Synthetic Biology Laboratory, Wageningen University & Research, Wageningen, the Netherlands; Corbion, Gorinchem, The Netherlands; Department of Bionanoscience, Delft University of Technology, Delft, the Netherlands

## Abstract

Transcriptional termination efficiency is considered an important parameter for finetuning bacterial gene expression. Still, the design principles that determine transcription termination efficiency remain poorly understood. In this study, we aimed to investigate the impact of the 3’ untranslated region (3’UTR) on gene expression in *Escherichia coli* and other bacteria. First, 3’UTR variant sequences were generated, with randomized 30 bp sequences inserted between the STOP-codon and an intrinsic terminator, consisting of a GC-rich hairpin and a downstream poly(U)-tail. Using three reporter genes, it was found that different 3’UTR sequences resulted in an up to five-fold difference in protein production, independent of the upstream coding sequence. The highest protein production was achieved when an adenosine was present directly upstream of the terminator hairpin. This was consolidated by systematic substitution of key nucleotides of the terminator and assessing their effect on mRNA and protein levels. Subsequently, we developed a predictive random forest machine learning model trained on the termination efficiency of different natural and synthetic terminator sequences, revealing an important role for the nucleotides directly upstream of the terminator hairpin. Altogether, this study showed that an additional adenosine nucleotide upstream of the terminator hairpin leads to improved protein production while reducing terminator read-through.

**GRAPHICAL ABSTRACT:** 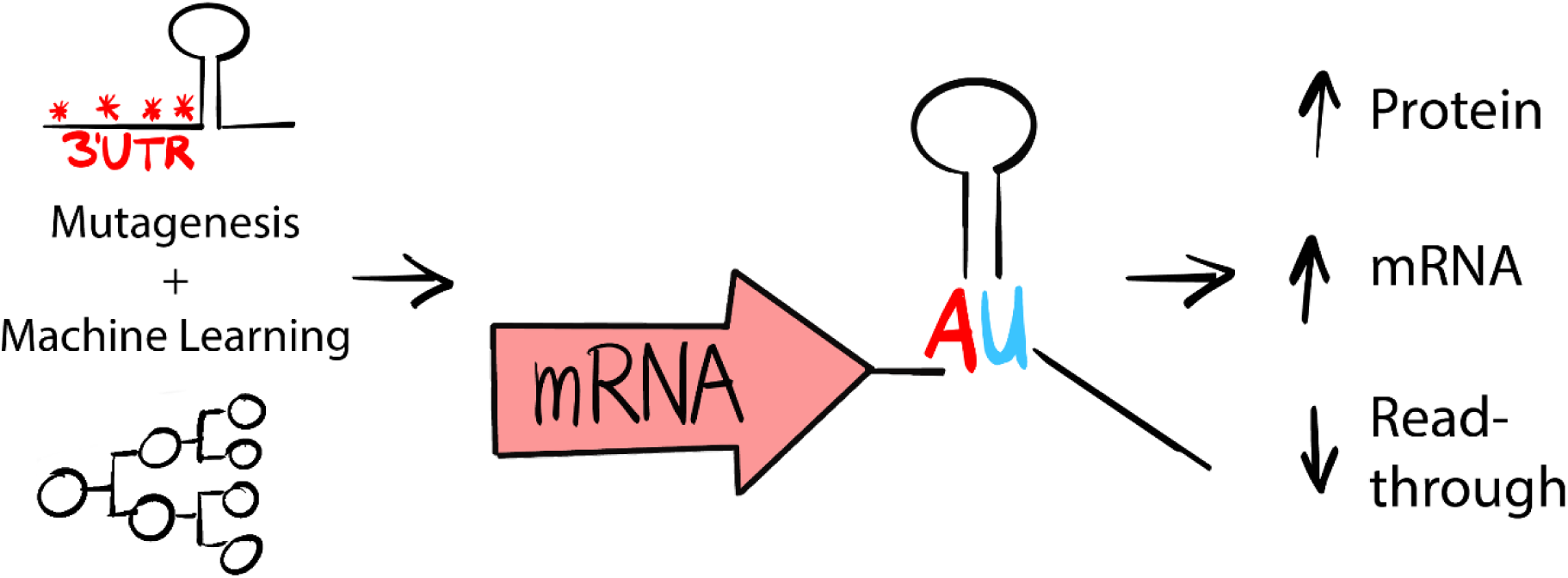

## INTRODUCTION

The mRNA sequence of a gene has a profound effect of the level of protein levels that the corresponding gene encodes. This is especially relevant when designing a synthetic gene for heterologous protein production in *E. coli* or other bacteria. Aside from the coding region of mRNA, where translation efficiency may be linked to codon usage (1), the 5’ and 3’ untranslated regions (UTRs) of mRNA transcripts strongly influence how much protein is produced. While there is an abundance of studies focusing on the effect and secondary structure of the 5’UTR (including the ribosome binding site) and the coding sequences for maximizing or tuning protein production ((2–5), extensively reviewed by (6–9), the contribution of the 3’UTR is less explored. The 3’UTR influences transcript abundance through transcription termination, either mediated through pyrimidine-rich ‘Rho utilisation’ (rut) sites in Rho-dependent terminators (10), or through G-C rich mRNA hairpins followed by a poly(U)-tract in intrinsic terminators (11, 12). Intrinsic terminators are generally used for heterologous gene expression, as these are more predictable and are used in expression vectors and/or available as standardized cloning parts. For example, the strong intrinsic T1 terminator, derived from the *rrnB* gene (13), is frequently used to terminate transcription. The termination efficiencies of libraries of intrinsic terminators have been determined experimentally in some instances (14–17), and a variety of bioinformatic prediction tools of intrinsic termination efficiency are available (18–20).

The ability to rationally predict and tune termination efficiency of intrinsic terminators is an important feature in synthetic biology. While the optimization of termination efficiency can significantly improve gene expression, tuning at terminator level is rarely considered for the heterologous expression of a synthetic construct. More than half of native intrinsic terminators in *E. coli* have a termination efficiency of less than 95 % (21). Incomplete termination has an important regulatory effect in the cell, i.e. the tuning of relative transcription levels within multi-gene operons (22), and the regulation of the level of reinitiation by the RNA polymerase (RNAP) in the transcription bubble (23). However, incomplete termination can have negative effects in heterologous gene expression, as transcriptional read-through can lead to unwanted expression of downstream genes, wasting cellular resources, and lower transcription rates due to run-off RNAP complexes. Additionally, mRNA secondary structure of the 3’UTR also impacts transcript stability and mRNA decay rates, which in turn can influence protein production rates.

For accurate control in synthetic biology, including the emerging field of synthetic genomics , it is essential to be able to accurately control genome-wide gene expression. For instance, strong terminators could be used to boost protein production rates, or terminators could be used to tune metabolic pathways by asymmetrical transcription in multi-gene operons, with genes separated by terminators of suboptimal strength . Such approaches are rarely applied in bioengineering efforts. The underutilization of terminator parts for transcriptional control represents a reservoir of underexplored future potential in synthetic biology.

The proposed mechanism by which intrinsic terminators do terminate transcription has been extensively studied (11, 15). In short, upon transcription of the terminator, the RNA transcription by RNAP stalls due to the stretch of weak poly-A-U DNA-RNA base pairing at the poly(U)-tail of the transcribed terminator, which destabilizes the RNAP polymerase complex. This pause in transcription allows time for the upstream GC-rich stem-loop of the transcribed terminator to start forming inside the RNA exit tunnel of the RNAP. It is then hypothesized that the formation of the hairpin puts strain on the DNA-RNA duplex at the poly-A-U stretch, causing the RNAP to dissociate from the template strand, and to release the nascent mRNA. Although the poly(U)-tail and strength of the hairpin are key factors in termination efficiency, the spacing between the stop codon and the terminator hairpin also affects the transcription termination efficiency. Li *et al*., observed that spacing sequences of less than ±20 base pairs causes repression of the terminator efficiency due to sterically hindrance of co-translationally active ribosomes (24). However, the effect of the nucleotide identity, particularly on translation efficiency, has not yet been studied in detail. Besides aiding transcription termination, the strong GC-rich stem structure stabilizes the mRNA by protecting it against RNase II, RNase R and PNPase degradation (25).

While the mechanisms by which intrinsic termination controls transcription and translation are quite well understood, our understanding of the role of the 3’UTR nucleotide sequence is limited. For example, it is not known how the base composition of the hairpin, the poly(U)-tail, the 3’UTR region upstream of the hairpin, and any other upstream- or downstream sequence elements influence termination efficiency. This was demonstrated by Cambray *et al.*, who showed a high dependency on sequence context beyond the terminator hairpin and poly(U)-tail alone, including the sequences surrounding the terminator (26). In addition, there is evidence that intrinsic termination is also influenced by extrinsic (protein) factors, such as NusA,

NusG or bacteriophage λ N factor in *E. coli*, and NusA, NusG and *rho* factor in *Bacillus subtilis*. These proteins may interact with certain sequence elements, affecting termination efficiency (27–30). Furthermore, as pointed out by Dierksheide *et al.*, studying native terminators is impeded by the extreme sequence dissimilarity of these terminators and sparsity of the naturally covered sequence space, making it challenging to inform which sequence elements contribute to termination efficiency from native sequences (21). As indicated by the authors, gaining such mechanistic insight will require small single-nucleotide perturbation experiments, assessing termination efficiency for each variant. Especially the role of the sequence upstream of the terminator hairpin is elusive. This region is routinely referred to as the (A)-tract, as this region often contains a stretch of adenosine nucleotides (15, 20). Although the (A)-tract is mostly associated with bidirectionality of terminators (11), there is evidence for a role of the (A)-tract in enhancing termination efficiency (15, 31). As this region is not consistently A-rich we refer to this region further as ‘3’UTR spacer sequence’, which in the definition of this work includes all the bases between STOP-codon and start of the G-C hairpin.

In this study, we aimed to increase understanding of the molecular details of the transcription termination process. We specifically focused on the 3’UTR spacer sequence as there is limited data on the role and exact sequence design requirements of this sequence on termination efficiency. By a combination of complete randomization of this sequence and single-nucleotide variant libraries, we set out to gain insight in the sequence-function relationship of terminators. As such, we identified specific mutations that can double termination efficiency and protein production levels.

## MATERIAL AND METHODS

### Strain and culture conditions

Generally, experiments were performed using *E. coli* K-12 DH10B (New England Biolabs [NEB], C3019H). *E. coli* K-12 BW25113 strains harbouring single gene knockouts in mRNA turnover genes originated form the KEIO collection (32), with the genomic Kanamycin resistance removed using plasmid pCMT-flp (a gift from Mark Liles; Addgene plasmid #6727 (33)), and parental strain *E. coli* K-12 BW25113 as control (34). Routinely, bacteria were grown at 37 °C in Lysogeny Broth medium (LB; 1 % NaCl, 0.5 % yeast extract, and 1 % tryptone) or on LB agar plates, where LB was supplemented with 1, 5 % agar. Fluorescence assays were performed in M9TG medium (1X M9 media salts [Sigma Aldrich], 10 g/L Trypton, 5 g/L glycerol) which allows for high cell density and low auto-fluorescence. Kanamycin (Kan) resistant transformants were selected on media supplied with 50 μg/mL kanamycin (Sigma Aldrich). Bacterial cultures were grown at 37 °C, and cultures were stored as long-term in aliquots in LB + Kan supplied with 30 % glycerol (v/v) at -70 °C.

### General molecular biology techniques

All cloning products were transformed and maintained in *E. coli* K-12 DH10B heat-shock competent cells (NEB, C3019H). Unless indicated otherwise, PCR was performed using Q5 High-Fidelity 2X Master Mix (NEB, M0492L) according to manufacturer’s instructions. PCR products were purified and concentrated using DNA clean and concentrator-5 (Zymo Research, D4004) or Zymoclean Gel DNA Recovery Kit (Zymo Research, D4002). Plasmid isolation was performed using the QIAprep Spin Miniprep Kit (Qiagen, 27106). Plasmid sequences were verified by EZ-Seq Sanger sequencing (Macrogen Europe) and Nanopore whole-plasmid sequencing (Plasmidsaurus). Primers and ssDNA oligos were generally ordered as desalted ssDNA oligos (IDT) and are indicated with a BGXXXXX number where relevant, all primer sequences used in this study are reported in Supplementary Table S2. Apart from the 3’spacer library generation, Golden Gate cloning was performed using 25 fmol backbone, 50 fmol insert, 1X T4 buffer (NEB, B0202S), 200U T4 ligase (NEB, M0202S) and 5U BsmBI-v2 (NEB, R0739S) in 25 cycles of digestion at 37 °C and annealing at 16 °C. Final digestion and enzyme inactivation was omitted due to an internal BsmBI site in the kanamycin gene. All genetic sequences are listed in the supplementary information.

### Plasmid assembly

*3’ spacer library generation.* The 3’ spacer library was ordered at Integrated DNA Technologies (IDT, ) as single-stranded DNA (ssDNA) oligos containing 30 fully degenerate nucleotides (Nx30) flanked on both sides with 4 nucleotide overhang and BsaI recognition sites (BG18017, Table S2). The ssDNA oligos were converted to double stranded DNA by PCR amplification using 400 pmol primer BG18018 and 200 pmol ssDNA in 50 μL reachtion using OneTaq Master Mix (NEB, M0482S), with the following cycling conditions: annealing at 50 °C and elongation at 72 °C for 5 s, repeated for 99 cycles. The backbones containing GFPuv and mRFP1 (See supplementary data for all sequences) were prepared by first inserting a substantial piece of nonsense DNA flanked by outward facing BsaI sites between the stop codon and terminator which allowed for more precise gel separation later on. The plasmids were sequence verified and pre-digested using BsaI-HFv2 (NEB, R3733S) to reduce transformation background. The digested backbone was separated on an agarose gel and purified. The dsDNA was inserted into the backbone using a Golden Gate Assembly Kit (BsaI-HF®v2 [NEB, E1601S]) with a 3:1 ratio. 300 colonies were picked and the fluorescence quantified using a Attune NxT Flow Cytometer (Thermo Fisher Scientific). 96 cultures covering the full fluorescent range were reinoculated in liquid M9TG + kanamycin. The fluorescence of the cultures was measured again and the associated plasmids were isolated and sent for Sanger sequencing.

*Spacer variant generation.* Ten spacer variant plasmids from the library were selected ordered as ssDNA oligos [BG20363-BG20382] with staggered ends upon annealing. ssDNA was annealed by mixing the oligos in an equimolar ratio. The oligos were boiled at 95 °C for 5 minutes in a thermocycler and then cooled down at a temperature ramp of 0.3 °C/s. The annealed dsDNA oligos were inserted in the respective backbones as indicated before and sequence verified using Sanger sequencing.

### Rational variant generation

The plasmid harbouring spacer 3 was used as a template for generation of the rational 3’UTR variants. The plasmid was linearized by PCR including the spacer 3, introducing BsmBI sites for Golden Gate cloning of the terminator inserts. The terminator inserts were ordered as ssDNA oligos (IDT) and annealed as described before. The inserts were inserted in the linearized plasmids via Golden Gate assembly. Plasmids were verified using Nanopore sequencing.

### Plasmids for read-through assays

All plasmid variants tested in the read-through assay (spacers1-10, (A)-tract variants 1, 3, 7, extended hairpin variants 1, 3, 7 and (U)-tail variants 0, 4, 6) were linearized by PCR, introducing BsmBI sites. The superfolder GFP (sfGFP) ORF including T0 terminator was amplified by PCR introducing BsmBI restriction sites. Terminator L3S2P52 from Chen *et al.*, was chosen as strong second terminator and was ordered as two partial ssDNA oligos BG36338/9 including BsmBI sites. The Mango-III(10AU) was chosen due to its low signal-to-noise ratio (35), and ordered as two partial ssDNA oligos including BsmBI sites BG37684/5. These oligos were turned into a dsDNA fragment by overlap extension PCR using only the two ssDNA oligos as templates. The backbone was linearized as described above, a backbone omitting the terminator was included as a positive control for Mango-III experiments. GFP, a second terminator or Mango-III were cloned behind the different 3’UTR variants by Golden Gate assembly as described above. For Mango-III experiments, the full expression cassette (P*bla,* mRFP, 3’UTR) was linearized for the transcription assay using primers BG38694/38703, introducing an additional stabilizing terminator behind the Mango-III aptamer.

### Growth and fluorescence measurements

Strains were grown in a pre-culture of M9TG+Kan in a 96 wells 2 mL Masterblock (Greiner Bio-one, 780280). The block was covered with a gas-permeable membrane and incubated overnight at 37°C. GFP fluorescence was measured at an excitation wavelength of 488/9 nm and emission wavelength of 508/90 nm, with a gain of 50-75. RFP fluorescence was measured at an excitation wavelength of 584/9 nm, and an emission wavelength of 607/9nm, with a gain of 100. All fluorescence was measured in black μClear 96-well plates (Greiner Bio-one, 655096). Optical density was measured as culture absorbance at a wavelength of 600 nm (OD600).

### Endpoint measurements

Pre-cultures were diluted 10000X in 200 μL M9TG and grown in the same way as the pre-cultures. The cultures were then cooled down to room temperature and diluted 5x in 1x PBS pH 7.4, and fluorescence of 100 μL of the dilution was measured with a BioTek Synergy MX microplate reader (Agilent). Fluorescence was calculated as raw fluorescence per OD600 for 100 μL 5x dilution.

### Kinetic measurements

Pre-cultures were diluted 200X in 200 μL M9TG and grown in a BioTek SynergyH1 or a BioTek Synergy Neo2 plate reader (Agilent) at 37 °C and continuous double orbital shaking at 282 cpm. Fluorescence was calculated as raw fluorescence over OD600.

### LacZ assay measurements

LacZ activity was assayed in E. coli DH10B T1R in triplicate using a Miller assay as described by

(36). In short, after overnight growth at 37 °C, 20 μL of culture was mixed with 80 μL of permeabilization solution and incubated at 30 °C for 30 min. 600 μL of pre-warmed substrate solution was added and incubated at 30°C until sufficient colour had developed. 700 μL of stop solution was added to quench the reaction. The reaction was filtered through a 0.2 μM filter and measured in a spectrophotometer at 420 nm in a 1 cm cuvette.

### Total RNA isolation

For total RNA isolation, cells were grown overnight in a 5 mL LB-Kan. Cells were diluted 1:1000 and grown until late exponential phase. A volume corresponding to approximately 10^9^ cells were pelleted at 4 °C at 3, 600 ×*g* for 10 minutes, medium was decanted, and pellets were flash frozen in liquid nitrogen and stored at -80 °C until further processing. Total RNA was isolated using a Monarch Total RNA Miniprep kit (NEB, T2010S), including DNaseI processing, with the following modifications. Frozen pellets were directly resuspended in 200 μL DNA/RNA protection reagent containing 1 mg/mL Lysozyme solution (Thermo Fisher Scientific, #900082), and RNA was eluted in 60 μL nuclease-free water. When needed, RNA was concentrated further by vacuum centrifugation in a Concentrator plus vacuum centrifuge (Eppendorf) for 20 min at 30 °C at the V-AQ setting. Purity of RNA was assessed measuring absorbance ratios at 260/280 nm (∼2.0) and 260/230 nm (2.0-2.2) using a NanoPhotometer UV-Vis spectrophotometer (Implen). Integrity was assessed by separation of denatured RNA (containing 1X RNA loading dye [NEB, B0363S], heated to 70 °C for 5 minutes, directly placed on ice) on a 1 % agarose gel for and visual inspection of the presence of tRNA, 16S rRNA and 23s rRNA bands, and absence of degradation products. Quantity was determined using the Qubit RNA BR Assay Kit (Thermo Fisher Scientific, Q10210). RNA was stored at -70 °C until further analysis.

### Reverse transcription digital droplet PCR (RT-ddPCR)

First strand cDNA synthesis was performed using GoScript Reverse Transcription Mix containing random primers (Promega, #9PIA280) according to the manufacturer’s protocol. 2 μg of total RNA, isolated from three biological replicate cultures per strain, was used per reaction, denatured at 70 °C for 5 min and immediately placed on ice before adding it to the reverse transcription reaction. Three biological replicates were used per strain. The RNA-cDNA product was diluted 100x in nuclease-free water and directly used as template for digital droplet PCR (ddPCR). Primers were designed to contain a GC-clamp, have a Tm of approximately 60 °C and amplify 100-150 base pair amplicon in the 3’ end of the coding sequence (BG36332/3 for RFP, BG36334/5 for FBA1). A reaction mix containing 1X QIAcuity EvaGreen Mastermix (Qiagen, 250111), 0.75 μM forward primer, 0.75 μM reverse primer was added to 2 μL of 100X diluted RNA-cDNA and loaded on a 96-wells 8.5K Qiacuity Nanoplate (Qiagen, 250021) and analyzed on a QIAcuity digital PCR instrument (Qiagen). A No-Template Control omitting cDNA was included in each run. The plate was run using the EvaGreen priming profile with a Tm of 60 °C. The resulting number of copies/mL of RFP mRNA were divided by the copies/mL of household gene FBA to arrive at a normalized number of RFP mRNA copies/mL.

### Term-seq RNA sequencing

A Term-seq protocol was designed based on protocols by Dar *et al.* and Choe *et al.* (37, 38). RNA was isolated from three biological replicates as described above. All samples were equilibrated to 0.5 μg/μL. All materials were cleaned with RNaseZap (Thermo Fisher Scientific, AM9780). All steps were performed in RNase-free consumables and buffers, on ice, unless indicated otherwise. Intermediate cleanup steps were performed using Mag-Bind® TotalPure NGS magnetic beads (Omega Bio-tek, M1378-00). A detailed Term-seq protocol is described in the supplementary information. In short, i5 barcodes for multiplexing compatible with Illumina Nextera Next-generation sequencing kits (Table S2, supplementary data SD_15) were ligated to the 3’end of total RNA using T4 ssRNA ligase 1 (NEB, M02040L). rRNA was depleted using a NEBNext rRNA depletion kit for bacteria (NEB, E7850S) according to manufacturer’s protocol. rRNA depleted RNA was fragmented using NEBNext Magnesium RNA Fragmentation Module (NEB, E6150S). First strand cDNA synthesis was done using Protoscript II First Strand cDNA Synthesis Kit (NEB, E6560S) according to the manufacturer’s protocol. A universal i7 DNA adapter for multiplexing compatible with Illumina Nextera Next-generation sequencing kits was ligated to the cDNA using T4 ssDNA ligase 1 (NEB, M02040L). cDNA pools were amplified with a universal primer for the i5 index and specific primers binding the universal i7 adapter (Table S2, Supplementary data SD_15) and containing a barcode, introducing unique i7 barcodes to each of the pooled cDNA samples using NEBNext Ultra II Q5 Master Mix (NEB, M0544S). Libraries were sent for short-read (150 bp) paired-end sequencing using Illumina NovaSeq X plus sequencing chemistry, performed at Novogene, UK. Raw data is stored at the Bioprojects/Sequence Read Archive (SRA, NCBI) under BioProject ID PRJNA1472557. The resulting fastq files were processed using a the European Galaxy server (https://usegalaxy.eu/, (39)). Fastq files and a FASTA reference of the *E. coli* DH10b genome and the different plasmids containing 3’UTR variants were aligned using the bwa_mem2 algorithm (version 2.2.1). The Galaxy workflow is deposited online, together with all .fasta files used for alignment (Zenodo, https://doi.org/10.5281/zenodo.20717389). Coverage was visualized using in R (version 4.4.1) using the BioConductor Rsamtools package (40)

#### *In vitro* transcription assay

*Protein production. E. coli* RNA polymerase was produced from plasmid pRM756 (41), sigma 70 was produced from plasmid RpoD_b3067 (42), NusA was produced from pExp-NusA, which was a gift from Marko Hyvönen (Addgene plasmid #112571), NusG, GreA and GreB were expressed from pET28 plasmids. GreA and GreB were included in all IVT reactions, are part of the RNAP elongation complex and improve *in vitro* transcription (43). Proteins were produced in *E. coli* BL21-Codon Plus (DE3) (Agilent, #230245), induced in 400 μM IPTG, grown overnight at 17 °C and 180 RPM. Cells were resuspended in lysis buffer (see supplementary methods for compositions). Bacteria were lysed by sonication on ice with an amplitude of 40 % for 2.5 min, 5s on, 5s off). Lysate was spun at 12, 000 *×g* for 20 min at 4 °C and supernatant was used for protein purification using affinity chromatography. Details on affinity chromatography procedures are described in the supplementary data. Final protein concentration was determined using Quick Start Bradford Dye Reagent (Bio-Rad, 5000205).

*In vitro transcription assay.* All steps were performed on ice in RNase-free conditions. RNAP was added to σ70 (RpoD) in a 1:5 ratio, and incubated for 10 min at 30 °C, then directly placed on ice, to form holo-enzymes. All linearized DNA templates were equilibrated to 200 ng/μL. *In vitro* transcription occurred in IVT buffer (40 mM Tris-HCl, 150 mM KCl, 10 mM MgCl2 5 mM DTT, 0.01 % Triton X-100, 4.5 % glycerol) containing 0.5 mM of each ribonucleotide rATP, rCTP, rUTP, rGTP (NEB, N0450S), 500 nM TO1-3PEG-Biotin Fluorophore (Applied Biological Materials Inc., G7955), 20 nM RNasin (Promega, N2511), 50 nM RNAP holoenzyme, 1 μM GreA, 1 μM GreB. Each reaction was supplemented with 1 μg of DNA template, and where applicable, 1 μM of NusA and/or 1μM of NusG. *In vitro* assays were performed in 50 μL in a black μClear 384-well plate (Greiner Bio-one, 781096). Fluorescence was measured every 6 minutes using a Synergy Neo2 plate reader (BioTek) at 37 °C with 5 s orbital shaking prior to each measurement, at an excitation wavelength of 505/10 nm and emission wavelength of 535/10 nm, with a gain of 100. The Mango-III transcription rate was calculated using linear regression with a sliding window of six to ten time points, optimizing for a linear relationship with a maximum R^2^ value. Fold-change rate was calculated by dividing the rate of a condition containing NusA and/or NusG by the rate observed when no NusA and NusG were added.

### Computational methods

All code needed to replicate data parsing, featurization, model training, and feature inference can be found in the MEWTWO (mRNA Expression Wizard 2) repository at Zenodo (doi: 10.5281/zenodo.20714791; GitHub: https://github.com/BTheDragonMaster/mewtwo).The code is implemented in Python (v3.14.0).

#### RNA secondary structure predictions

RNA secondary structures were predicted using the ViennaRNA RNAfold WebServer and ViennaRNA package (44). RNA secondary structures were visualized with forna (45).

#### Datasets

Our models were trained based on (subsets of) two datasets of terminators and their cognate predicted secondary structures and termination efficiencies: a dataset of synthetic (265) and natural (317) terminators tested in *E. coli* using a fluorescent reporter gene read-through system (further referred to as the “Chen dataset”, (15)) and a dataset of *B. subtilis* (1214) and *E. coli* (691) natural terminators based on Term-Seq data (37), previously processed by the authors of TERMITe (further referred to as the “TERMITe dataset”, (17)). As the Term-Seq data comprised multiple replicates per organism, we only chose the replicate with the greatest number of reported terminators for each species (‘(d)’ for *B. subtilis* and ‘(a)’ for *E. coli*) to prevent duplicate and highly similar datapoints in our datasets.

For TERMITe terminators, we used the termination efficiencies as reported in the dataset. For terminators from Chen *et al.*, termination strengths were reported, which we converted to termination efficiencies with the following formula:

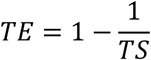

Where *TE* is the termination efficiency (fraction of transcription blocked), and *TS* is the termination strength.

For terminators from the Chen dataset, RNAFold secondary structures were available. For terminators from the TERMITe dataset, both RNAFold and TransTermHP RNA secondary structure predictions were available. For the latter, when both an RNAFold and a TransTermHP prediction were given for a terminator, the RNAFold prediction was used for hairpin determination.

Only terminators with termination efficiencies between 0.0 and 1.0 (inclusive), with available secondary structures, and with a single hairpin were kept. This yielded a final dataset of 313 natural and 265 synthetic terminators from the Chen dataset, and 1158 *B. subtilis* and 538 *E. coli* terminators from the TERMITe dataset.

#### Featurisation

To featurise our terminators for machine learning, we first separated our terminators into four parts: the A-tract, the stem, the loop, and the U-tract. We then zero-padded each element separately with ‘Zero’ bases such that each part had consistent length across the dataset and gave each base (or base pair for the stem) a numerical embedding. To ensure that each position in our feature vector corresponds to biologically similar bases across the dataset, we made the following embedding choices:

The bases of the **A-tract** are one-hot encoded (four binary integers, one for each base type) and right-aligned, such that bases closest to the stem align and zero-padding occurs furthest from the stem. Zero-padded bases are a vector of four zeros (Supplementary figure S1A).

The bases of the **stem** are encoded in pairs, starting at the base of the stem and moving up towards the loop. Each base pair is encoded as nine integers: eight one-hot-encoded integers to represent the bases in the upstream and downstream stem shoulders (in that order) and one binary integer to indicate if those bases are hydrogen bonded or not, determined by the predicted secondary structure given in the datasets. ‘Zero’ bases are used as pairing partners for bases in bulges, and to zero-pad the stem furthest from the base of the stem such that all stem embeddings in the dataset have the same feature vector length (Figure S1B).

The bases of the **loop** are one-hot encoded and middle-aligned, such that the middle base in the loop sequence is embedded at the center of each vector for each loop in the dataset. ‘Zero’ bases are therefore appended to the beginning and the end of each loop sequence to match their length to the longest loop in the dataset. Loops with an even number of bases, and thus have a two-base center, have an additional ‘Zero’ base inserted in the middle of the sequence Figure S1C).

The bases of the **U-tract** are one-hot encoded and left-aligned, such that zero-padding occurs furthest from the stem. For data from the TERMITe dataset (17), the point of termination was known and included as a Boolean feature (Figure S1D).

While the featurization choices for the A-tract and U-tract are intuitive, different choices could have been made to featurise stems and loops: the stem could have been embedded from the loop towards the base of the stem, and loops could have been encoded from the stem. Also, either of these alternative featurisation approaches could have been combined with the embedding we chose.

#### Random forest training

As termination efficiencies were determined with different methodologies for the Chen and TERMITe datasets, data may not be directly comparable. Also, the natural terminators from the Chen dataset likely shows significant overlap with the *E. coli* datapoints from the TERMITe dataset. Therefore, models were trained separately for each dataset. For the TERMITe dataset, we trained separate models for all *B. subtilis* terminators, all

*E. coli* terminators, and the full dataset, For the Chen dataset, we trained separate random forest models for the synthetic terminators, the natural terminators, and the full datasets.

Prior to training, data were split into test and cross-validation sets (test size: 0.5; number of cross-validation sets: 5), stratifying on termination efficiency. Stratification was done by assigning each terminator to one of five bins (termination efficiency = 0.0-0.2, 0.2-0.4, 0.4-0.6, 0.6-0.8, 0.8-1.0, upper bounds inclusive, 0.0 inclusive). These labels were then passed to the StratifiedShuffleSplit module for splitting the dataset into train and test sets, and the StratifiedKFold module for splitting the training set into cross-validation sets (scikit-learn 1.8.0). We used cross-validation datasets to select hyperparameters for training our random forest.

Random forests were trained using the RandomForestRegressor module from the scikit-learn library (v1.8.0; 1000 estimators, oob_score=True, default settings otherwise). Actual and predicted termination efficiencies were recorded for each datapoint in the test set and automatically plotted using our MEWTWO package.

We chose a large test set to rigorously analyse trends in the data (e.g. performance of the model on a range of termination efficiencies) and ensure that our data stratification strategies were appropriate, which was especially important as our data were highly biased: natural terminators displayed high TE on average (TE > 90% for most data points), while synthetic terminators took on extreme values (TE ∼ 0% and TE ∼ 100%; Figure ZA). However, this had negative impact on feature importance analysis, as feature importances were dependent on a randomly chosen subset of data points. For this reason, we decided to also train models on the full datasets purely for feature inference purposes (importantly: these models were not used to assess model performance or select parameters). While the most important features broadly correlated, there were some subtle but important differences (see feature importance tables on Zenodo). As models that have seen more data points can determine predictive features that are more broadly applicable, we decided that using the models trained on the full datasets for feature inference introduced the least bias. Feature importances were extracted and automatically visualised in hairpin representation, with more important features visualised as more saturated colours, using our MEWTWO package.

## RESULTS

### The 3’UTR spacer sequence upstream of the terminator hairpin affects protein production levels

The hairpin and poly(U)-tail of an intrinsic terminator play a clear role in transcription termination. However, the role of the bases between the STOP codon and the terminator hairpin have not been studied in detail. Therefore, this study sought to investigate the role of this ’3’UTR spacer sequence’. To this end, a completely randomized 30 bp 3’UTR spacer sequence library was generated and cloned between the STOP codon of a GFP or an RFP gene and the terminator hairpin of commonly used terminator BBa_B1002 (46) (Figure 1A). A spacer sequence shorter than 30 nucleotides can have a negative effect on the terminator’s termination efficiency, while increasing the size above 30 nucleotides does not seem to impact protein production (24). We transformed E coli with this plasmid library and measured expression levels for 56 transformants carrying RFP and a unique spacer sequence, which showed a 5.4-fold difference in fluorescence levels and 72 transformants carrying GFP and a unique spacer sequence with a 2.7-fold difference in fluorescence (Figure 1B, Supplementary data).

**Figure 1.**
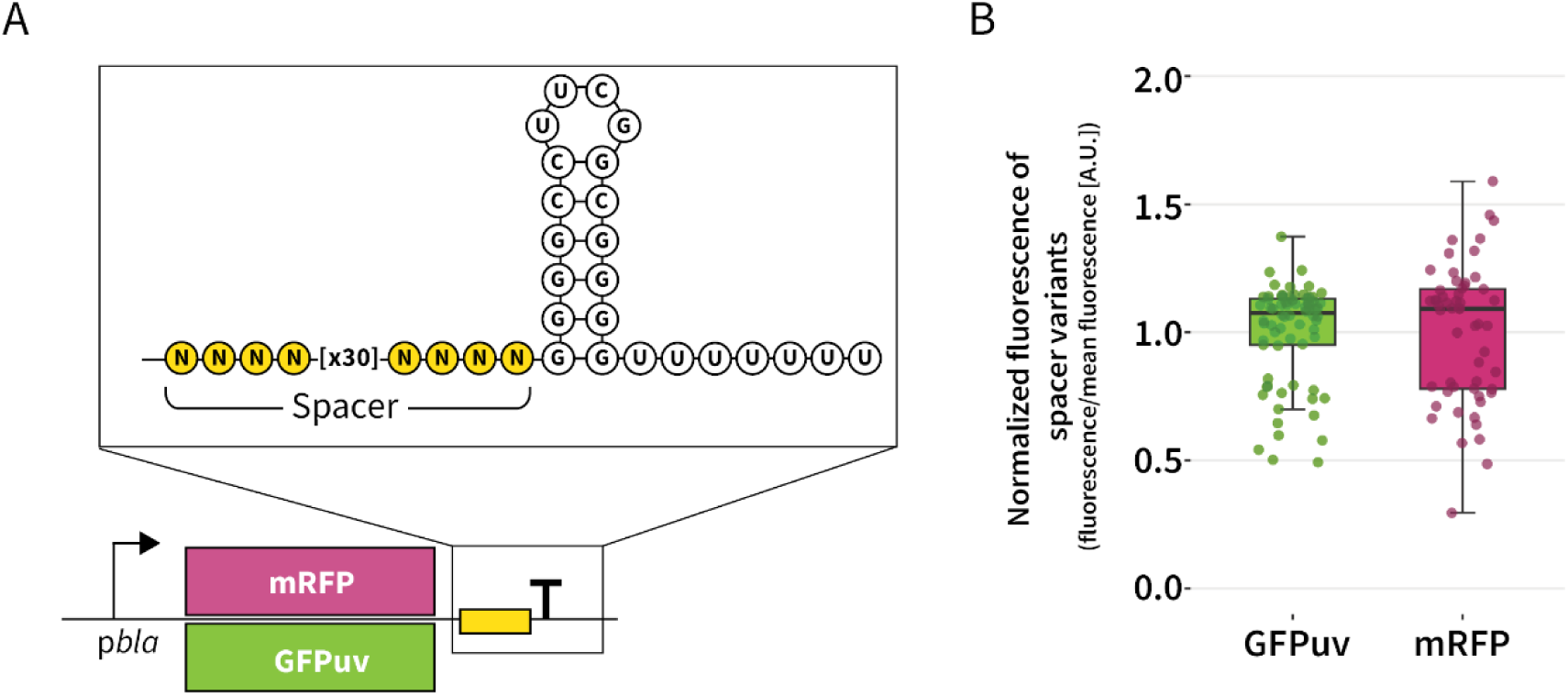
Generation and screening of a terminator library. (A) a completely randomized 30-bp sequence (yellow) was cloned between the hairpin of terminator BBa_B1002 and the coding sequences of an RFP and a GFP gene. The inset illustrates the resulting terminator RNA structure. (B) Flow cytometry analysis of unique transformants containing GFPuv + spacer (n = 72) and unique transformants containing mRFP + spacer (n = 56). The fluorescence of individual clones was normalized over the mean fluorescence of the respective population. Fluorescence of individual clones is shown as points, fluorescence distribution of the population is shown as a box-plot where the box represents the 25^th^ to the 75^th^ percentile, the line in the box represents the data median and whiskers represent minimum and maximum values for non-outlier data points that follow normal distribution.

### The effect of the 3’UTR on protein production appears mostly coding sequence independent

The difference in fluorescence resulting from expression of plasmids carrying the spacer library indicates that the 3’UTR spacer sequence influences protein production, and this effect appeared more pronounced for RFP than GFP. A previous study indicated termination efficiency can be influenced by the genetic context of the terminator sequence (26). Therefore, it was investigated whether a similar coding sequence context-specific effect also explains the observed effect of the 30-nucleotide spacer on protein production. To this end, ten 3’UTR spacer sequences were selected based on their respective resulting fluorescence levels covering a range of expression levels, five from the GFP library and five from the RFP library (supplementary data). These ten spacer sequences were cloned in the 3’UTR between an RFP gene, a GFP gene and a third reporter gene, the lacZ gene. After each spacer the same previously used terminator hairpin was added. The effect of the 3’UTR spacer sequence on protein production levels was measured by measuring the level of fluorescence for RFP and GFP and the level of substrate turnover for lacZ, and protein levels were normalized by the mean measured level of protein activity for the ten constructs for each reporter, to compare between the different conditions.

For all three genes, a similar trend was observed in the effect of the spacer sequence on protein activity, which was used a proxy for protein production levels (Figure 2A). The effect of the spacer on protein production was not exactly the same for all three proteins, the effect appears to be stronger for the RFP dataset compared to GFP and LacZ. Nevertheless, a strong positive correlation with an R^2^ of around 0.8 was observed between the relative GFP, RFP and LacZ levels (Figure S2), suggesting that the effect of the spacer sequence on protein production is hardly dependent on the coding sequence upstream of the spacer, but the effect appears mostly dependent on the spacer sequence itself. Interestingly, upon introduction of the same spacer and terminator sequences in the gram-negative *Pseudomonas putida* and gram-positive *Bacillus smithii*, a similar effect was observed, indicating that this termination feature appears conserved in multiple bacterial species (Figure S3).

**Figure 2.**
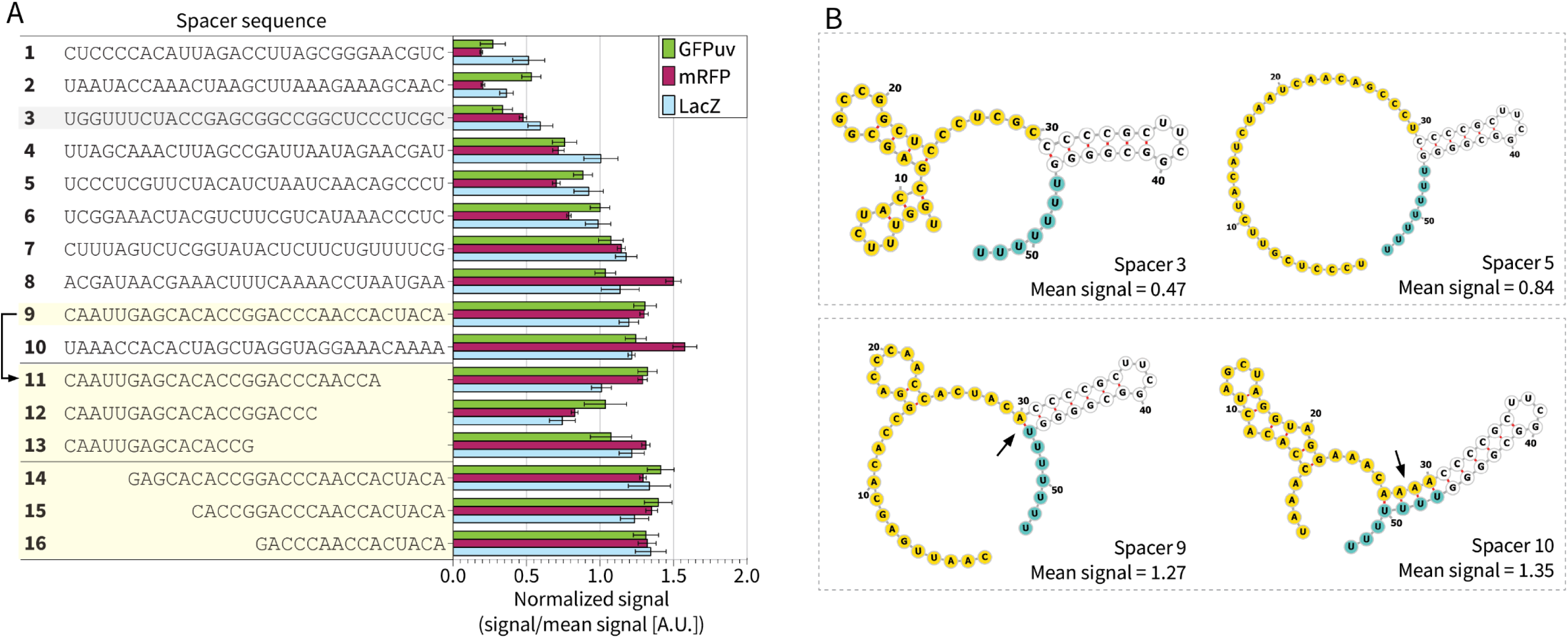
Analysis of the effect of 3’UTR spacer sequences on protein production. (A) Spacer sequences 1-10 were cloned behind GFPuv, mRFP and LacZ genes. Protein activity was used as a proxy for protein level. Protein activity of mRFP (magenta) and GFPuv (green) was determined by measuring fluorescence in a plate reader assay, LacZ activity (light blue) was determined by a Miller assay . To compare relative protein activity, the activity of each protein was divided by the mean activity over all spacers, resulting in a normalized signal. Spacer 9 (highlighted in yellow) was truncated from the 3’ end, resulting in spacers 10-13 and the 5’ end, resulting in spacers 14-16. Spacer 3, highlighted in grey, was selected as base sequences for further experiments (B) Secondary structures of 3’UTR regions containing selected spacers. The spacer is indicated in yellow, the poly(U)-tail is indicated in blue. Mean signal denotes the average normalized signal measured from GFP, RFP and LacZ analysis (panel A). The top box shows spacers 3 and 5 with a mean signal < 1, the bottom box shows structures of spacers 9 and 10, with a mean signal > 1. Arrows indicate predicted base pairing between the spacer and the poly(U)-tail. Predicted secondary structures of the remaining 3’UTR sequences are shown in Figure S4. mRNA secondary structures were visualized using Forna (45)

### Base pairing of hairpin-adjacent bases of the spacer and U-tract appears related to higher protein levels

To elucidate which part of the spacer sequence is responsible for the increased protein levels, one of the spacer sequences that lead to high expression, spacer 9, was truncated from both the 5’ and 3’ end. Truncation of the 5’ end up to 15 bases did not appear to have a significant effect on protein activity (Figure 2, spacers 14-16). When the spacer was truncated from the 3’ end by 5 or 15 bases, no general effect on protein levels was observed (Figure 2A, spacers 11 and 13). Only spacer 12, a 10-bp truncation ending in CCC, yielded lower activity of all reporter proteins. This indicated that not the length, but the base identity of the 3’ terminal bases of the spacer likely affected the level of protein production.

Interestingly, all spacers sequences associated with a normalized signal of > 1 ended in in either one or multiple adenine (A) or guanine (G) bases, which are all hypothetically able to base pair with the free uracil residues of the poly(U)-tail of the downstream terminator through A-U or G-U interactions. *In silico* prediction of the secondary structures of the 3’UTR sequences containing the different spacer sequences confirmed this. Modelling the mRNA secondary structures of all the of 3’UTR sequences that yielded a normalized signal of > 1 (spacers 6-11 and 13-16), predicted that the 3’ terminal bases of the spacer sequences base pair with the free uracil residues of the poly(U)-tail (Figure 2B, S4). 3’UTR sequences yielding a normalized protein signal of ≤ 1 (spacer sequences 2-6) appeared to have no strong interaction with the secondary structure of the terminator (Figure 2B, S4). One spacer sequence, spacer 1, completely disrupted the terminator secondary structure, due to the spacer being partially complementary to the terminator hairpin (Figure S4), also leading to the lowest protein levels (Figure 2A).

Through the predicted base pairing between the final base in the spacer sequence and the poly(U)-tail, two of the defining features of an intrinsic terminator may be altered: (i) the hairpin is potentially elongated by A/G-U base pairing in the bottom of the stem and (ii) the poly(U)-tail of the nascent mRNA is potentially shortened. Surprisingly, the number of A-U or G-U bases pairs does not necessarily seem to correlate with the increase in protein yield; spacer 9, with only a single predicted A-U bond leads to a higher protein signal compared to three A-U bonds for spacer 8.

### A single adenosine upstream of terminator G-C hairpin increases protein and mRNA levels

Previous experiments have showed that the hairpin of strong terminators is often preceded by an adenine-rich (A)-tract, which is often associated with bidirectionality of terminators and increased strength may indeed be attributed to elongation of the hairpin stem (15, 20). To understand if this structural change indeed contributes to increased protein production as we seem to observe in our results above, either by the lengthening of the hairpin or the shortening of free bases in the poly(U)-tail, a rational variant library was generated, where each of these features were systematically investigated.

The 3’UTR containing spacer 3 was used as the basis of the systematic variant library (Figure 3A). This spacer sequence was associated with a low corresponding protein level (protein/proteinMEAN ≤ 0.47, Figure 2A). From this spacer, three variant sets were generated, to systematically investigate which modifications in the stem or poly(U)-tail would lead to increased protein production, and whether the length of these modifications has an influence. In the first variant library, an (A)-tract of up to seven adenosine nucleotides was inserted at the end of the 3’UTR spacer sequence , which according to secondary structure predictions base pair with the poly(U)-tail, potentially leading elongation of the hairpin stem and reduction of free uracil residues in the poly(U)-tail (Figure 3B). Secondly, up to seven uracil nucleotides were added at the end of the 3’UTR spacer, while adding the same number of adenines downstream of the G-C hairpin (Figure 3C). This design ensures the folding of the hairpin has a similar predicted Gibbs free energy of the extended A-U hairpin part as the first set containing the (A)-tract (Figure 3B), while preventing base pairing between spacer and the poly(U)-tail. Thirdly, the number of free uracil bases was reduced by removing these from the DNA template, to investigate the effect of shortening the (U)-tail on protein production (Figure 3D).

**Figure 3.**
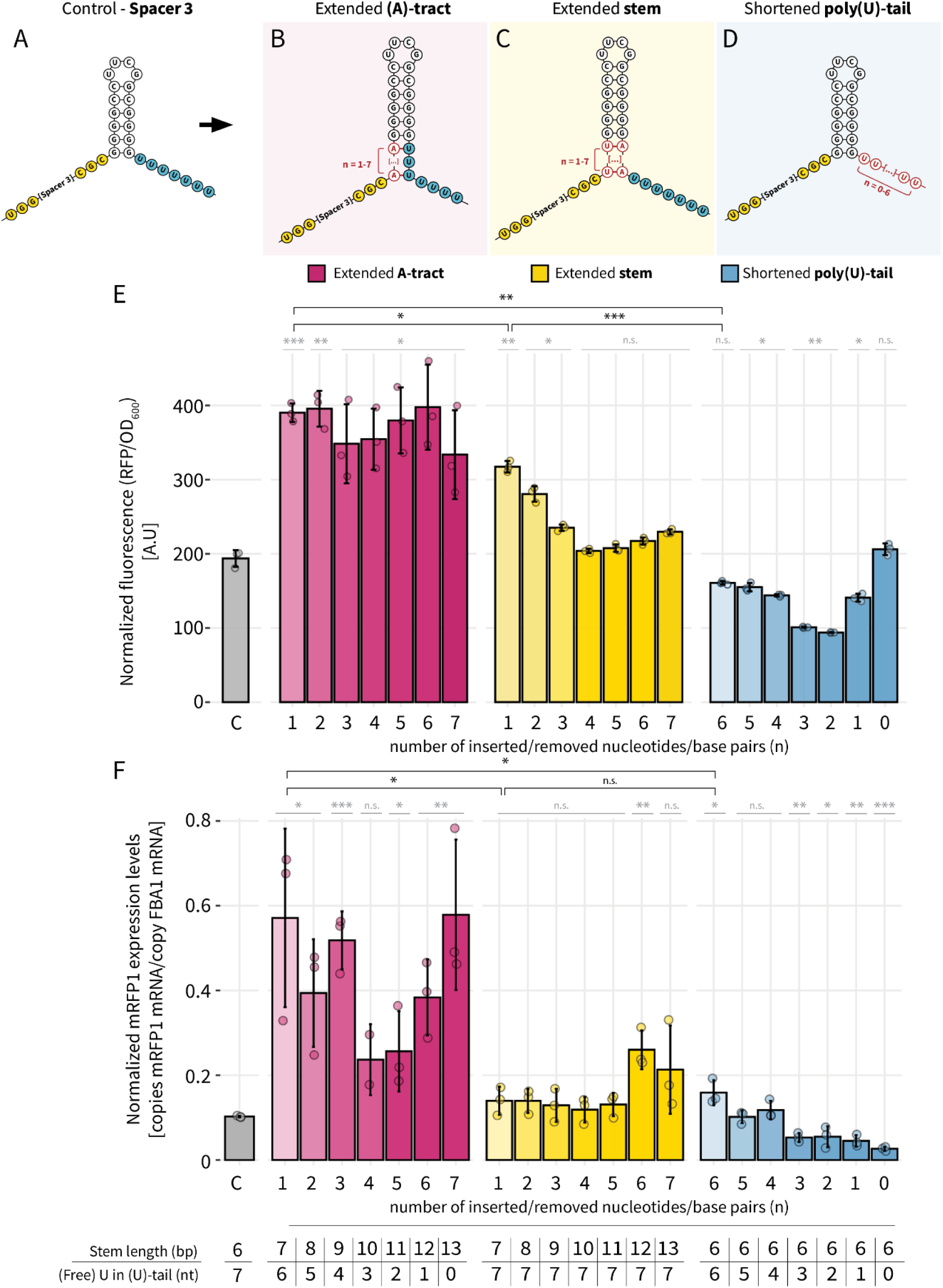
Systematic evaluation of the effect of base pairing within the 3’UTR. (A) from 3’UTR sequence containing spacer 3, several variants were made to assess their effect on protein levels. (B) The spacer was extended with an (A)-tract containing one up to seven adenosine nucleotides, resulting in a predicted base pairing between the (A)-tract and the poly(U)-tail. (C) The hairpin was extended by adding up to 7 U-A base pairs in the bottom of the hairpin stem. (D) The length of the poly(U)-tract was varied between zero to six uracil bases. (E) Average normalized fluorescence of *E. coli* cells expressing an RFP protein with the different downstream 3’UTR constructs. Bars represent average fluorescence of three biological replicate populations (circles) at the end of the exponential phase, measured by plate reader and normalized over OD600 of the cultures. (F) RFP mRNA copy number as determined by reverse-transcription digital droplet PCR (RT-ddPCR). Bars represent mean copy number of three biological replicates (circles) measured at the end of the exponential phase normalized over the copy number of glycolytic gene FBA1. For (E) and (F), the control was a 3’UTR with spacer #3 (grey, A) and was compared to 3’UTRs containing extended (A)-tract (pink, B), extended hairpin (yellow, C) and shortened poly(U)-tail (blue, D). The numbers on the y-axis indicate the number of added or removed bases, the length of the respective terminator hairpin stems, and poly(U)-tails are summarized in the table in the bottom of the figure. Biological variation is shown by error bars, and statistical significance between mean values is denoted with asterisks. Here, grey asterisks indicate the statistical significance of a sample compared to control C, and black asterisks show the statistical significance between specific samples, with the following significance levels: * p < 0.05, ** p < 0.005, *** p < 0.0005 and n.s., not significant.

All 22 variants were used as 3’UTR of an RFP coding sequence. When measuring fluorescence levels, it was observed that the insertion of a single adenine base at the end of the 3’UTR the spacer sequence , predicted to base pair with the first U in the poly(U)-tail, doubled the level of RFP fluorescence in the population, compared to the control spacer 3. Increasing the numbers of adenines in this position did not significantly improve protein levels further, although biological variation between the replicates increased with the number of adenines inserted (Figure 3E). Extending the hairpin alone, avoiding interactions between the spacer and poly(U)-tail, led to an increase of fluorescence compared to the original spacer 3 by approximately 1.5-fold, when a single U-A base pair was added in the bottom of the stem (Figure 3E). Fluorescence levels went down again as the stem length was increased. Similar data was observed when the hairpin stem was elongated with additional G-C base pairs, which yields more stable hairpins with a lower Gibbs free energy (Figure S5). Therefore, extension or increased stability of the hairpin alone did not explain the increase in protein level associated with base pairing of the spacer and the poly(U)-tail. Shortening the poly(U)-tail resulted in reduced levels of fluorescence. Since the poly(U)-stretch is required to initiate RNAP removal from the RNA-DNA hybrid during transcription, removing these likely leads to reduced termination efficiency. Surprisingly, when the poly(U)-tail was completely removed, fluorescence levels were similar to fluorescence of the control strain.

To investigate whether the effect occurred also at transcription level, the number of RFP mRNA copies was determined in each strain using RT-ddPCR. This showed that mRNA copy number significantly increased only for variants with predicted base pairing between the (A)-tract in the spacer and the poly(U)-tail (Figure 3F).

From this data, it can be concluded that the specific presence of a 3’UTR spacer ending with one or multiple adenosine nucleotides directly upstream of the terminator G-C hairpin improves protein up to two-fold and mRNA levels up to six-fold. This adenine is predicted to base pairing with a uracil base in the poly(U)-tail, leading to subsequent extension of the terminator hairpin stem. This base pairing hypothesis is enforced by the observation that insertion of a guanine base upstream of the hairpin, which can also base pair with uracil, leads to the same effect (Figure 2A). Nevertheless, extending the hairpin or shortening the poly(U)-tail alone did not explain this effect, highlighting the potential importance the specific interaction of this base with the poly(U)-tail.

### No clear effect of 3’UTR spacer sequence on mRNA stability

Experimental data showed that the presence of adenine residues upstream of the G-C hairpin have a positive effect on protein and mRNA levels, possibly due to predicted base pairing with the poly(U)-tail, There are multiple ways in which the mRNA level of a given gene could be influenced by the terminator secondary structure (47). On the one hand, increased termination efficiency has been associated with higher transcript levels, which for example by the result of increased RNAP available for re-inititation mediated by strong termination (21, 48). At the same time, increases in transcript levels could also be associated with reduced mRNA decay, as the poly(U)-tail serves as a toehold sequence for the RNA degradosome proteins poly(A)-polymerase, PNPase and RNAse III, while a strong hairpin protects an mRNA from 3’-end degradation by RNAses (25, 49).

It was investigated whether the increased mRNA levels of strains with adenosines at the end of the 3’ UTR spacer was caused by a reduction in turnover of the mRNA transcript, as a previous study indicated the 3’UTR is associated with mRNA stability (37). First, the terminator variants containing adenosine nucleotides at the spacer end and control spacer 3 without adenosines at the end (Figure 2A, B), were introduced in a series of *E. coli* strains harbouring single deletions in various proteins required for mRNA turnover. However, the RFP production between constructs with or without adenosines in the spacer, was not affected by removing components of the mRNA degradosome, indicating RNA decay may not play a role in the observed effect of the adenosine nucleotides upstream of the hairpin on fluorescence (Figure S6). Additionally, specific sequencing of mRNA 3’ ends (Term-seq (37, 38)) was used to determine mRNA decay and corresponding modifications of the mRNA 3’-ends. We hypothesized that reduced turnover would be visible as an increase in 3’end sequencing reads on the 3’UTR, while degradation or cleavage products would have high coverage on locations upstream of the 3’UTR (17). In addition, if the free poly(U)-tail was too short to serve as a toehold for RNAses, because of base pairing with the base pairing with the adenosine nucleotides, it will be polyadenylated by poly(A) polymerase (25). However, using TERM-seq, no polyadenylation of terminators was detected. Additionally, increased degradation products were only detected for strains with a shorter or no poly(U)-tail (Figure S7 and S8). In most instances, there was sequencing coverage of the terminators, but there was no clear difference in coverage of the terminators between the samples, therefore no strong conclusions could be drawn on the effect of the 3’UTR on mRNA stability.

### mRNA levels and fluorescence are increased by reduced terminator read-through

Alternatively, the increase in mRNA levels may also be attributed to increased transcription termination efficiency (TE). (21, 48). To verify if the predicted base pairing between the (A)-tract and the poly(U)-tail indeed affected transcription termination, a read-through experiment was performed. Here, a GFP coding sequence, including RBS but without promoter, was cloned behind the RFP+3’UTR variants with extended spacers with terminal adenosine nucleotides, extended hairpin and shortened poly(U)-tail (Figure 4A). Higher termination efficiency would be measured as reduced expression of the GFP gene. The control construct containing a 3’UTR with spacer 3 showed both RFP and GFP fluorescence, indicating read-through through the terminator and therefore a lower termination efficiency (Figure 4A). Insertion of adenosine nucleotides at the end of the 3’UTR spacer resulted in nearly a doubling of RFP fluorescence, consistent with previous results (Figure 2E), while the GFP fluorescence caused by read-through was decreased by 30 % compared to the control without adenosine nucleotides when only 1 adenosine was inserted, up to 60 % when 3 or 7 adenines were inserted (Figure 4A). When extending the hairpin with 1 or 3 U-A base pairs, red fluorescence increased slightly, by approximately 25 %, while GFP fluorescence was reduced slightly by 30 %, suggesting better TE, but substantially less improvement in TE was observed than for the A-U base pairs. In addition, when the stem was extended by 7 U-A base pairs, read-through dropped by 50% compared to the control. Shortening the poly(U)-tail alone did not significantly impact RFP fluorescence levels, while GFP fluorescence by read-through increased with shorter poly(U)-tails.

**Figure 4.**
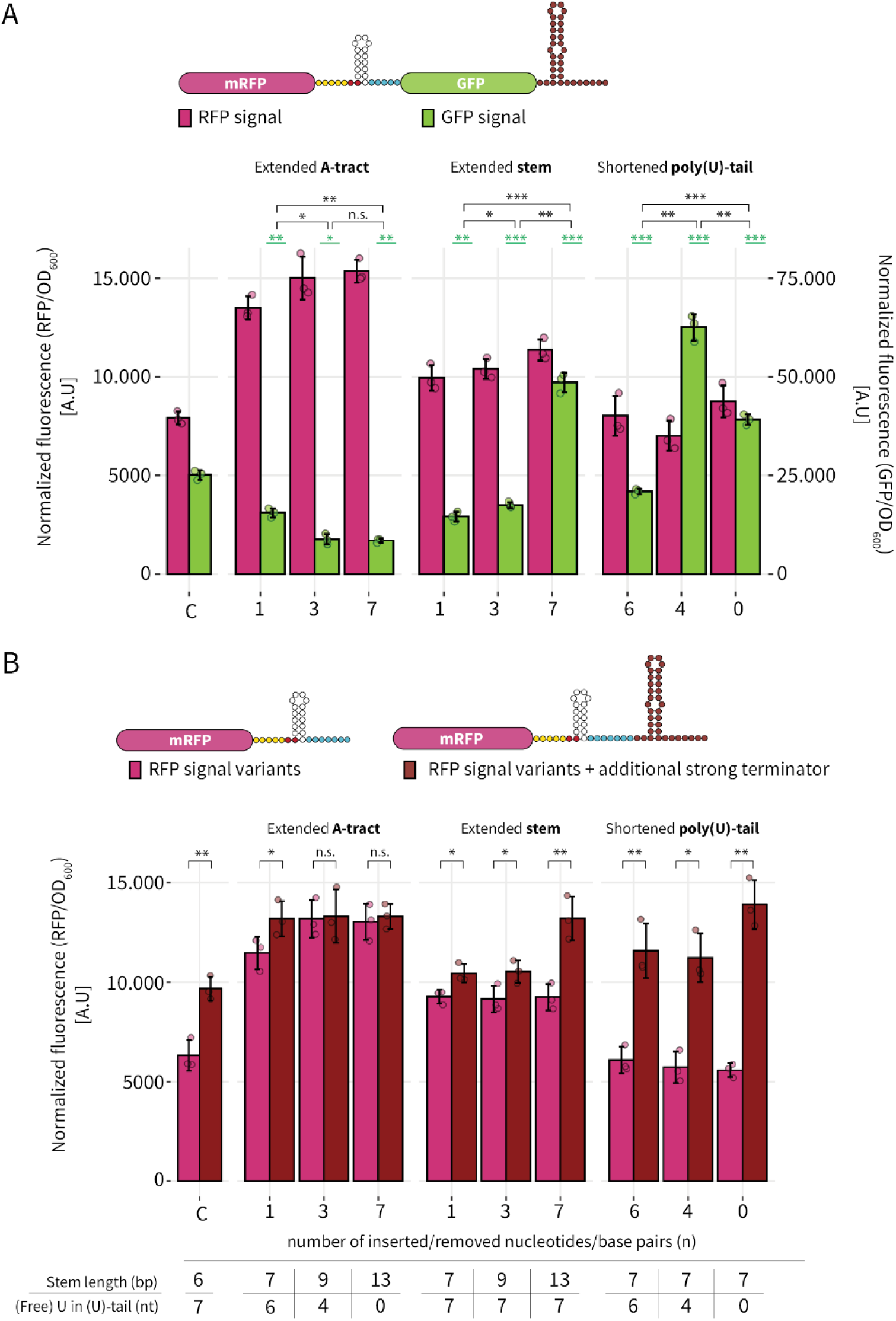
Termination efficiency of different 3’UTR variants. (A) GFP gene without promoter was cloned behind RFP genes with different 3’UTR variants (illustrated in Figure 3A-D) in order to assess read-through. Bars show average RFP fluorescence (magenta) and average GFP fluorescence (green) of three biological replicates (circles). (B) RFP fluorescence of an RFP gene with 3’UTRs illustrated in Figure 2A-D were compared to the same constructs with an additional terminator with high termination efficiency. Bars show average RFP fluorescence with the 3’UTR variants alone and (magenta) and average RFP fluorescence of constructs with a second terminator with high TE (dark red) of three biological replicates (circles). In both panels, fluorescence was measured by plate reader and normalized over OD600 of the cultures. The numbers on the y-axis indicate the number of added or removed bases, the length of the respective terminator hairpin stems and poly(U)-tails are summarized in the table in the bottom of the figure. Biological variation is shown by error bars, and statistical significance between mean values is denoted with asterisks. Here, green asterisks indicate the statistical significance of GFP fluorescence sample compared to green fluorescence in control spacer C, and black asterisks show the statistical significance between specific samples, with the following significance levels: * p < 0.05, ** p < 0.005, *** p < 0.0005 and n.s., not significant.

To further investigate whether the increase in RFP fluorescence is indeed a result of the increased termination efficiency, a second set of constructs was generated where a second terminator with a reported TE of close to 100 % (15) was cloned behind the 3’UTR sequences (Figure 4B), to ensure strong termination for all constructs. Here, all constructs showed a significant increase in fluorescence when the second strong terminator was added, except for the constructs containing the adenosine nucleotides in the spacer which already terminated efficiently (Figure 4B). This indicates that an increase in protein production is indeed associated with a high termination efficiency. Similar results were obtained for the 10 spacer variants originating from the random library, reinforcing the finding that increased protein production is associated with high termination efficiency, which is likely caused by base pairing between the (A)-ending spacer and the poly(U)-tail (Figure S9).

### Machine learning on wide set of terminators reveals that the identity of bases directly upstream of the G-C hairpin is a key feature for termination efficiency.

We next investigated if the identity of the bases directly upstream of the hairpin is predictive of termination strength in a wider range of natural and synthetic terminators. To assess this, we chose an explainable machine learning approach: we trained random forest regressors on natural and synthetic terminator sequences and their corresponding termination efficiencies, gathered from publicly available Term-Seq and reporter read-through data for *Bacillus subtilis* and *E. coli,* and subsequently performed feature inference to identify predictive sequence features. Critical for interpretability is a feature design method that ensures that each feature vector element consistently corresponds to the same terminator part in all sequences. To this purpose, we used mRNA secondary structure predictions to guide feature design (Figure S1; see methods). We were then able to encode each feature as a yes or no question in the following format: “Is the X^th^ base (pair) of the [terminator component] a [identity of the base]?”, e.g. “Is the 4^th^ base of the loop a C?” We trained six different random forest models on various data (sub)sets (Table 1) to study if predictive features depended on bacterial species or on whether the terminator was natural or not.

**Table 1.** Overview of datasets used for random forest training.

| Model name | Data collection method | Species of origin | Natural or synthetic | # datapoints | Data origin |
| --- | --- | --- | --- | --- | --- |
| **Term-Seq all** | Term-Seq | *E. coli* + *B. subtilis* | Natural | 1905 | Kosiński *et al.* (17) |
| **Term-Seq *E. coli*** | Term-Seq | *E. coli* | Natural | 691 |  |
| **Term-Seq *B. subtilis*** | Term-Seq | *B. subtilis* | Natural | 1214 |  |
| **Read-through all** | Read-through | *E. coli* | Natural + synthetic | 582 | Chen *et al.* (15) |
| **Read-through natural** | Read-through | *E. coli* | Natural | 317 |  |
| **Read-through synthetic** | Read-through | - | Synthetic | 265 |  |

While training a high-accuracy termination efficiency predictor was not our primary objective, we computed the Pearson and Spearman correlations from the actual and predicted termination efficiencies (TEs) for each data point in our test and cross-validation sets. The model trained on read-through data of synthetic terminators performed best, with a strong Pearson R of 0.90 between actual and predicted values (Figure 5A; Table S3). This is not surprising, as the synthetic terminators typically either terminate very strongly or not at all, turning a complex regression problem into a near-binary classification problem (Figure 5B). Models performed substantially worse on natural terminators (Pearson R = 0.34-0.48) but still showed weak to moderate correlation between actual and predicted TEs (Figure 5A; Figure 5C). Notably, the model trained on *E. coli* and *B. subtilis* Term-Seq data combined significantly outperformed models trained on terminators from either species individually (Figure 5A), suggesting that using sequence features that are important for both species may allow the model to generalise better.

**Figure 5.**
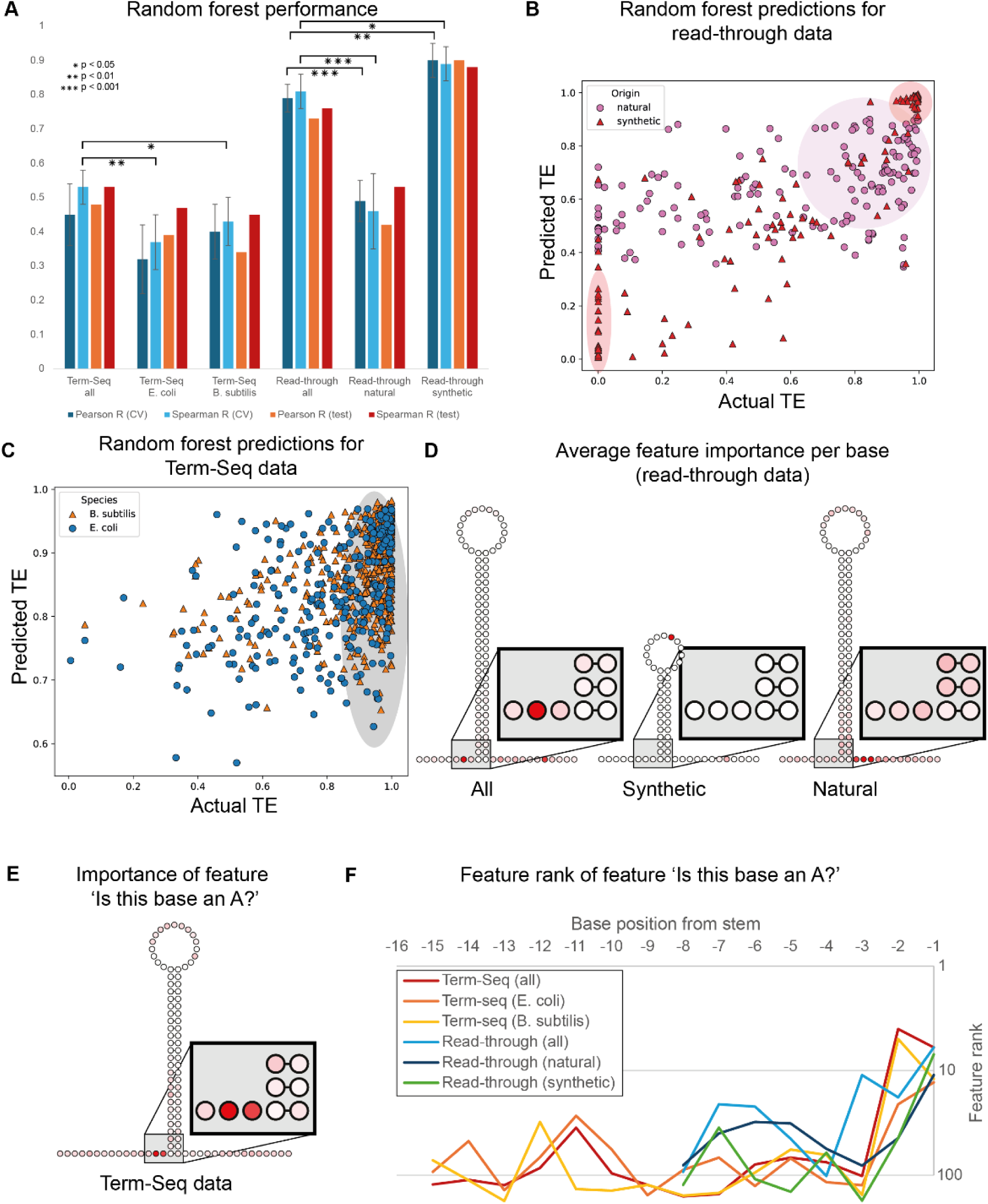
Feature inference reveals wide-spread importance of A bases in the spacer near the stem. (A) Pearson R and Spearman R of Plot showing actual termination efficiencies vs Random Forest predicted termination efficiencies for all six models. Error bars indicate standard deviation. CV: cross-validation. (B) Actual vs predicted termination efficiencies for a random forest classifier trained on synthetic and natural terminators from the read-through dataset (test set: 50% of the data). Pink: natural terminators. Red: synthetic terminators. Pink and red ovals show areas of high density in each dataset, showing high average termination efficiencies for natural terminators (pink), and extreme values for synthetic terminators (red). (C) Actual vs predicted termination efficiencies for a random forest classifier trained on *E. coli* and *B. subtilis* terminators from the Term-Seq dataset (test set: 50% of the data). The grey oval shows an area of high density in the dataset, showing high average termination efficiencies for both *E. coli* and *B. subtilis* terminators. (D) Average feature importance per base for three random forest regressors trained on all synthetic and natural terminators from the read-through dataset. The redder the base, the more important the features for that base. While the bases of the spacer proximal to the stem are unimportant relative to other bases in random forests trained on synthetic and natural terminators, individually, they harbour the most predictive bases in the model trained on both. (E) Feature importance per base of the feature ‘is this base an A’ for a random forest regressor trained on all *E. coli* and *B. subtilis* data from the Term-Seq dataset. The redder the base, the more important the feature. The two bases most proximal to the stem are shown to be the most predictive of termination efficiency. (F) Feature rank plotted against base position of the spacer. The six lines show the feature ranks of the feature ‘is this base an A?’ at the indicated positions for each of 6 random forest regressors. The X-axis displays the base position from the stem. The Y-axis displays the rank of the feature ‘Is the X^th^ base of the spacer an A’?

For feature inference, we proceeded with models trained on our train and test sets combined (see methods for our rationale). For models trained on natural terminators collected with Term-Seq data, the identity of the bases in the poly(U)-tail were most predictive of termination efficiency, with the most predictive feature in the model trained on both *E. coli* and *B. subtilis* terminators “Is the 3^rd^ base of the U-tract a U?”. This is not unexpected, as the poly(U)-tail is known to be an important component for intrinsic transcription termination. Features describing the poly(U)-tail were also most predictive for the models separately trained on *E. coli* Term-Seq data and *B. subtilis* Term-Seq data.

Notably, the feature “Is the 14^th^ base of the spacer an A” was the fourth most predictive feature in the model trained on all Term-Seq data (Table S4). This corresponds to the second base in the 3’UTR spacer upstream of the predicted G-C stem. This feature ranked highly for the models trained on *E. coli* and *B. subtilis* data separately as well (21^st^ and 5^th^ most important feature out of 437 features, respectively; Figure 5F). Upon inspecting all “Is the X^th^ base of [terminator component] an A” features, the most important were those mapping to the two bases directly upstream of the hairpin (Figure 5E). This is congruent with our experimental observation that the presence of A bases at the end of the 3’ spacers close to the G-C hairpin is predictive of termination efficiency and suggests that this applies broadly to intrinsic termination in both species.

For models trained on read-through data, most top-ranking features similarly pertained to the identity of bases in the U-tract. There were two notable exceptions: For the model trained on synthetic terminators only, the most important feature for this model was “Is the 8^th^ base of the loop a C?” and was 3.6 times as important than the next most important feature (“Is the 8^th^ base of the U-tract an A?”; Figure 5D, second panel; Table S4). This is most likely an artefact of the generation of the synthetic terminator datasets by Chen *et al*. Their dataset comprised three separate libraries, the first and second of which contained 192 weak to moderately weak terminators, while the third library contained mostly strong terminators. With the exception of 8 terminators with a pure poly(U)-tail and above-average termination strength, all terminators from libraries 1 and 2 contained a ‘C’ at position 8 of the loop, which was not the case for terminators from library 3. Therefore, we assume that this feature does not reflect biology, which is further supported by the fact that this feature (“Is the 10^th^ base of the loop a C?” f.i. = 0.008) is only ranked 20^th^ in the model trained on both natural and synthetic terminators from the read-through dataset. This bias in training data may also contribute to the high Pearson and Spearman correlations we observed for the model trained on synthetic terminators (Figure 5A; Table S3).

The most important feature for the model trained on read-through data of both synthetic and natural terminators was “Is the 7^th^ base of the spacer a U?”. This base, located two positions upstream of the stem, is the same base that was identified as important in the spacer for models trained on Term-Seq data, further indicating that the identity of this base is broadly predictive of termination efficiency. The “Is the 7^th^ base of the spacer an A?”, corresponding to the same base two positions upstream of the hairpin, was also highly ranked (19^th^ out of 409 features). For the model trained on natural terminators only, both features were less highly ranked (73 (U) and 44 (A) out of 409 features, respectively). As (by the authors’ design) almost all weak synthetic terminators contained a ‘U’ at this position, while almost all strong synthetic terminators contained an ‘A’, data bias almost certainly contributes to the importance of this feature. However, as natural terminators made up a large proportion of the training set for this model, biological signal likely also contributes to the importance of this base.

We also obtained per-base(pair) importances by summing all importances for features that ask a question about that specific base(pair). For the full read-through dataset (natural + synthetic), the base two positions upstream of the stem, as also identified for the Term-Seq dataset, was the most predictive of TE. In contrast, this base was much less important for models trained independently on natural or synthetic terminators, respectively (Figure 5D). This may indicate that the model needs this feature to generalise better across a range of terminators, hinting that the identity of this base is a feature that is generally predictive of termination efficiency, regardless of whether the terminator is of synthetic or natural origin.

Finally, we demonstrated the importance of the bases proximal to the stem in the spacer by comparing the feature rank of each “Is the X^th^ base of the spacer an A” feature against its position: for all six models, the feature rank peaks in one of the two bases directly upstream of the stem, which is not the case for other bases at these positions (Figure 5F; Figure S10). This further supports our experimental findings that A bases directly upstream of the terminator hairpin are important for intrinsic termination.

### NusA and NusG do not significantly influence *in vitro* termination efficiency

As the occurrence of the adenosine nucleotides upstream of the hairpin in natural terminators with high TE appears an important feature in termination efficiency *E. coli* and *B. subtilis*, and the strength of these terminators cannot be explained by extension of the hairpin alone, we questioned whether there are additional proteins that play a role in intrinsic termination that may interact with the adenines upstream of the hairpin and improve termination efficiency.

Literature research on proteins described to be associated with intrinsic terminators resulted in two candidates: NusA and NusG. The role of NusA and NusG has been studied in the context of intrinsic termination and their interaction *in vivo* and *in vitro* with A-U base have been hypothesized before (31). Both proteins are part of the transcription complex, where NusA is already known to be associated with transcriptional pausing, rho-dependent termination and phage λ N-mediated antitermination (50), while NusG is demonstrated to be involved as an anti-pausing factor, coupling of transcription and translation and rho-dependent termination (51, 52). Additionally, several reports indicate that NusA is required for transcription termination *in vitro* of specific *E. coli* terminators, including the *rrnB* and the *trp* intrinsic terminator (28, 53), and in *Bacillus subtilis* NusA and NusG are essential for termination of intrinsic terminators that contain an A-U or G-U base pair in the bottom of the terminator stem (27, 54). Combined with our experimental findings, this suggests that also in *E. coli*, NusA, and potentially NusG, may improve termination efficiency of intrinsic terminators, especially terminators with an A-U or G-U bond in the bottom of the terminator hairpin stem.

This hypothesis was investigated in an *in vitro* assay, as we cannot study this in vivo as NusA and NusG are essential proteins and deletion results in loss of viability (32). By measuring the *in vitro* read-through rate of purified *E. coli* RNAP in presence or absence of either NusA and/or NusG, the effect of these proteins could be determined. Read-through was measured at transcriptional level by cloning a fluorescent Mango-III aptamer behind the different 3’UTR variants, allowing for real time monitoring of transcriptional read-through (Figure 6A). Spacer 3 was taken as a reference and compared to read-through of the (A)-extended spacer, the extended hairpin and the shortened poly(U)-tail. A construct with a Mango-III aptamer without any upstream terminator was used as positive control for Mango-III signal in presence and absence of NusA and NusG.

**Figure 6.**
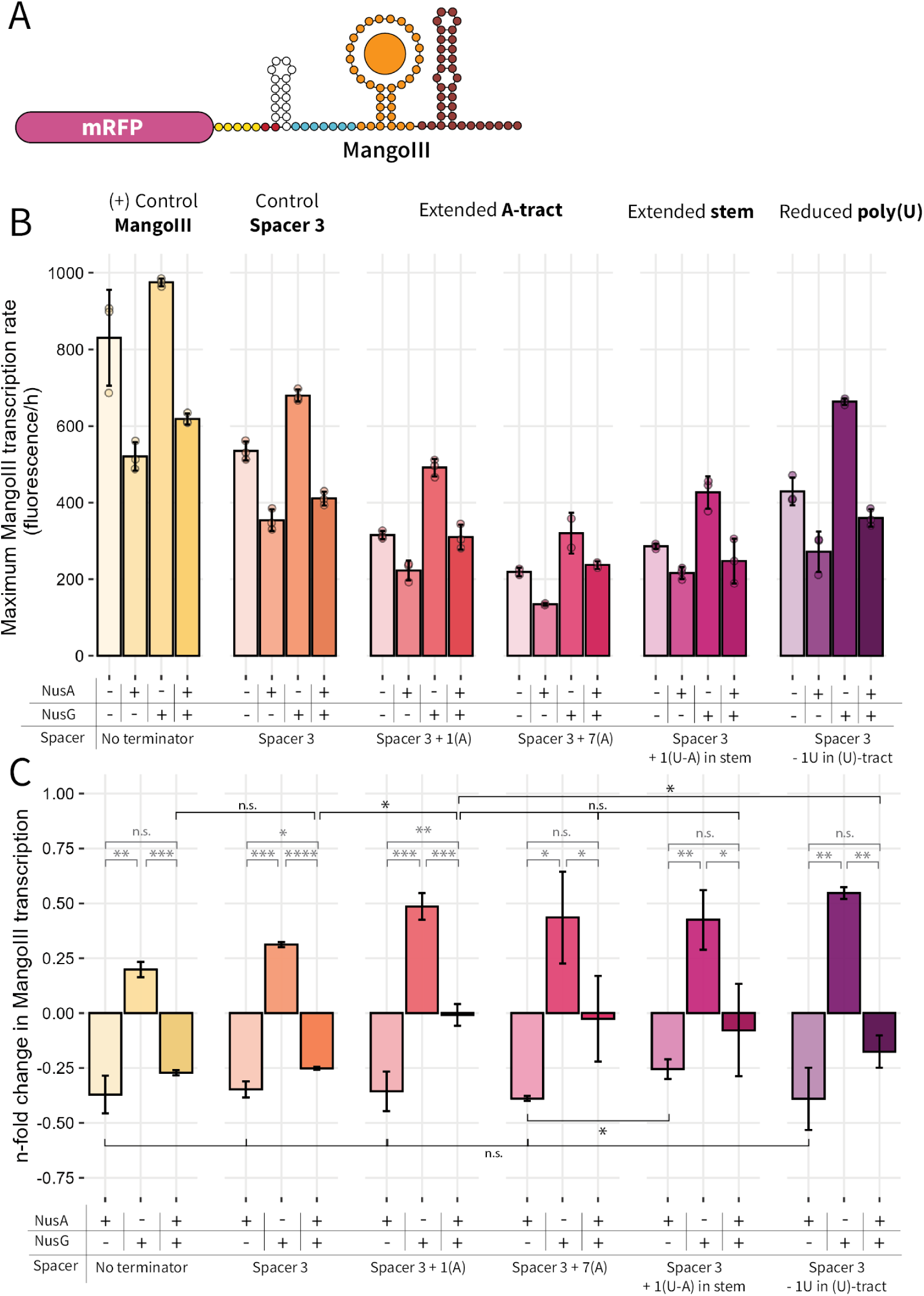
**in vitro assay to test the effect of NusA and NusG on different terminator designs**. (A) Schematic of the mRNA construct generated in the *in vitro* assay. A linear DNA fragment containing a *bla* promoter was used to express an RFP mRNA with different variants of 3’UTRs (colored circles). A Mango-III aptamer, able to bind fluorophore TO1-3PEG-biotin, was cloned directly behind the 3’UTR variants, and would only be transcribed upon terminator read-through. Another strong terminator was cloned behind the Mango-III aptamer. The 3’UTR upstream of the Mango-III aptamer was omitted in the (+) control. (B) maximum Mango-III transcription rates in in *in vitro* transcription assay in presence (+) or absence (-) of NusA and/or NusG. Bars represent the average maximum rate of fluorescence production measured by plate reader of three technical replicates (circles). The 3’UTR variants correspond to the variants illustrated in figure 3A-D. (C) Fold-change effect of addition of NusA and or NusG in on Mango-III transcription rates versus the condition where no Nus-proteins were added. N-fold change is defined of the maximum transcription rate of a condition divided by the maximum transcription rate measured when neither NusA nor NusG were added. Bars represent the mean n-fold change of three technical replicates. Technical variation is shown by error bars, and statistical significance between mean values is denoted with asterisks. Here, grey asterisks indicate the statistical significance between conditions of a single 3’UTR construct while black asterisks show the statistical significance between specific samples, with the following significance levels: * p < 0.05, ** p < 0.005, *** p < 0.0005, **** p < 0.00005 and n.s., not significant.

When looking at absolute Mango-III transcription levels, in all samples a measurable level of Mango-III signal was detected, meaning that *in vitro* none of the terminators had a 100% termination efficiency, regardless of the presence of NusA or NusG. Generally, read-through was highest for the 3’UTR containing spacer 3 with no further modifications, and for the 3’UTR with spacer 3 including a shortened poly(U)-tail (Figure 6B, Figure S11), which is consistent with the *in vivo* read-through experiments (Figure 5A). The lowest read-through was observed for the 3’UTR containing seven A bases in the extended spacer. The 3’UTR sequences containing a single adenine base and the 3’UTR with an elongated stem behaved similar, with approximately 50 % read-through compared to the control, which is again consistent with the *in vivo* read-through measurements.

In all samples, also in the positive control without terminator, a clear effect of the addition of NusA and NusG was observed: NusA generally reduced the Mango-III transcription rate, and therefore probably had a positive effect on termination efficiency, while addition of NusG increased Mango-III transcription and therefore had a presumed negative effect on termination efficiency. NusA is a pausing and termination factor and has been described to lower transcription rates (55). Thus, it is unsurprising that addition of NusA assists reduces transcription rates, while the role of NusG as an anti-pausing factor can promote read-through. Nevertheless, it should be noted that with this assay, it is impossible to confidently distinguish between improved termination efficiency or reduced transcription, which can both cause a decrease in Mango-III transcription rate. The RNA yield of each reaction was analyzed by separation on an agarose gel but no significant differences in RNA-yield were detected (Figure S12).

To investigate if the function of NusA or NusG is directly linked to the occurrence of the adenosine nucleotides upstream of the terminator hairpin, the relative effect of NusA and/or NusG on transcription rate was determined. This was done by calculating the fold-change effect of adding one or both proteins compared to the condition where no proteins are added (Figure 6C). Sole addition of NusA did overall decrease transcription read-through, but with no clear differences in fold-change across different terminator design, only for the hairpin with a U-A pair at the bottom this fold-change was less strong than all the others. So, we could not detect a specific improvement of termination efficiency by a combination of NusA and the A-U base pairing at the bottom of the hairpin specifically, as the fold-change effect on e.g. the control spacer 3 without an A at the spacer end was similar.

The addition of NusG generally led to a significant increase in Mango-III transcription and therefore decreased termination efficiency between the control spacer and the extended (A)-base spacers. However, this increase was also visible for the 3’UTR with an extended stem or shortened poly(U)-tail and is therefore not exclusively linked to the presence of the adenosines.

The combined addition of both NusA and NusG led to a reduced transcription rate of Mango-III when using control spacer 3 and the 3’UTR with shortened poly(U)-tail. These reduced transcription rates were not significantly different from the transcription rate of the positive control (Mango-III without terminator). For all other spacers, including the A-ending spacers, the addition of both proteins appeared neutral, meaning the Mango-III transcription levels of these variants were similar to the condition where no proteins were added. The difference in effect between the A-containing spacers and control spacer 3 may suggest a biological difference. However, there is no significant difference between transcription rates of the 3’UTR with adenosine nucleotides upstream of the hairpin and the 3’UTR with an extended stem, while *in vivo* transcription and read-through rates clearly differed between these samples (Figure 2E, Figure 4A). Therefore, these *in vitro* results may not fully represent the observations done *in vivo*. Additionally, it is difficult to distinguish reduction in transcription termination and transcription rate, as both would lead to a reduction in fluorescence of the Mango-III aptamer. Since similar effects were observed for the positive control, it is possible that the detected changes were due to changes in transcription rate rather than termination efficiency.

## DISCUSSION

This study presents evidence that the inclusion of a number of adenosine nucleotides upstream of the hairpin of an intrinsic terminator increases termination efficiency, resulting in an increased level of mRNA and consequently, a higher protein production rate.

Although the (A)-tract is frequently included in intrinsic terminators, its biological function is mostly attributed to bidirectionality of the terminator. Nevertheless, the relation between higher TE and the presence and potential base pairing of an (A)-tract and the terminator poly(U)-tail has been observed before, and was hypothesized be caused by a more stable terminator hairpin . While the necessity of the (A)-tract was questioned before, the A-tract’ t and (U)-tail have often been considered the key factors in termination efficiency. In the present study, we demonstrated that the presence of one or more adenosine nucleotides upstream of the terminator are linked to higher termination efficiency. However, besides bidirectionality, which was not assessed in the current study, the length of the (A)-tract may be as short as one or two adenosines. We did not find evidence that a longer (A)-tract resulted in stronger gene expression or strong termination. Moreover, the increase in TE is not related to extension of the hairpin alone, as extension of the stem-loop structure by the much stronger G-C base pair or by an inverted U-A base pair did not yield the same results. Extension of the hairpin stem through other interventions did reduce the termination efficiency, indicating the insertion of adenosine nucleotides upstream alone improved TE. These findings highlight the importance of the nucleotide identity of these specific bases upstream of the hairpin, being an adenine, or possibly a guanine, rather than their effect on the secondary structure of the terminator. Concluding, our study indicates neither an extended (A)-tract, nor a longer hairpin, are required for strong intrinsic termination efficiency in the sense-strand direction, but that at least one adenosine nucleotides may be sufficient to increase termination efficiency.

The importance of the base identity combined with its position and feature in the 3’UTR was corroborated by our machine learning model. Especially when our models were trained on mixed data (natural + synthetic terminators, or *B. subtilis* + *E. coli* terminators) and therefore were forced to generalize, the importance of an adenine upstream of the terminator hairpin became more important, highlighting this could be a feature generally associated with strong termination in bacteria.

The observation that specifically A (or G) bases at the start of the hairpin lead to increased TE, suggest a mechanistical explanation, as only these bases allow for base-pairing with the downstream uracil bases in the poly(U)-tail. A hypothetical mechanism could be that, once the RNAP elongation complex pauses on the poly(A)-tract, including at least the first uracil base of the nascent poly(U)-tail in the hairpin formation may aid in hypertranslocation of the RNAP from the RNA-DNA duplex, or the helps in more efficient hybrid shearing of the RNA-DNA duplex (11). When the hairpin is simply extended, but does not specifically base pair with the poly(U)-tail (which resides in the RNAP core tunnel), this effect may be less strong, explaining why termination efficiency is improved with and A (or G)-ending spacer, rather than extended hairpins containing e.g. U-A or G-C in the base of the stem.

Alternatively, previous studies, especially in *B. subtilis*, have suggested that weak terminator hairpins with A-U or G-U base pair at their base. require both NusG and NusA to efficiently terminate (27, 54). We tested the effects of the *E. coli* NusA and NusG proteins *in vitro*, but could not demonstrate a specific effect of these proteins on the termination efficiency of the G-C hairpins preceded by adenines described in this study. However, we were only able to test this in *in vitro* conditions, with concentrations not representative of *in vivo* conditions. Further detailed mechanistic studies are needed to more clearly reveal effects of these or other factors to further explain the clear role of adenosine nucleotides before the G-C hairpin.

Other than revealing the relevance of these bases to further improve our understanding on bacterial intrinsic termination, this study also demonstrates that considering the base identity at the end of the ‘3-UTR spacer/bottom of the hairpin is an important, and up until now, often overlooked factor for synthetic gene design for heterologous gene expression and synthetic genome and circuit design. Including at least one adenine at this position seems a robust strategy to improve gene expression levels and generate strong transcription terminators, which, as shown in this study, holds for various genes in *E. coli* as well as in some other bacteria. Further synthetic terminator design algorithms, which are currently emerging (e.g. (20)) should take this feature into account to improve their termination efficiency. By using 3’UTR spacer sequences containing an adenosine upstream of the hairpin, transcriptional read-through is reduced and transcription and protein levels can be boosted, providing a simple and useful insight when tuning transcription or translation of a (heterologous) gene.

## ACKNOWLEDGEMENTS

The authors thank Hilke Schenkel for his guidance in the *in vitro* transcription assay, Jaco van der Torre for providing expression vectors for GreA, GreB and NusG, and Thijmen Zegers for assistance in experimental work. This publication is part of the project “Evolving life from non-life (EVOLF)” of the research programme SUMMIT, which is financed by the Dutch Research Council (NWO) and “BaSyC – Building a Synthetic Cell” Gravitation grant of the Netherlands Ministry of Education, Culture and Science (OCW) and the Netherlands Organisation for Scientific Research (NWO).

## AUTHOR CONTRIBUTIONS

Charlotte C. Koster: Conceptualization, Data curation, Formal analysis, Investigation, Project Administration, Resources, Validation, Visualization, Writing – original draft. Barbara Terlouw: Conceptualization, Data curation, Formal analysis, Investigation, Software, Validation, Visualization, Writing – original draft. Thijs Nieuwkoop & Sjoerd Creutzburg: Conceptualization, Formal analysis, Investigation, Validation, Visualization. Maria Martin-Pascual, Miguel Paredes-Barrada, Panagiotis Kopfsiaftis, Hans Heilig & Theo van Laar: Investigation, Resources. John van der Oost & Nico J. Claassens: Conceptualization, Project Administration, Supervision, Writing – review & editing.

## DATA AVAILABILITY

The data underlying all figures this article are available in the article and in its online supplementary material. All code needed to replicate data parsing, featurization, model training, and feature inference can be found in the MEWTWO (mRNA Expression Wizard 2) repository at Zenodo, via doi: 10.5281/zenodo.20714791; and GitHub: https://github.com/BTheDragonMaster/mewtwo). Raw sequencing data are available in the Bioprojects/Sequence Read Archive (SRA) at https://www.ncbi.nlm.nih.gov/bioproject/ and can be accessed with BioProject ID PRJNA1472557. The Galaxy workflow and .fasta files used for alignment can be accessed via Zenodo at doi: 10.5281/zenodo.20717389.

## APPENDIX

Supplementary files accompanying the manuscript:

### Supplementary data and methods (supplementary_data_methods.pdf)

Contents:

#### Supplementary methods

Term-seq RNA sequencing

Affinity chromatography of proteins

### Supplementary figures S1-S12

Figure S1. Embedding strategy for terminators.

Figure S2. Pearson correlation coefficients between constructs GFPuv, mRFP and LacZ activity containing spacers 1-16 in the 3’UTR.

Figure S3 Analysis of the effect of 3’UTR spacer sequences on protein production in *Pseudomonas putida*.

Figure S4. Secondary structures of 3’UTRs with spacer variants 1-16.

Figure S5. Extending the terminator hairpin by A-U and G-C base pairing.

Figure S6. Effect of genomic deletion of inessential mRNA turnover machinery on protein production of 3’UTRs containing an (A)-tract.

Figure S7. Per-base coverage of a Termseq dataset of plasmids containing all 3’UTR rational variants.

Figure S8. Per-base coverage of a Termseq dataset of plasmids harboring 3’UTR spacers 1-10. Figure S9. Termination efficiency of different 3’UTR containing spacers 1-10.

Figure S10. Feature rank plotted against base position of the spacer.

Figure S11. Mango-III fluorescence as a proxy for transcription over time in an *in vitro* transcription assay in presence or absence of NusA and/or NusG. Figure S11. Transcription products after *in vitro* transcription assay.

Figure S12. Transcription products after *in vitro* transcription assay.

Table S1. 3’UTR sequences used in this study. Table S2. ssDNA oligos used in this study

Table S3: Mean Pearson R and Spearman R on cross-validation and test sets. Table S4: Top 10 features per model (trained on the full dataset).

*DNA sequences used in this study*

### Supporting data (supporting_data.xlsx)

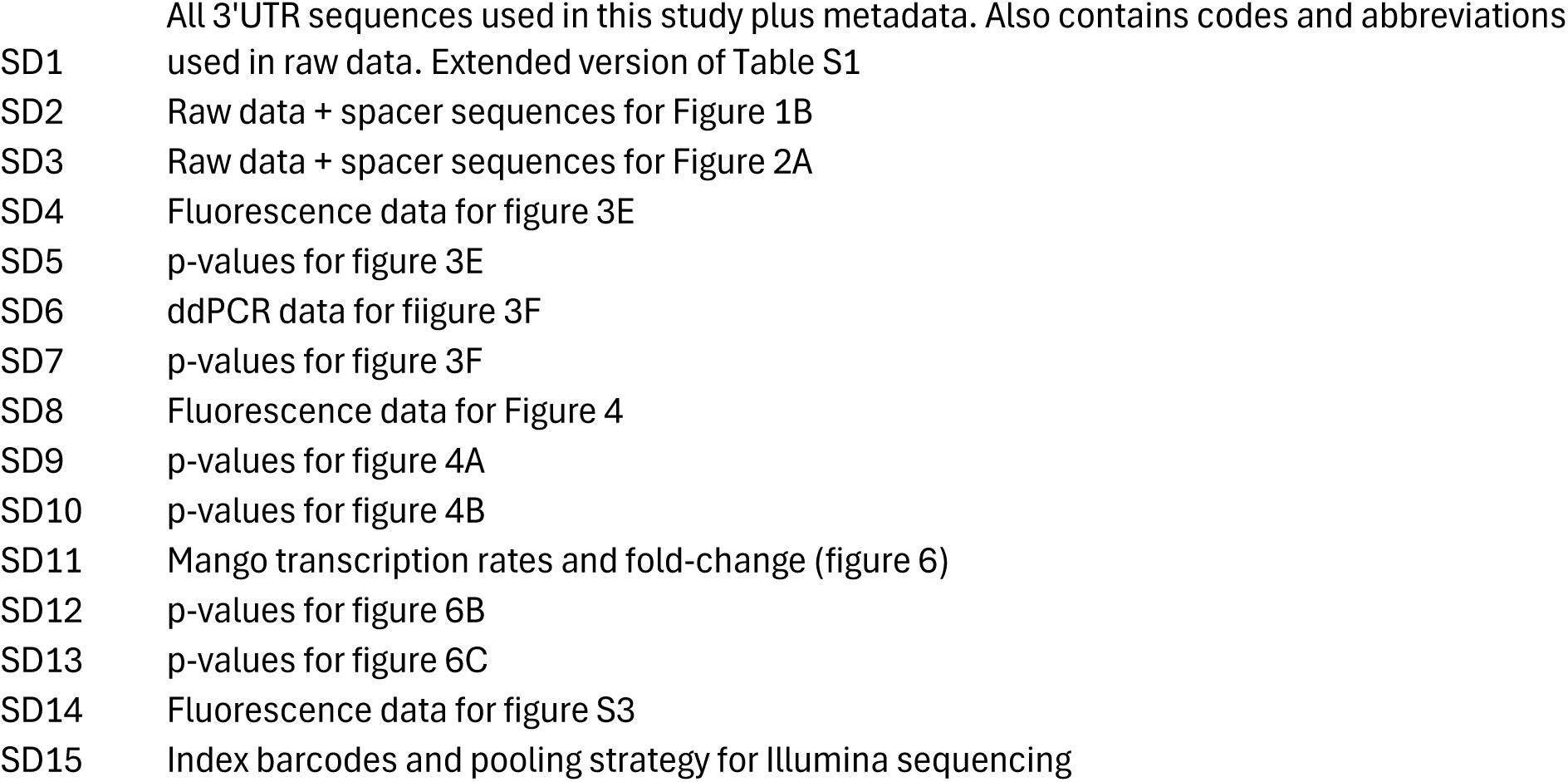

