## Supplementary data and methods for "Systematic engineering and machine learning analysis of intrinsic terminators reveal crucial nucleotides directly upstream of the terminator hairpin"

#### Contents:

##### Supplementary methods

Term-seq RNA sequencing

Affinity chromatography of proteins

##### Supplementary figures S1-S11

Figure S1. Embedding strategy for terminators.

Figure S2. Pearson correlation coefficients between constructs  GFPuv, mRFP and LacZ activity containing spacers 1-16 in the 3'UTR.

Figure S3 Analysis of the effect of 3'UTR spacer sequences on protein production in *Pseudomonas putida*.

Figure S4. Secondary structures of 3'UTRs with spacer variants 1-16.

Figure S5. Extending the terminator hairpin by A-U and G-C base pairing.

Figure S6. Effect of genomic deletion of inessential mRNA turnover machinery on protein production of 3'UTRs containing an (A)-tract.

Figure S7. Per-base coverage of a TERM-seq dataset of plasmids containing all 3'UTR rational variants.

Figure S8. Per-base coverage of a TERM-seq dataset of plasmids harboring 3'UTR spacers 1-10.

Figure S9. Termination efficiency of different 3'UTR containing spacers 1-10.

Figure S10. Feature rank plotted against base position of the spacer.

Figure S11. Mango-III fluorescence as a proxy for transcription over time in an *in vitro* transcription assay in presence or absence of NusA and/or NusG. Figure S11. Transcription products after *in vitro* transcription assay.

Figure S12. Transcription products after *in vitro* transcription assay.

#### Supplementary tables S1-S4

Table S1. 3’UTR sequences used in this study.

Table S2. ssDNA oligos used in this study

Table S3: Mean Pearson R and Spearman R on cross-validation and test sets.

Table S4: Top 10 features per model (trained on the full dataset).

#### DNA sequences used in this study

Backbone

Insert sequences

- GFPuv
- RFP
- LacZ

Readthrough parts

- eGFP + T0t
- L3S2P52 terminator

Mango-III(10AU) constructs

### Supplementary methods

#### Term-seq RNA sequencing

A Term-seq protocol was based on protocols by Dar *et al.* and Choe *et al.* (20, 34). RNA was isolated from three biological replicates as described above. All samples were equilibrated to 0.5 µg/µL. All materials were cleaned with RNAseZap (Thermo Fisher Scientific, AM9780). All steps were performed in RNase-free consumables and buffers, on ice, unless indicated otherwise.

*3' end ligation*. Nine custom barcoded i5 DNA adapters for multiplexing compatible with Illumina Nextera Next-generation sequencing kits (Table S2) were ordered as HPLC-purified ssDNA oligos with 5' and 3' phosphorylated ends (Sigma-Aldrich). Adapters were ligated to the 3' end of total RNA. Replicates of the same strains were separated in three groups where each RNA sample within a group was ligated to one of nine unique i5 adapters, which to allows for pooling at a later stage (Table S2, Supplementary data, SD_15). 2.5 µg RNA of each sample was ligated to their respective i5 adapter using T4 ssRNA ligase 1 (NEB, M02040L) in a ligation reaction containing 150 µM i5 3' end adapter solution, 1X T4 RNA ligase I buffer, 1 mM ATP, 8 % DMSO, 20 % PEG8000, 1U T4 RNA ligase I, incubated at RT for 2.5h.

*Magnetic bead purification*. The ligation reaction was cleaned up using Mag-Bind® TotalPure NGS magnetic beads (Omega Bio-tek, M1378-00), by adding a 1.2X volume of beads and separating these on a magnetic stand. The supernatant was removed and beads were washed twice with 120 µL 70 % ethanol.  Beads were air-dried for 5 minutes to remove ethanol traces, and RNA was eluted in 11 µL of nuclease-free water. Beads were removed on a magnetic stand and the supernatant was transferred to a fresh tube.

*rRNA removal*. Samples were pooled in twelve different pools, as such that each pool did not contain multiple biological replicates or multiple of the same i5 DNA adapter barcodes (Table S2, Supplementary data, SD_15). 11 µL of each pool was used for rRNA depletion. rRNA was depleted using a NEBNext rRNA depletion kit for bacteria (NEB, E7850S) according to manufacturer's protocol. rRNA depleted RNA was eluted in 7 µL of nuclease-free water.

*Fragmentation*. 5 µL of rRNA depleted RNA was fragmented using NEBNext Magnesium RNA Fragmentation Module (NEB, E6150S). 5 µL of rRNA depleted RNA was added to 15 µL 1X RNA fragmentation buffer. The fragmentation reaction was placed in a pre-heated thermocycler at 95 °C for 1.5 minute, then directly placed on ice and stopped with stop solution.  Fragmented RNA was was purified with magnetic beads as described above, but with a 2.2X volume of magnetic beads and eluted in 7 µL of nuclease-free water.

*First strand synthesis*. First strand cDNA synthesis was done using Protoscript II First Strand cDNA Synthesis Kit (NEB, E6560S) using primer BG34795, specific for the i5 adapter ligated to the total RNA. First strand cDNA synthesis was done according to the manufacturer's protocol. The cDNA-RNA hybrid was cleaned up using magnetic beads as described above, but with a 1.8X volume and elution in 6 µL.

*cDNA adapter ligation*. A universal i7 DNA adapter for multiplexing compatible with Illumina Nextera Next-generation sequencing kits was ordered as a HPLC-purified ssDNA oligo with 5' and 3' phosphorylated ends (Sigma-Aldrich, Table S2). The ligation reactions were set up as described before (see 3'end ligation), and the reaction was incubated for 16 h at 16 °C. The ligation product was cleaned up using magnetic beads as described above, with a 1.8X volume and elution in 23 µL.

*Library amplification*. Each of the twelve adapter-ligated cDNA pools was amplified using a universal primer for the i5 index and one of twelve primers binding the universal i7 adapter (Table S2) and containing a barcode, introducing unique i7 barcodes to each of the pooled cDNA samples which allowed for pooling samples prior to sequencing (Table S2, Supplementary data, SD_15). The amplification reaction was set up containing 1X NEBNext Ultra II Q5 Master Mix ()NEB, M0544S), 22 µL of ligation product, 1.5 µM of each primer and amplification occurred by firs melting the cDNA for 30 s at 98 °C, followed by 16 cycles of 10 s at 98 °C, 75 s at 63 °C, 10 s at 72 °C, and a final elongation of 5 minutes by 72 °C. The amplification product was cleaned up using magnetic beads as described above, with a 1.2X volume and elution in 20 µL.

*Sequencing and data processing*. The twelve pools were combined in four pools and sent for short-read (150 bp) paired-end sequencing using Illumina NovaSeq X plus sequencing chemistry, performed at Novogene, UK. Raw data is stored at The Sequence Read Archive (SRA, NCBI) at https://www.ncbi.nlm.nih.gov/bioproject/ and can be accessed with BioProject ID PRJNA1472557. The resulting fastq files were processed using a the European Galaxy server (https://usegalaxy.eu/, (35)). Fastq files and a FASTA reference of the *E. coli* DH10b genome and the different plasmids containing 3'UTR variants were aligned using the bwa_mem2 algorithm (version 2.2.1). The resulting .bam files were used to calculate per-base coverage using mosdepth (version 0.3.8). Statistics on coverage and coverage depth were calculated using samtools (version 1.15.1). The number of reads per genetic feature was determined using FeatureCounts (version 2.0.3). The Galaxy workflow is deposited online, together with all .fasta files used for alignment (Zenodo, doi: 10.5281/zenodo.20717389). Coverage was visualized using in R (version 4.4.1) using the BioConductor Rsamtools package (36)

#### Affinity chromatography of proteins

*RNAP/sigma70*: Proteins were produced in *E. coli* BL21-Codon Plus (DE3) (Agilent, #230245), induced in 400 µM IPTG, grown overnight at 17 °C and 180 RPM. Cells were resuspended in lysis buffer (50 mM Tris-HCl pH 6.9, 500 mM NaCl, 10 % glycerol, 1 mM DTT, protease inhibitors [Roche] and 0.3 mM PMSF). Bacteria were lysed by sonication on ice with an amplitude of 40 % for 2.5 min, 5s on, 5s off). Lysate was spun at 12,000 ×*g* for 20 min at 4 °C and supernatant was used for protein purification using affinity chromatography. Supernatant was run over an Akta Pure equipped with a Cytiva HisTrap HP colomn 5-ml (Cytiva cat # 17524802). Proteins were washed using wash buffer (50 mM Tris-HCl pH 6.9, 500 mM NaCl, 10% glycerol, 50 mM imidazole pH 7.6, 0,5 mM DTT) and eluted in elution buffer (As wash buffer, but with 250 mM imidazole). Protein-containing fractions were pooled and dialyzed 2x 2 hours against RNAP dialysis buffer (50 mM Tris-HCl pH 6.9, 75 mM NaCl, 0.5 mM EDTA, 10% glycerol, 0,5 mM DTT) in a Slide-A-Lyzer Dialysis Cassette, 10.000 MWCO, (Thermo Scientific #66810) and run through a 5-ml Hi Trap Heparin column (Cytiva, #17-0407-01), washed with the same buffer. Protein was eluted by increasing NaCl concentration to 1.5 M. Dialysis and heparin affinity chromatography were repeated, then eluate was loaded on a MonoQ column 5-ml (Cytiva, #17-5179-01), washed with the same dialysis buffer and eluted by increasing NaCl concentration to 1.5 M. Eluate was loaded on a Superdex 200 pg 26/600 column (Cytiva, #28989336), washed with water and eluted in RNAP GF buffer (10 mM Tris-HCl pH 7.5, 100 mM NaCl, 0.1 mM EDTA, 10% glycerol, 0.1 mM DTT).

*NusA/G:* Proteins were produced in *E. coli* BL21-Codon Plus (DE3) (Agilent, #230245), induced in 400 µM IPTG, grown overnight at 17 °C and 180 RPM. Cells were resuspended in lysis buffer (50 mM Tris-HCl pH 6.9, 1.2 M NaCl, 10 % glycerol, 1 mM DTT, protease inhibitors [Roche] and 0.3 mM PMSF). Bacteria were lysed by sonication on ice with an amplitude of 40 % for 2.5 min, 5s on, 5s off). Lysate was spun at 12,000 ×*g* for 20 min at 4 °C and supernatant was used for protein purification using affinity chromatography. Supernatant was run over an Akta Pure equipped with a Cytiva HisTrap HP column 5-ml (Cytiva cat # 17524802). Proteins were washed using wash buffer (20 mM $NaH_2PO_4$ pH 6.9, 500 mM NaCl, 10 % glycerol, 1.5 mM DTT, 50 mM imidazole pH 7.6) and eluted in elution buffer (As wash buffer, but with 400 mM imidazole). Protein-containing fractions were pooled and concentrated in a Amicon Ultra-15 centrifugal filter Ultracell 10k (Sigma Aldrich, UFC901024) at 4000 rpm and purified on a HiLoad 16/600 Superdex 200 prepgrade column (GE Healthcare 28989335) washed with water and eluted in NusA GF buffer (20 mM Tris-HCl pH 7.5, 100 mM $NH_4Cl$, 10 mM MgCl, 0.5 mM EDTA, 10% glycerol, 0.1 mM DTT).

*GreA/B:* Proteins were produced in *E. coli* BL21-Codon Plus (DE3) (Agilent, #230245), induced in 400 µM IPTG, grown overnight at 17 °C and 180 RPM. Cells were resuspended in lysis buffer (50 mM Tris-HCl pH 6.9, 1.2 M NaCl, 10 % glycerol, 0.1 % Tween20, 1 mM DTT, protease inhibitors [Roche] and 0.3 mM PMSF). Bacteria were lysed by sonication on ice with an amplitude of 40 % for 2.5 min, 5s on, 5s off). Lysate was spun at 12,000 ×*g* for 20 min at 4 °C and supernatant was used for protein purification using affinity chromatography. Supernatant was run over an Akta Pure equipped with a Cytiva HisTrap HP column 5-ml (Cytiva cat # 17524802). Proteins were washed using wash buffer (50 mM Tris-HCl pH 6.9, 1.2 M NaCl, 10 % glycerol, 0.1 % Tween20, 1 mM DTT, 50 mM imidazole pH 7.6) and eluted in elution buffer (As wash buffer, but with 250 mM imidazole). Protein-containing fractions were pooled and concentrated in a Amicon Ultra-15 centrifugal filter Ultracell 10k (Sigma Aldrich, UFC901024) at 4000 rpm and purified on a HiLoad 16/600 Superdex 200 prepgrade column (GE Healthcare 28989335) washed with water and eluted in GF buffer (20 mM Tris-HCl pH 7.9,  1 M NaCl, 10 mM MgCl, 0.1 mM EDTA, 10% glycerol, 0.1 mM DTT).

### Supplementary figures

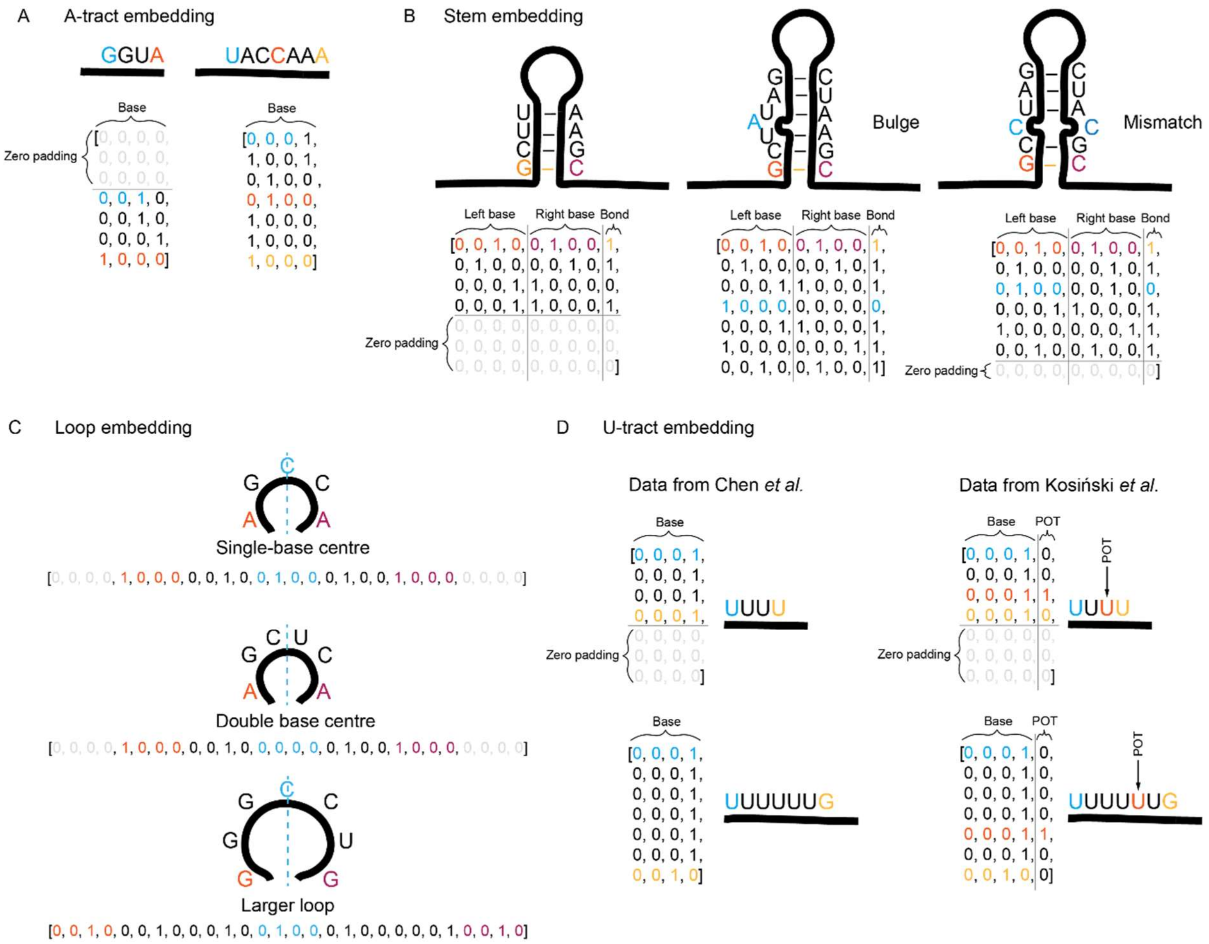


**Figure S1. Embedding strategy for terminators. A.** Embedding of the A-tract (sequences are right-aligned and padded upstream). **B.** Embedding of the stem (base pairs are bottom-aligned and padded towards the loop). The last feature of each base pair representation indicates whether the bases of the pair are bonded or not according to the secondary structure prediction. **C.** Embedding of the loop. Loops are middle-aligned and padded bilaterally. **D.** Embedding of the U-tract (sequences are left-aligned and padded downstream). For Term-Seq data from Kosiński *et al.*, the point of termination was known and was included as a Boolean feature for each base pair.

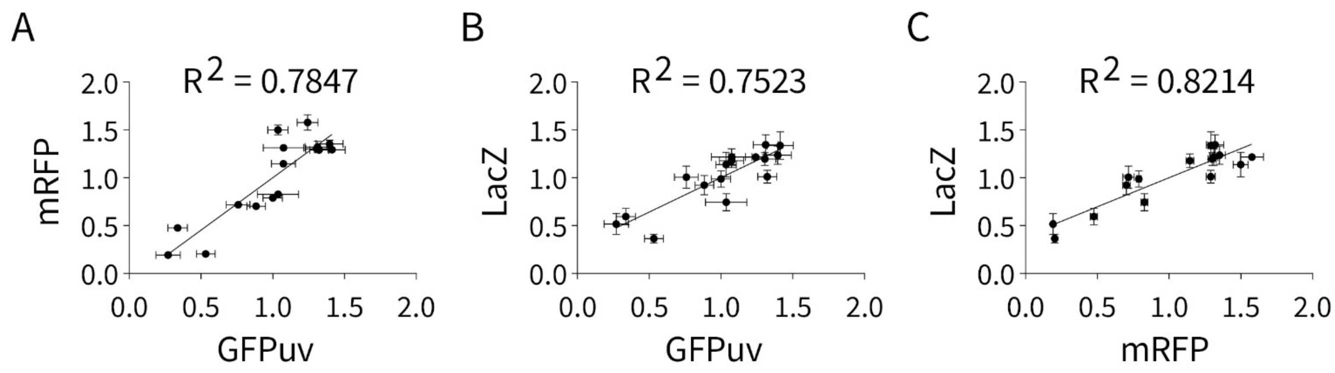


**Figure S2. Pearson correlation coefficients between constructs GFPuv, mRFP and LacZ activity containing spacers 1-16 in the 3'UTR.** Dots show mean normalized protein signal per spacer sequence for (A) mRFP vs GFPuv fluorescence, (B) LacZ activity vs. GFPuv fluorescence and (C) LacZ activity vs mRFP fluorescence. Error bars indicate standard deviation. Pearson correlation coefficient $R^2$ is plotted as a line and shown in each graph.

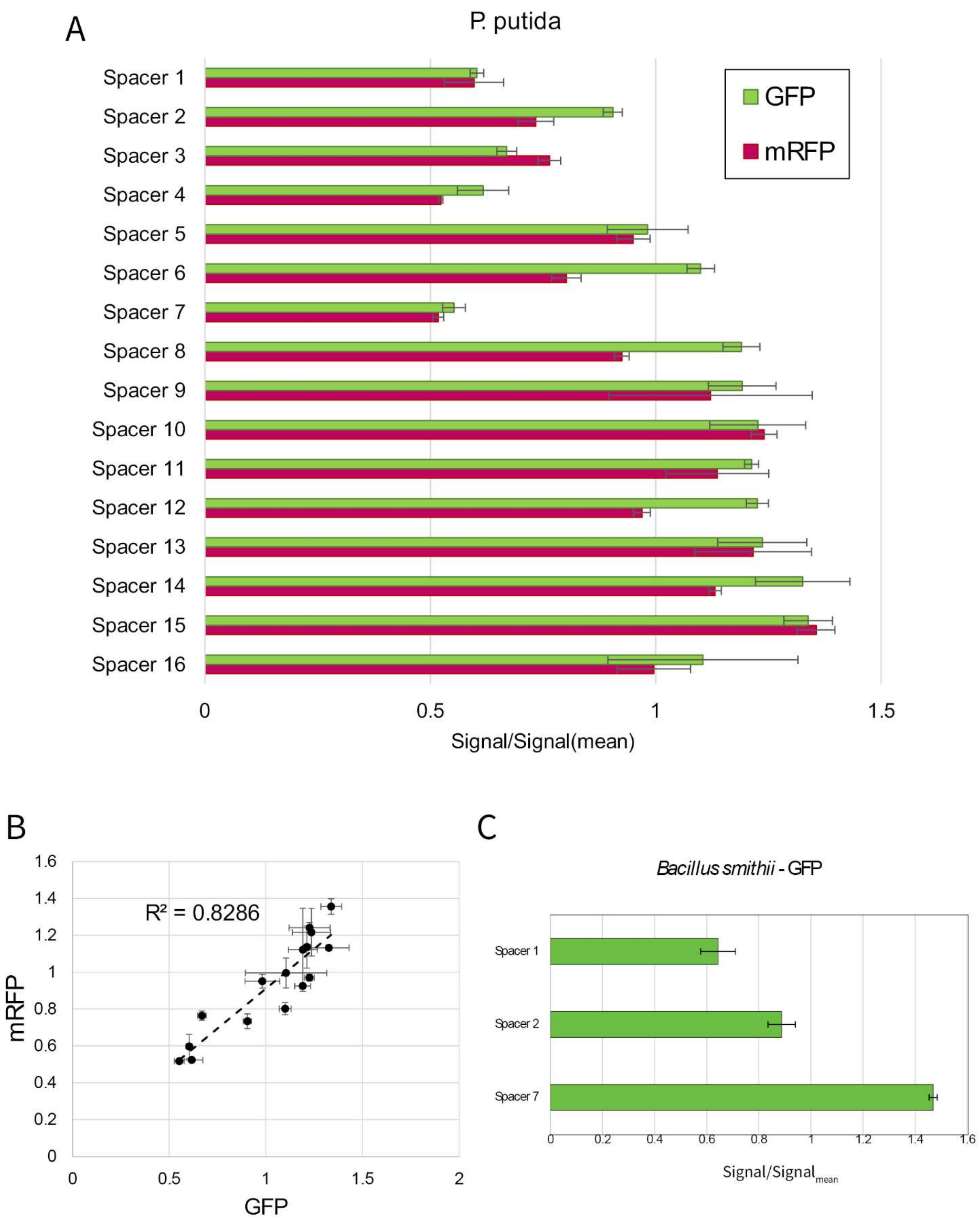


**Figure S3 Analysis of the effect of 3'UTR spacer sequences on protein production in *Pseudomonas putida*.** (A) Effect of spacer sequences 1-16 on RFP and GFP production in *P.*

*putida*. Protein activity of mRFP (magenta) and GFPuv (green) was determined by measuring fluorescence in a plate reader assay, To compare relative protein activity, the activity of each protein was divided by the mean activity over all spacers, resulting in a normalized signal. (B) Pearson correlation coefficients between GFP and mRFP fluorescence of constructs containing spacers 1-16 in the 3'UTR in *P. putida*. (C). Effect of spacers 1, 2 and 7 on GFP production in gram-positive *Bacillus smithii*.

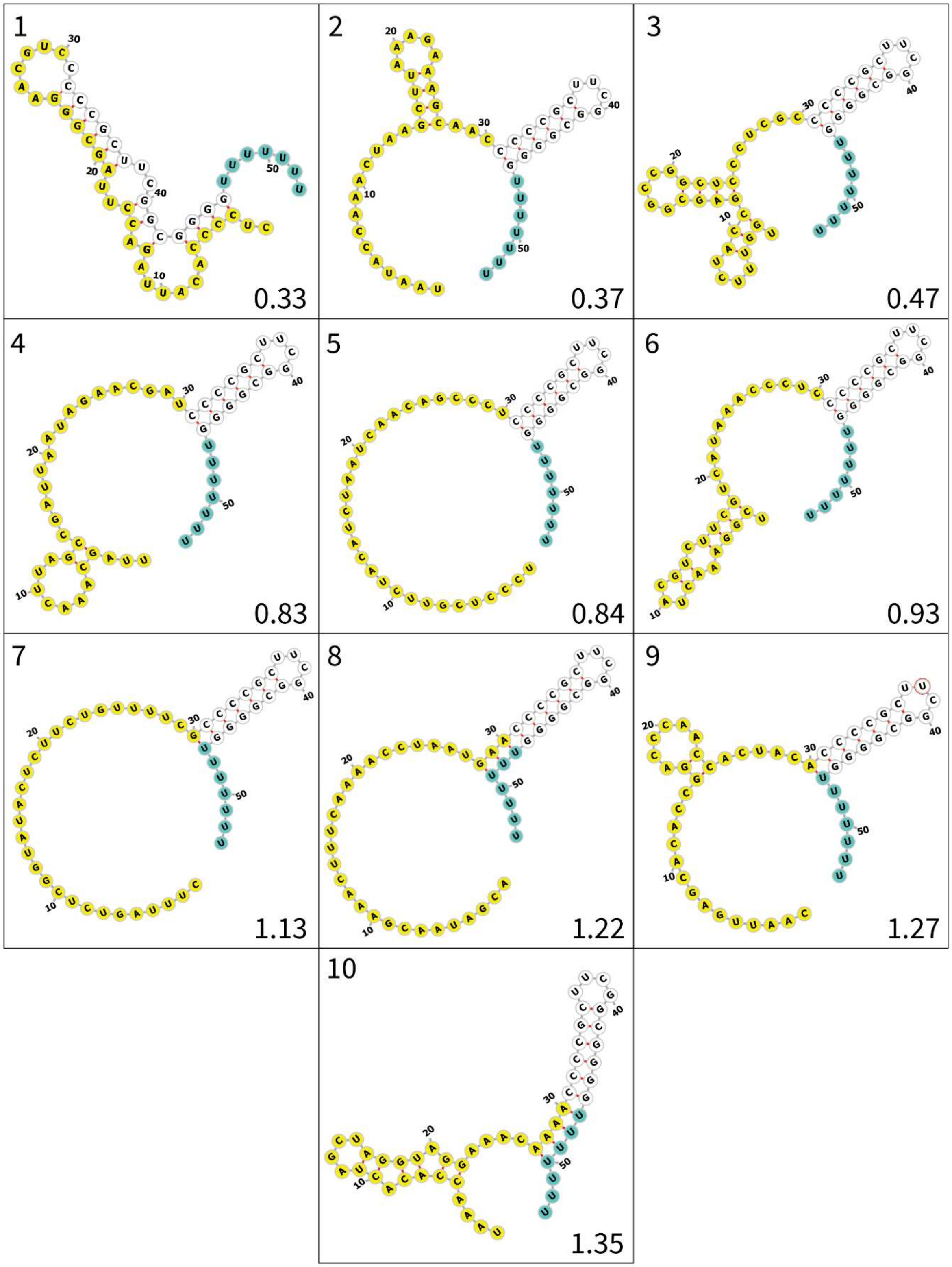


**(figure continues on next page)**

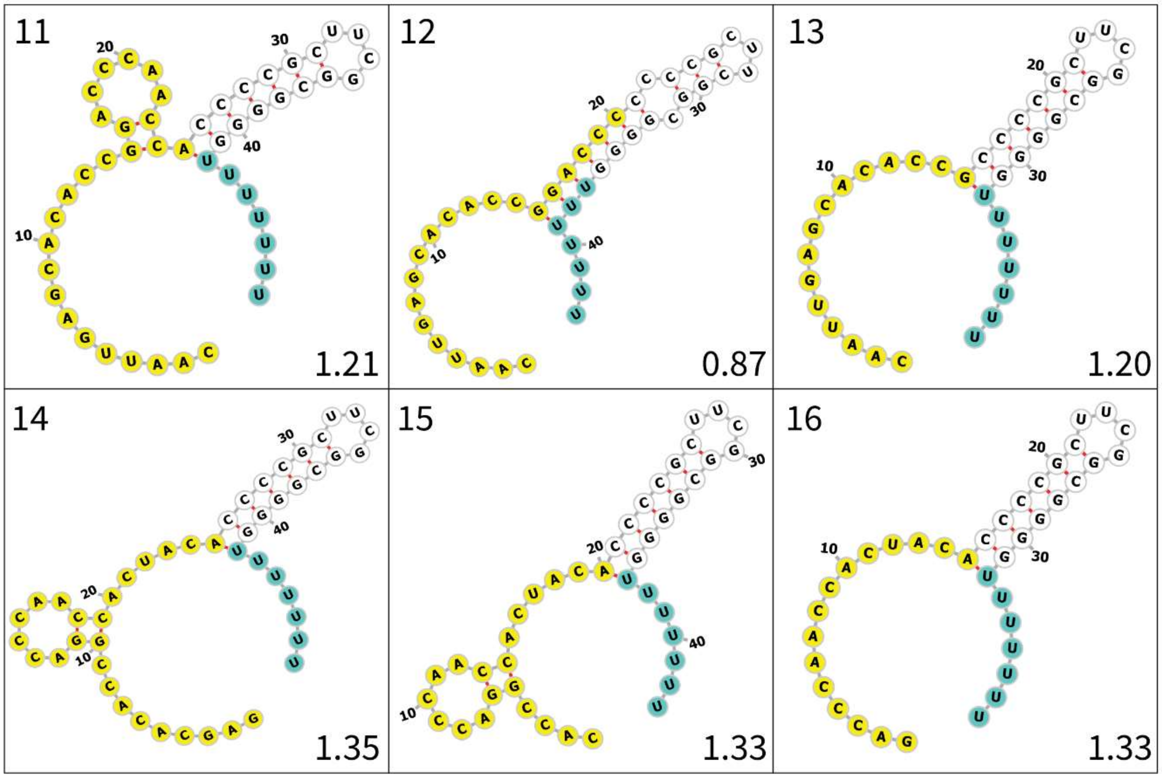


**Figure S4. Secondary structures of 3'UTRs with spacer variants 1-16.** RNAFold predictions of secondary structures of 3'UTR regions containing selected spacers. The spacer number is indicated in the top left corner, the mean signal corresponding to each signal is indicated in the bottom right corner. The spacer sequence is indicated in yellow, the poly(U)-tail is indicated in blue. Structures were visualized using Forna.

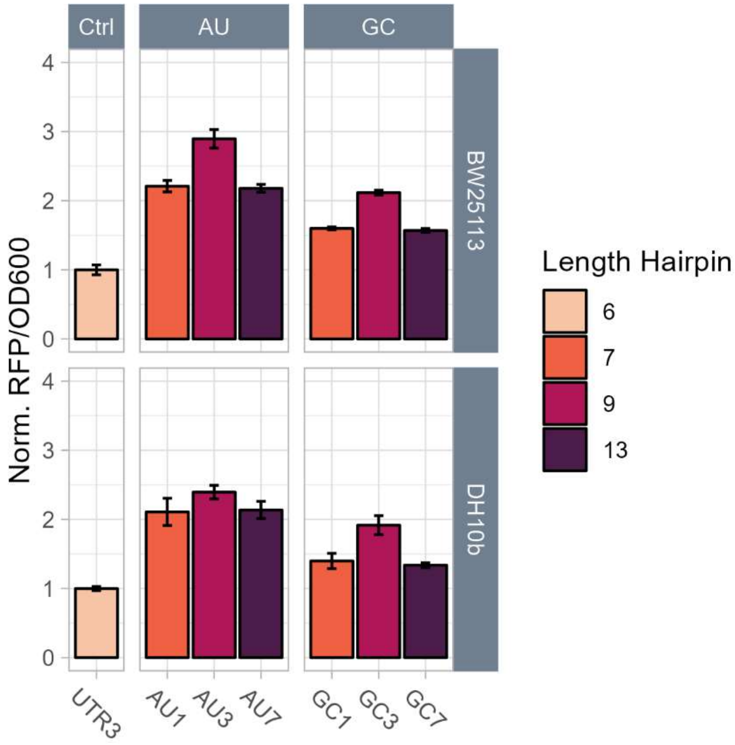


**Figure S5. Extending the terminator hairpin by A-U and G-C base pairing.** The control spacer 3 (UTR3) was modified by inserting an (A)-tract of one, three and seven adenine bases between the spacer and the terminator hairpin (AU1, AU3, AU7, resp.). Similarly, the 3'UTR containing spacer 3 was modified by extending the terminator hairpin with one, three and seven G-C base pairs in the bottom of the stem (GC1, GC3 and GC7, resp.). Each UTR was cloned behind an RFP gene to investigate the effect of each 3'UTR on protein production. Bars represent mean RFP fluorescence measured by plate reader normalized over OD600 of the culture and relative to the fluorescence of the control, from three biological replicates, error bars indicate standard deviation. Colors of the bars indicate the length of the hairpin predicted by RNAfold. The experiment was performed in both *E. coli* strain BW25113 and DH10B.

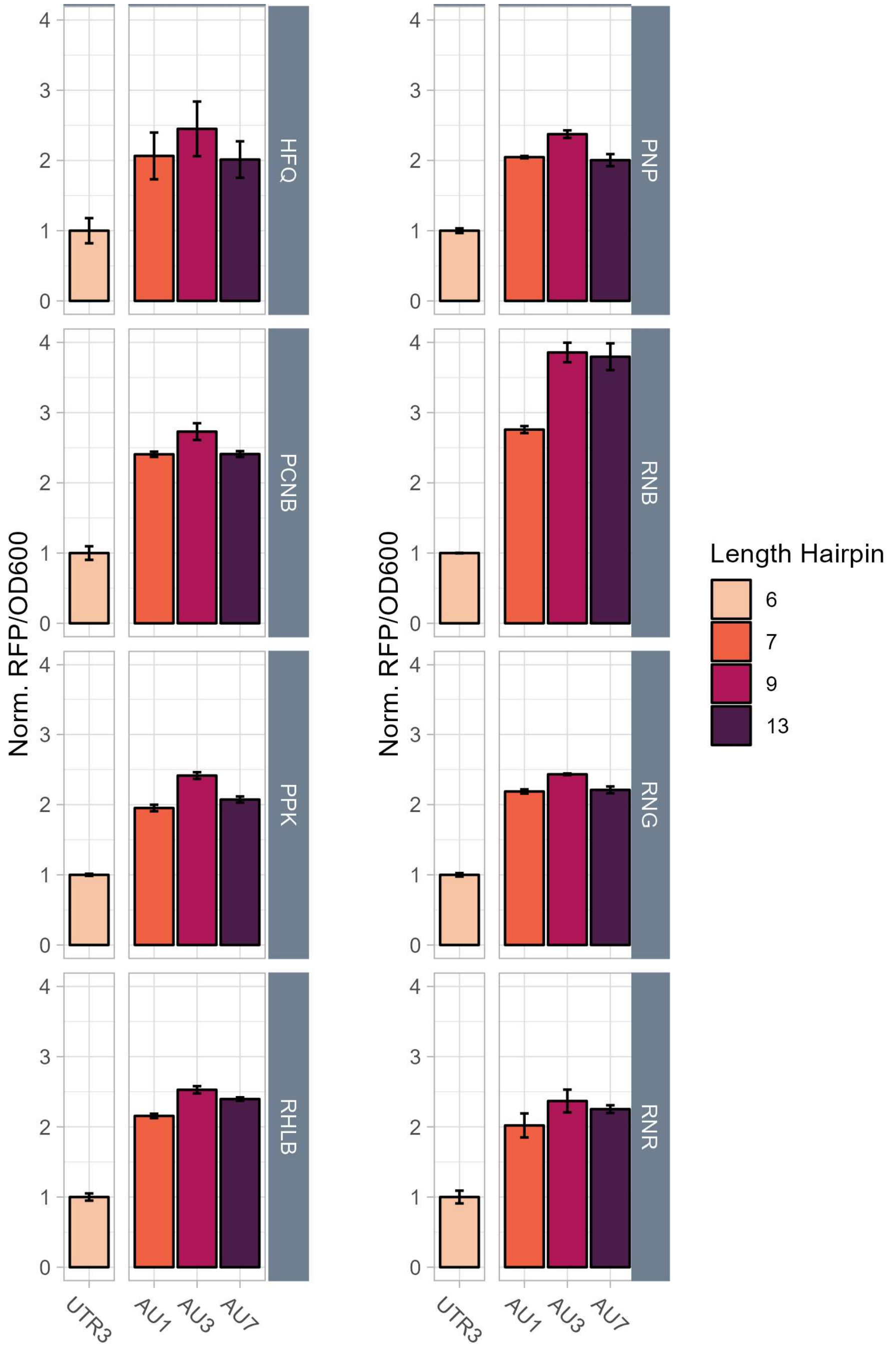


**Figure S6. Effect of genomic deletion of inessential mRNA turnover machinery on protein production of 3'UTRs containing an (A)-tract.** The control spacer 3 (UTR3) was modified by inserting an (A)-tract of one, three and seven adenine bases between the spacer and the terminator hairpin (AU1, AU3, AU7, resp.) Each UTR was cloned behind an RFP gene to investigate the effect of each 3'UTR on protein production and transformed in a strain of the KEIO collection harboring a genomic deletion of a gene essential in mRNA turnover. Respective gene deletions are indicated on the right of each graph. Bars represent mean RFP fluorescence measured by plate reader normalized over OD600 of the culture and relative to the fluorescence of the control, from three biological replicates, error bars indicate standard deviation. Colors of the bars indicate the length of the hairpin predicted by RNAfold.

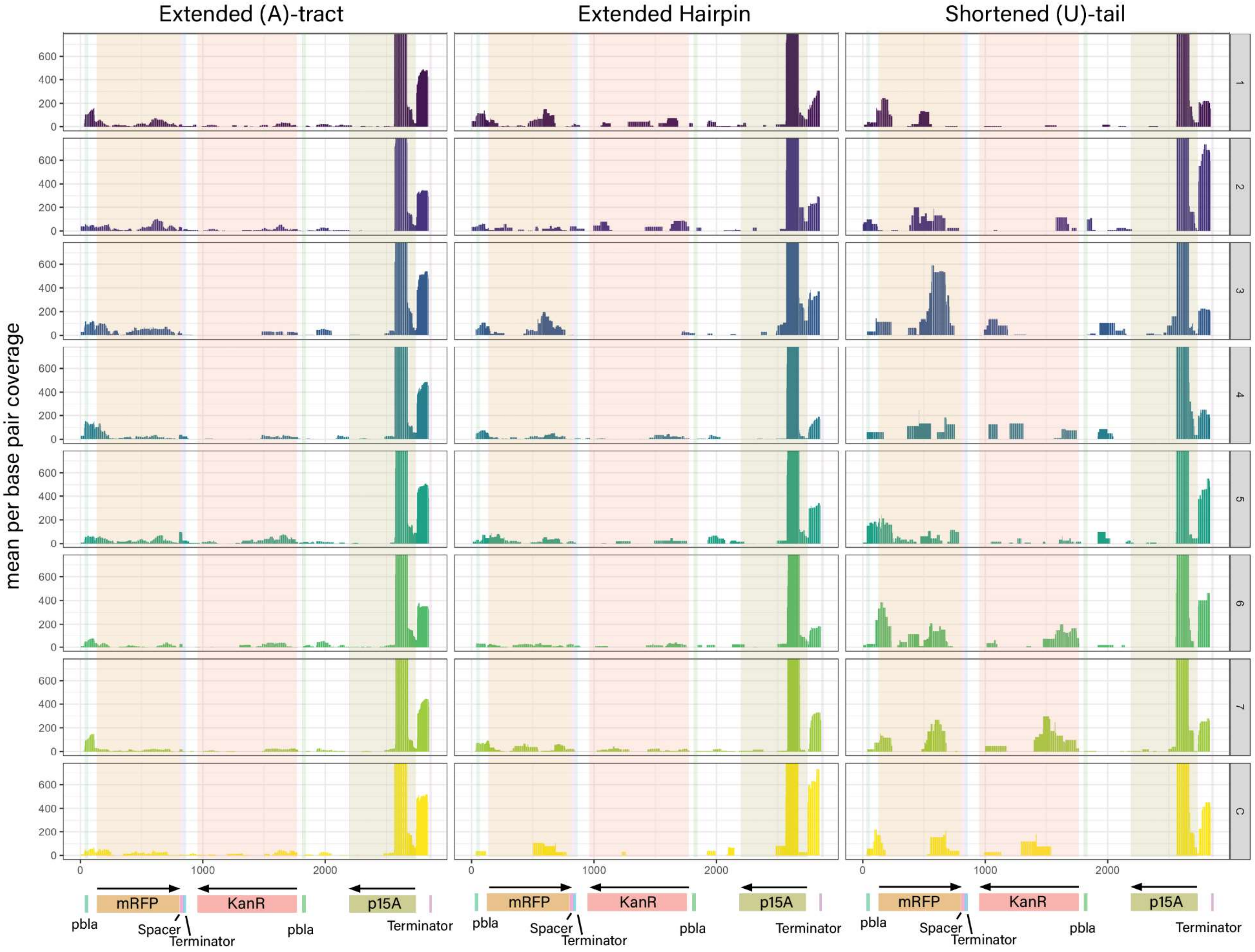

Extended (A)-tract
Extended Hairpin
Shortened (U)-tail
mean per base pair coverage
0
200
400
600
1
2
3
4
5
6
7
C
0
1000
2000
mRFP
KanR
p15A
pbla
Spacer
Terminator
pbla
Terminator

**Figure S7. Per-base coverage of a TERM-seq dataset of plasmids containing all 3'UTR rational variants.** The y-axis represents the mean per-base coverage of TERM-seq datasets of three biological replicates, specifically sequencing the 3' ends of mRNA in the cell. The Y axis represents the base pair position of the plasmid sequence used to express the different UTR variants, a schematic of the plasmid map is also depicted below each column and the shaded areas correspond to the respective positions of the plasmid map. The first column shows sequencing data from samples with an (A)-tract upstream of the terminator, with (A)-tracts between 1 and 7 adenine bases long, corresponding to the numbers on the right of the figure. The middle column contains samples with a hairpin extended with 1 up to 7 U-A base pairs, corresponding to the numbers on the right. The right column shows sequencing data for 3'UTRs where the poly(U)-tail was removed, where 7 means 7 uracil bases were removed, resulting in no (U)-tail while 1 means one uracil base was removed, resulting in a (U)-tail of six uracil bases. The bottom row (C) shows sequencing coverage of a plasmid containing a 3'UTR with control spacer 3.

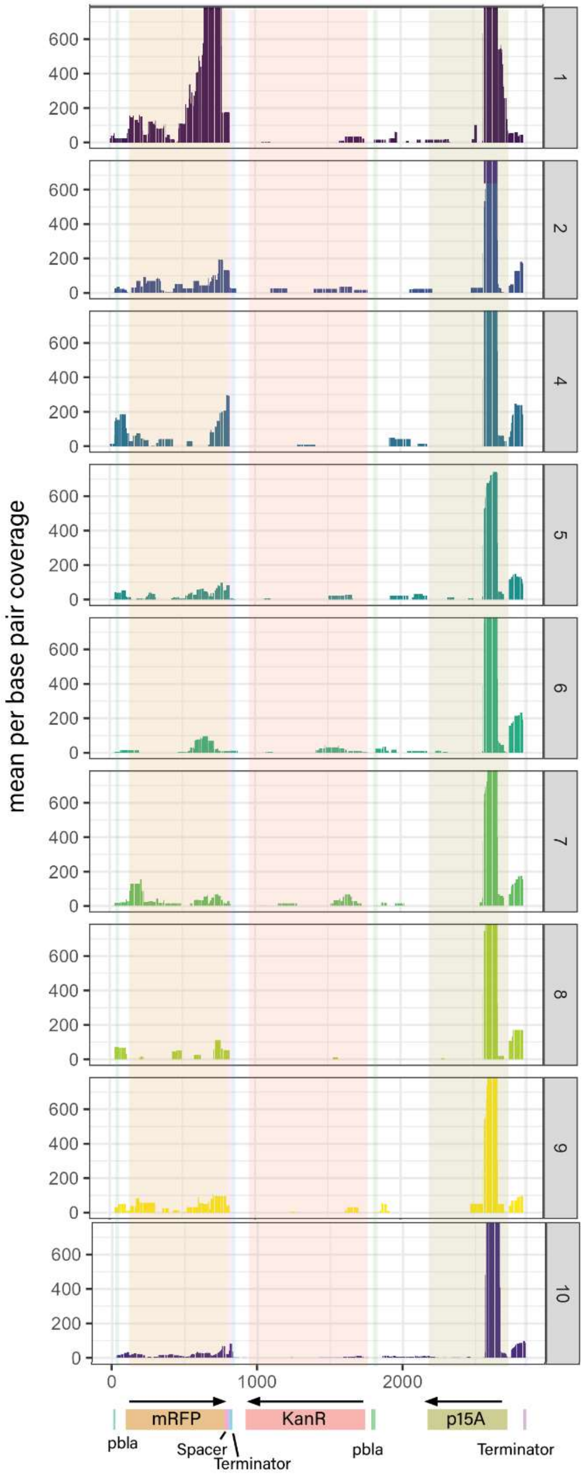


**Figure S8. Per-base coverage of a TERM-seq dataset of plasmids harboring 3'UTR spacers 1-10.** The y-axis represents the mean per-base coverage of TERM-seq datasets of three biological replicates, specifically sequencing the 3' ends of mRNA in the cell. The Y axis represents the base pair position of the plasmid sequence used to express the different UTR variants, a schematic of the plasmid map is also depicted below the graph column and the shaded areas correspond to the respective positions of the plasmid map. The numbers on the right correspond to the spacer numbers as shown in figure 2 of the main text. Sequencing data for spacer 3 is shown in Supplementary figure 7.

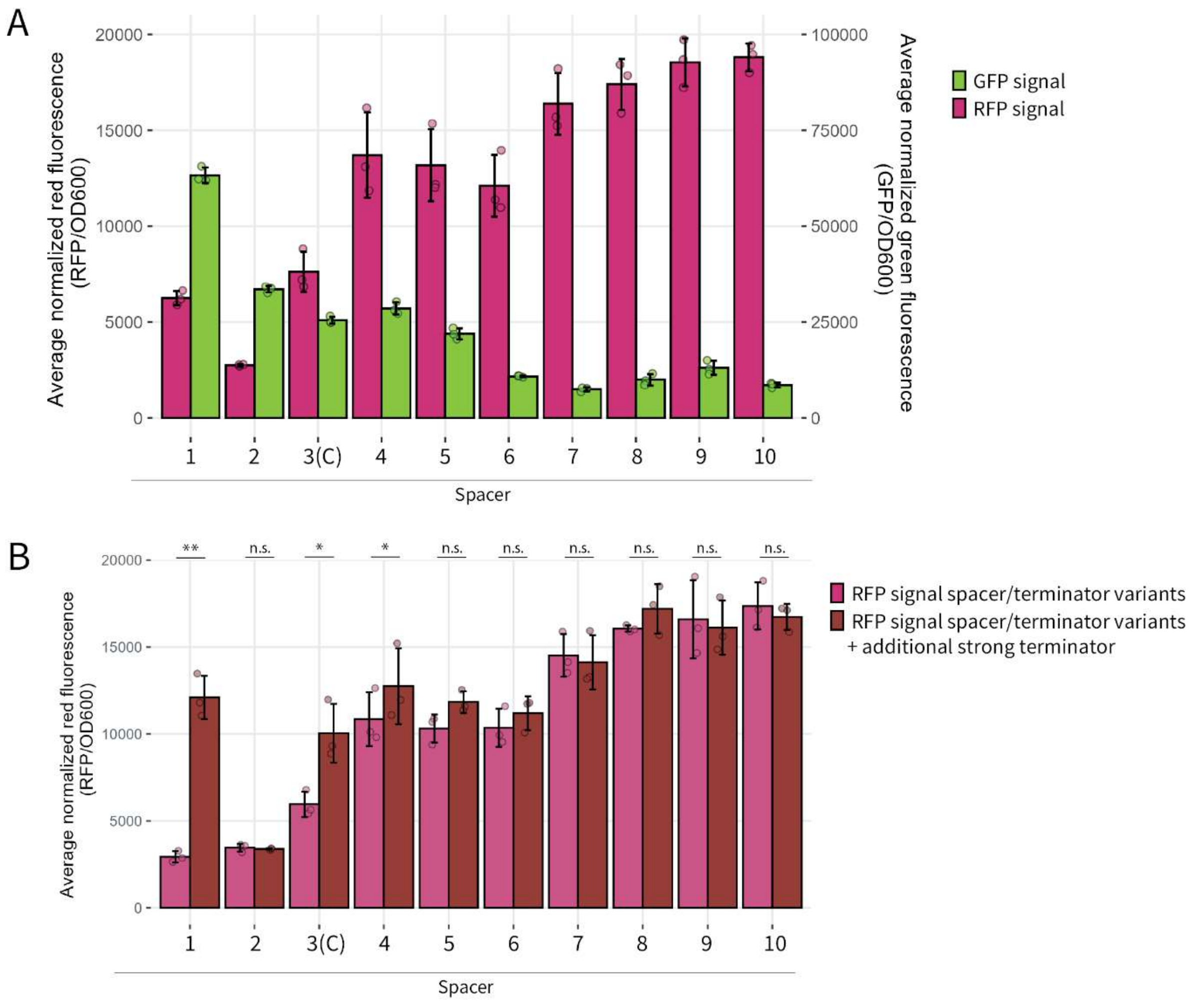


**Figure S9. Termination efficiency of different 3'UTR containing spacers 1-10.** (A) GFP gene without promoter was cloned behind RFP genes with different 3'UTR spacers 1-10 (corresponding to figure 2) in order to assess readthrough. Bars show average RFP fluorescence (magenta) and average GFP fluorescence (green) of three biological replicates (circles). (B) RFP fluorescence of an RFP gene with 3'UTRs containing spacers 1-10 were compared to the same constructs with an additional terminator with high termination efficiency. Bars show average RFP fluorescence with the 3'UTR variants alone and (magenta) and average RFP fluorescence of constructs with a second terminator with high TE (dark red) of three biological replicates (circles). In both panels, fluorescence was measured by plate reader and normalized over $OD_{600}$ of the cultures. Biological variation is shown by error bars, and statistical significance between mean values is denoted with asterisks.Black asterisks show the statistical significance between specific samples, with the following significance levels: * $p < 0.05$, ** $p < 0.005$ and n.s., not significant.

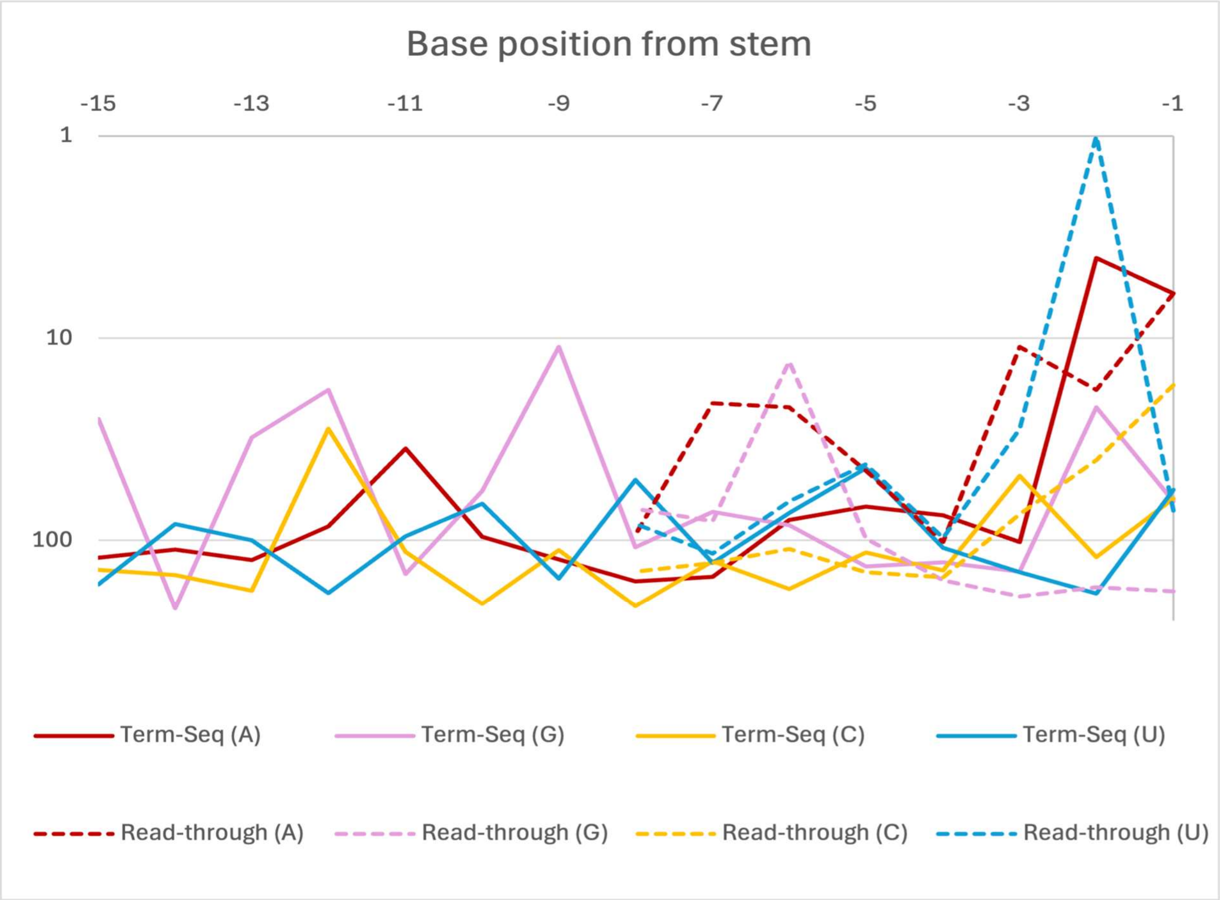


**Figure S10. Feature rank plotted against base position of the spacer.** The eight lines show the feature ranks of the feature 'is this base an A/C/G/U?' at the indicated positions for each of the two random forest regressors trained on the full Term-Seq/Read-through datasets. The X-axis displays the base position from the stem. The Y-axis displays the rank of the feature 'Is the $X^{th}$ base of the spacer an A/C/G/U'? The feature 'is this base an A' is substantially more important than other spacer features in the 1-2 bases directly upstream of the hairpin (with the exception of one outlier, 'is this base a U', at position -2).

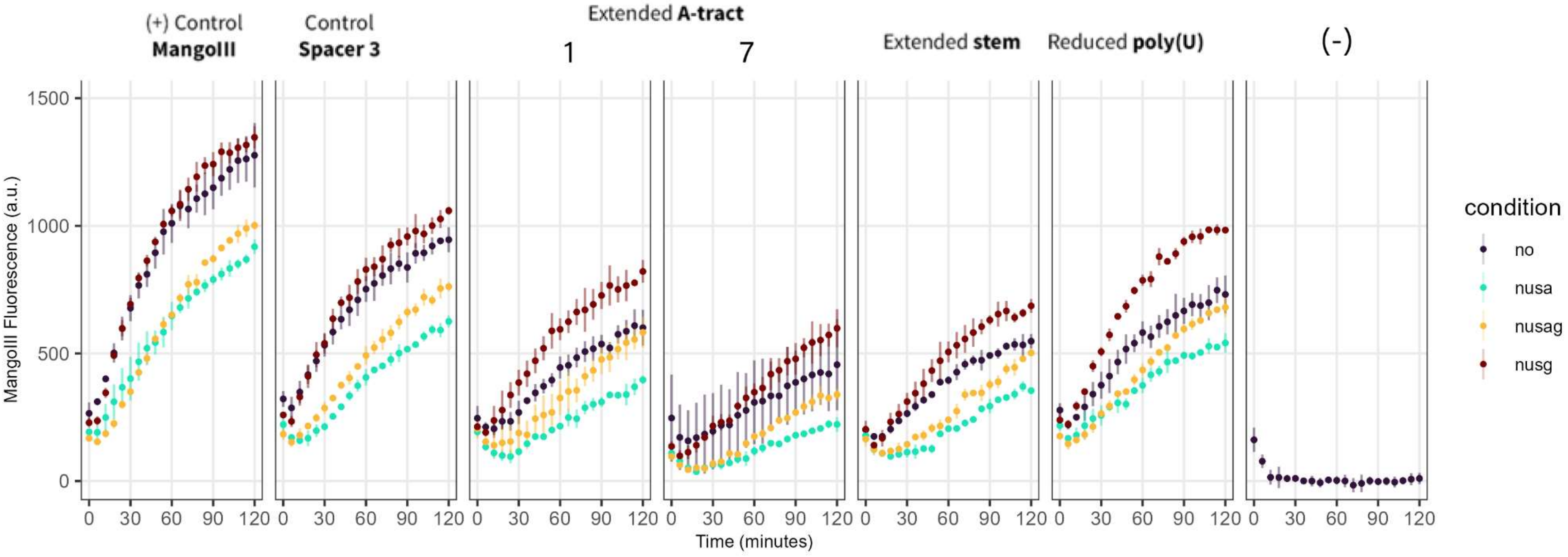


**Figure S11. Mango-III fluorescence as a proxy for transcription over time in an *in vitro* transcription assay in presence or absence of NusA and/or NusG.** Upstream of each MangoIII aptamer was no terminator (+ control), a terminator containing control spacer 3, a spacer with an A-tract of 1 or 7 , an extended hairpin stem or a shortened poly(U)-tail. The negative control did not contain a MangoIII aptamer. Circles represent mean values of three technical replicates at each given time point, vertical bars present the standard deviation of each measurement. Colors represent the different conditions: No proteins added (blue), NusA (blue), NusG (red), NusA+NusG (yellow). Using a sliding window method, the maximum transcription rate was calculated from each slope

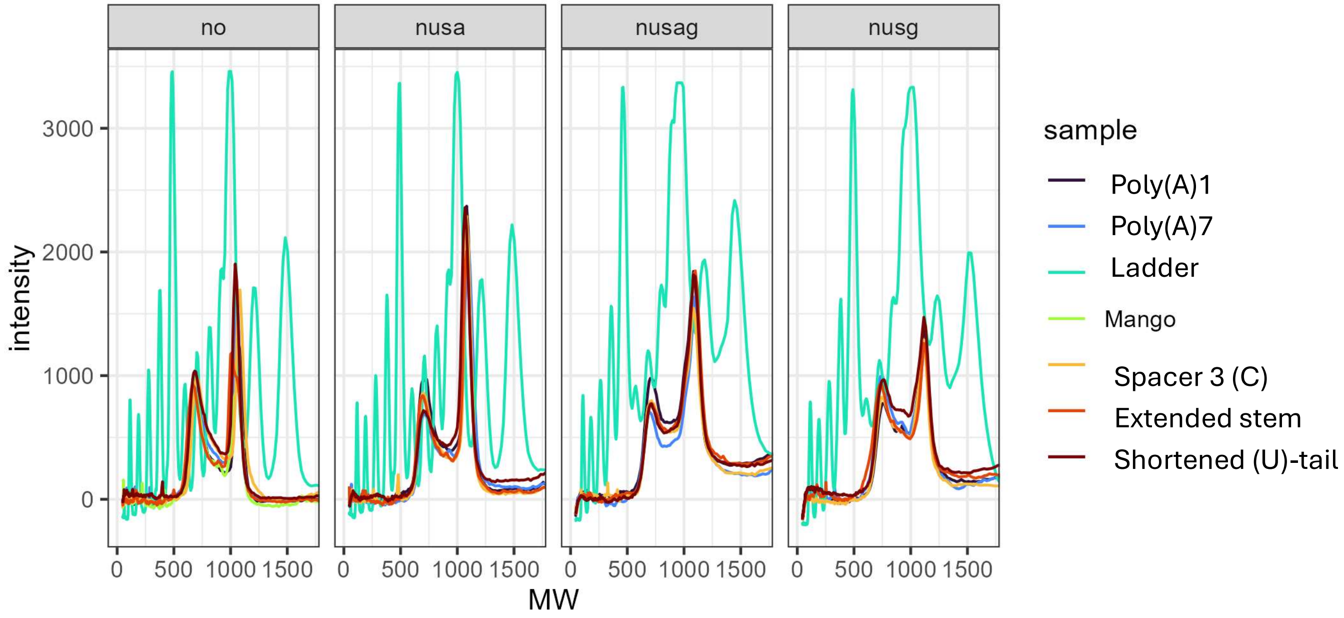


**Figure S12. Transcription products after *in vitro* transcription assay.** Transcription products were loaded on a 1,2 % agarose gel and separated for 30 minutes at 100V. Profiles of each lane were quantified using imageJ Gel plotting Macro and the average of each lane profile per strain per condition determined and presented here. Products were analyzed per condition, with no enzymes added (no), NusA added (nusa), NusG added (nusg) and Nus A and NusG added (nusag). The molecular weight on the X-axis is determined using the NEB 1kb plus molecular weight marker (Ladder) plotted in light green. Intensity corresponds to the pixel intensity measured using ImageJ and can be used as a proxy for RNA abundance.

#### Supplementary tables

**Table S1. 3'UTR sequences used in this study.** 3'UTRs were directly placed downstream of the STOP-codon of the upstream gene.

| 3'UTR indicator | Origin and Modification | Hairpin length[1] | Free poly(U)-length[1] | 3'UTR sequence[3] |
|---|---|---|---|---|
| Spacer 1 | Selected from random library[2] | 6 | 7 | CTCCCCACATTAGACCTTAGCGGGAACGTCCCCCGCTTCGGCGGGGTTTTTTT |
| Spacer 2 | Selected from random library[2] | 0 | 7 | TAATACCAAACTAAGCTTAAAGAAAGCAACCCCCGCTTCGGCGGGGTTTTTTT |
| Spacer 3 | Selected from random library[2] | 6 | 7 | **TGGTTTCTACCGAGCGGCCGGCTCCCTCGC**CCCCGCTTCGGCGGGGTTTTTTT |
| Spacer 4 | Selected from random library[2] | 6 | 7 | TTAGCAAACTTAGCCGATTAATAGAACGATCCCCGCTTCGGCGGGGTTTTTTT |
| Spacer 5 | Selected from random library[2] | 6 | 7 | TCCCTCGTTCTACATCTAATCAACAGCCCTCCCCGCTTCGGCGGGGTTTTTTT |
| Spacer 6 | Selected from random library[2] | 6 | 7 | TCGGAAACTACGTCTTCGTCATAAACCCTCCCCCGCTTCGGCGGGGTTTTTTT |
| Spacer 7 | Selected from random library[2] | 7 | 6 | CTTTAGTCTCGGTATACTCTTCTGTTTTCGCCCCGCTTCGGCGGGGTTTTTTT |
| Spacer 8 | Selected from random library[2] | 9 | 4 | ACGATAACGAAACTTTCAAAACCTAATGAACCCCGCTTCGGCGGGGTTTTTTT |
| Spacer 9 | Selected from random library[2] | 7 | 6 | CAATTGAGCACACCGGACCCAACCACTACACCCCGCTTCGGCGGGGTTTTTTT |
| Spacer 10 | Selected from random library[2] | 10 | 3 | TAAACCACACTAGCTAGGTAGGAAACAAAACCCCGCTTCGGCGGGGTTTTTTT |
| Spacer 11 | Spacer 9 shortened from 3'end | 7 | 6 | CAATTGAGCACACCGGACCCAACCACCCCGCTTCGGCGGGGTTTTTTT |

| | | | | |
|---|---|---|---|---|
| Spacer 12 | Spacer 9 shortened from 3'end | 11 | 4 | CAATTGAGCACACCGGACCCCCCCGCTTCGGCGGGGTTTTTTT |
| Spacer 13 | Spacer 9 shortened from 3'end | 7 | 6 | CAATTGAGCACACCGCCCCGCTTCGGCGGGGTTTTTTT |
| Spacer 14 | Spacer 9 shortened from 5'end | 7 | 6 | GAGCACACCGGACCCAACCACTACACCCCGCTTCGGCGGGGTTTTTTT |
| Spacer 15 | Spacer 9 shortened from 5'end | 7 | 6 | CACCGGACCCAACCACTACACCCCGCTTCGGCGGGGTTTTTTT |
| Spacer 16 | Spacer 9 shortened from 5' end | 7 | 6 | GACCCAACCACTACACCCCGCTTCGGCGGGGTTTTTTT |
| Extended A-tract 1 | Spacer 3 - 1 A between spacer and hairpin | 7 | 6 | **TGGTTTCTACCGAGCGGCCGGCTCCCTCGC**ACCCCGCTTCGGCGGGGTTTTTTT |
| Extended A-tract 2 | Spacer 3 - 2 A between spacer and hairpin | 8 | 5 | **TGGTTTCTACCGAGCGGCCGGCTCCCTCGC**AACCCCGCTTCGGCGGGGTTTTTTT |
| Extended A-tract 3 | Spacer 3 - 3 A between spacer and hairpin | 9 | 4 | **TGGTTTCTACCGAGCGGCCGGCTCCCTCGC**AAACCCCGCTTCGGCGGGGTTTTTTT |
| Extended A-tract 4 | Spacer 3 - 4 A between spacer and hairpin | 10 | 3 | **TGGTTTCTACCGAGCGGCCGGCTCCCTCGC**AAAACCCCGCTTCGGCGGGGTTTTTTT |
| Extended A-tract 5 | Spacer 3 - 5 A between spacer and hairpin | 11 | 2 | **TGGTTTCTACCGAGCGGCCGGCTCCCTCGC**AAAAACCCCGCTTCGGCGGGGTTTTTTT |
| Extended A-tract 6 | Spacer 3 - 6 A between spacer and hairpin | 12 | 1 | **TGGTTTCTACCGAGCGGCCGGCTCCCTCGC**AAAAAACCCCGCTTCGGCGGGGTTTTTTT |
| Extended A-tract 7 | Spacer 3 - 7 A between spacer and hairpin | 13 | 0 | **TGGTTTCTACCGAGCGGCCGGCTCCCTCGC**AAAAAAACCCCGCTTCGGCGGGGTTTTTTT |
| Extended stem 1 | Spacer 3 - 1 U-A at bottom of hairpin | 7 | 7 | **TGGTTTCTACCGAGCGGCCGGCTCCCTCGC**UCCCCGCTTCGGCGGGGATTTTTTT |
| Extended stem 2 | Spacer 3 - 2 U-A at bottom of hairpin | 8 | 7 | **TGGTTTCTACCGAGCGGCCGGCTCCCTCGC**UUCCCCGCTTCGGCGGGGAATTTTTTT |
| Extended stem 3 | Spacer 3 - 3 U-A at bottom of hairpin | 9 | 7 | **TGGTTTCTACCGAGCGGCCGGCTCCCTCGC**UUUCCCCGCTTCGGCGGGGAAATTTTTTT |
| Extended stem 4 | Spacer 3 - 4 U-A at bottom of hairpin | 10 | 7 | **TGGTTTCTACCGAGCGGCCGGCTCCCTCGC**UUUUCCCCGCTTCGGCGGGGAAAATTTTTTT |
| Extended stem 5 | Spacer 3 - 5 U-A at bottom of hairpin | 11 | 7 | **TGGTTTCTACCGAGCGGCCGGCTCCCTCGC**UUUUUCCCCGCTTCGGCGGGGAAAAATTTTTTT |

| | | | | |
|---|---|---|---|---|
| Extended stem 6 | Spacer 3 - 6 U-A at bottom of hairpin | 12 | 7 | **TGGTTTCTACCGAGCGGCCGGCTCCCTCGC**UUUUUUCCCCGCTTCGGCGGGGAAAAAATTTTTTT |
| Extended stem 7 | Spacer 3 - 7 U-A at bottom of hairpin | 13 | 7 | **TGGTTTCTACCGAGCGGCCGGCTCCCTCGC**UUUUUUUCCCCGCTTCGGCGGGGAAAAAAATTTTTTT |
| Shortened (U)-tail 0 | Spacer 3 - 0 U after hairpin | 6 | 0 | **TGGTTTCTACCGAGCGGCCGGCTCCCTCGC**CCCCGCTTCGGCGGGG |
| Shortened (U)-tail 1 | Spacer 3 - 1 U after hairpin | 6 | 1 | **TGGTTTCTACCGAGCGGCCGGCTCCCTCGC**CCCCGCTTCGGCGGGGT |
| Shortened (U)-tail 2 | Spacer 3 - 2 U after hairpin | 6 | 2 | **TGGTTTCTACCGAGCGGCCGGCTCCCTCGC**CCCCGCTTCGGCGGGGTT |
| Shortened (U)-tail 3 | Spacer 3 - 3 U after hairpin | 6 | 3 | **TGGTTTCTACCGAGCGGCCGGCTCCCTCGC**CCCCGCTTCGGCGGGGTTT |
| Shortened (U)-tail 4 | Spacer 3 - 4 U after hairpin | 6 | 4 | **TGGTTTCTACCGAGCGGCCGGCTCCCTCGC**CCCCGCTTCGGCGGGGTTTT |
| Shortened (U)-tail 5 | Spacer 3 - 5 U after hairpin | 6 | 5 | **TGGTTTCTACCGAGCGGCCGGCTCCCTCGC**CCCCGCTTCGGCGGGGTTTTT |
| Shortened (U)-tail 6 | Spacer 3 - 6 U after hairpin | 6 | 6 | **TGGTTTCTACCGAGCGGCCGGCTCCCTCGC**CCCCGCTTCGGCGGGGTTTTTT |
| Control | Same as Spacer 3 | 6 | 7 | **TGGTTTCTACCGAGCGGCCGGCTCCCTCGC**CCCCGCTTCGGCGGGGTTTTTTT |
| Extended stem 1 GC[4] | Spacer 3 - 1 G-C at bottom of hairpin | 7 | 7 | **TGGTTTCTACCGAGCGGCCGGCTCCCTCGC**GCCCCGCTTCGGCGGGGCTTTTTTT |
| Extended stem 3 GC[4] | Spacer 3 - 3 G-C at bottom of hairpin | 10 | 7 | **TGGTTTCTACCGAGCGGCCGGCTCCCTCGC**GGGCCCCGCTTCGGCGGGGCCCTTTTTTT |
| Extended stem 7 GC[4] | Spacer 3 – 7 G-C at bottom of hairpin | 13 | 7 | **TGGTTTCTACCGAGCGGCCGGCTCCCTCGC**GGGGGGGCCCCGCTTCGGCGGGGCCCCCCCTTTTTTT |

[1] As predicted by RNAfold secondary structure predictions. Details are found in the raw supplementary data file (supplementary_data.xslx, SD_1) accompanying the manuscript

[2] Random library sequences are described in the raw supplementary data file (supplementary_data.xslx, SD_2) accompanying the manuscript

[3] The starting sequence of terminator BBa_B1002 is underlined, spacer 3, the reference sequence for most experiments is highlighted.

[4] This construct has only been described in the supplementary data (Figure S5)

**Table S2. ssDNA oligos used in this study**

| Identifier | Sequence | Name | Purpose |
|---|---|---|---|
| BG18017 | ggtctcaGGGGNNNNNNNNNNNNNNNNNNNNNNNNNNNNNNTTAA tgagaccAATTCAGCTACGCT | Spacer library | cloning 3'UTR spacer library behind GFP/RFP |
| BG18018 | AGCGTAGCTGAATTggtctc | spacer library | cloning 3'UTR spacer library behind GFP/RFP |
| BG20363 | TAATAATACCAAACTAAGCTTAAAGAAAGCAAC | Spacer 2 FW | Inserting spacer behind GFP/RFP/LacZ |
| BG20364 | GGGGTTGCTTTCTTTAAGCTTAGTTTGGTATTA | Spacer 2 RV | Inserting spacer behind GFP/RFP/LacZ |
| BG20365 | TAATCGGAAACTACGTCTTCGTCATAAACCCTC | Spacer 6 FW | Inserting spacer behind GFP/RFP/LacZ |
| BG20366 | GGGGAGGGTTTATGACGAAGACGTAGTTTCCGA | Spacer 6 RV | Inserting spacer behind GFP/RFP/LacZ |
| BG20367 | TAATCCCTCGTTCTACATCTAATCAACAGCCCT | Spacer 5 FW | Inserting spacer behind GFP/RFP/LacZ |
| BG20368 | GGGAGGGCTGTTGATTAGATGTAGAACGAGGGA | Spacer 5 RV | Inserting spacer behind GFP/RFP/LacZ |
| BG20369 | TAAACGATAACGAAACTTTCAAAACCTAATGAA | Spacer 8 FW | Inserting spacer behind GFP/RFP/LacZ |
| BG20370 | GGGTTCATTAGGTTTTGAAAGTTTCGTTATCGT | Spacer 8 RV | Inserting spacer behind GFP/RFP/LacZ |
| BG20371 | TAATAAACCACACTAGCTAGGTAGGAAACAAAA | Spacer 10 FW | Inserting spacer behind GFP/RFP/LacZ |
| BG20372 | GGGTTTTGTTTCCTACCTAGCTAGTGTGGTTTA | Spacer 10 RV | Inserting spacer behind GFP/RFP/LacZ |
| BG20373 | TAACTCCCCACATTAGACCTTAGCGGGAACGTC | Spacer 1 FW | Inserting spacer behind GFP/RFP/LacZ |
| BG20374 | GGGGACGTTCCCGCTAAGGTCTAATGTGGGGAG | Spacer 1 RV | Inserting spacer behind GFP/RFP/LacZ |
| BG20375 | TAATGGTTTCTACCGAGCGGCCGGCTCCCTCGC | Spacer 3 FW | Inserting spacer behind GFP/RFP/LacZ |
| BG20376 | GGGGCGAGGGAGCCGGCCGCTCGGTAGAAACCA | Spacer 3 RV | Inserting spacer behind GFP/RFP/LacZ |
| BG20377 | TAATTAGCAAACTTAGCCGATTAATAGAACGAT | Spacer 4 FW | Inserting spacer behind GFP/RFP/LacZ |
| BG20378 | GGGATCGTTCTATTAATCGGCTAAGTTTGCTAA | Spacer 4 RV | Inserting spacer behind GFP/RFP/LacZ |
| BG20379 | TAACAATTGAGCACACCGGACCCAACCACTACA | Spacer 9 FW | Inserting spacer behind GFP/RFP/LacZ |
| BG20380 | GGGTGTAGTGGTTGGGTCCGGTGTGCTCAATTG | Spacer 9 RV | Inserting spacer behind GFP/RFP/LacZ |
| BG20381 | TAACTTTAGTCTCGGTATACTCTTCTGTTTTCG | Spacer 7 FW | Inserting spacer behind GFP/RFP/LacZ |
| BG20382 | GGGCGAAAACAGAAGAGTATACCGAGACTAAAG | Spacer 7 RV | Inserting spacer behind GFP/RFP/LacZ |
| BG29915 | GGCTCCCTCGCACCCCGCTTCGGCGGGGTTTTTTTG | Spacer3 Extended A-tract 1 FW | Inserting modified terminator behind spacer 3 |

| | | | |
|---|---|---|---|
| BG29916 | TTGCAAAAAAACCCCGCCGAAGCGGGGTGCGAGGGA | Spacer3 Extended A-tract 1 RV | Inserting modified terminator behind spacer 3 |
| BG29917 | GGCTCCCTCGCCCCCCGCTTCGGCGGGGGTTTTTTTG | Spacer3 Extended Stem1 GC FW | Inserting modified terminator behind spacer 3 |
| BG29918 | TTGCAAAAAAACCCCCGCCGAAGCGGGGGGCGAGGGA | Spacer3 Extended Stem1 GC RV | Inserting modified terminator behind spacer 3 |
| BG29919 | GGCTCCCTCGCAAACCCCGCTTCGGCGGGGTTTTTTTG | Spacer3 Extended A-tract 3 FW | Inserting modified terminator behind spacer 3 |
| BG29920 | TTGCAAAAAAACCCCGCCGAAGCGGGGTTTGCGAGGGA | Spacer3 Extended A-tract 3 RV | Inserting modified terminator behind spacer 3 |
| BG29921 | GGCTCCCTCGCCCCCCCCGCTTCGGCGGGGGGGTTTTTTTG | Spacer3 Extended Stem3 GC FW | Inserting modified terminator behind spacer 3 |
| BG29922 | TTGCAAAAAAACCCCCCCGCCGAAGCGGGGGGGGCGAGGGA | Spacer3 Extended Stem3 GC RV | Inserting modified terminator behind spacer 3 |
| BG29923 | GGCTCCCTCGCAAAAAAACCCCGCTTCGGCGGGGTTTTTTTG | Spacer3 Extended A-tract 7 FW | Inserting modified terminator behind spacer 3 |
| BG29924 | TTGCAAAAAAACCCCGCCGAAGCGGGGTTTTTTTGCGAGGGA | Spacer3 Extended A-tract 7 RV | Inserting modified terminator behind spacer 3 |
| BG29925 | GGCTCCCTCGCCCCCCCCCCCCGCTTCGGCGGGGGGGGGGGTTTTTTTG | Spacer3 Extended Stem7 GC FW | Inserting modified terminator behind spacer 3 |
| BG29926 | TTGCAAAAAAACCCCCCCCCCCGCCGAAGCGGGGGGGGGGGGCGAGGGA | Spacer3 Extended Stem7 GC RV | Inserting modified terminator behind spacer 3 |
| BG29927 | GCGCTCTTCGCAAGTGGCACTTTTCGGGG | RFP/BB_FW | Linearization backbone + RFP for insertion of modified terminator by Golden Gate (Compatible with BG29915-28) |
| BG29928 | CGGCTCTTCAGCCGGCCGCTCGG | RFP/BB_RV | |
| BG34009 | GATCCGTCTCTGCAAGTGGCACTTTTCGGGGAAATGTGC | RFP/BB_FW | Linearization backbone + RFP for insertion of modified terminator by Golden Gate (Compatible with BG34011-46) |
| BG34010 | GATCCGTCTCTGCGAGGGAGCCGGCCGC | RFP/BB_RV | |
| BG34011 | TCGCAACCCCGCTTCGGCGGGGTTTTTTT | Spacer3 Extended A-tract 2 FW | Inserting modified terminator behind spacer 3 |
| BG34012 | TTGCAAAAAAACCCCGCCGAAGCGGGGTT | Spacer3 Extended A-tract 2 RV | Inserting modified terminator behind spacer 3 |
| BG34013 | TCGCAAAACCCCGCTTCGGCGGGGTTTTTTT | Spacer3 Extended A-tract 4 FW | Inserting modified terminator behind spacer 3 |

| | | | |
|---|---|---|---|
| BG34014 | TTGCAAAAAAACCCCGCCGAAGCGGGGTTTT | Spacer3 Extended A-tract 4 RV | Inserting modified terminator behind spacer 3 |
| BG34015 | TCGCAAAAACCCCGCTTCGGCGGGGTTTTTTT | Spacer3 Extended A-tract 5 FW | Inserting modified terminator behind spacer 3 |
| BG34016 | TTGCAAAAAAACCCCGCCGAAGCGGGGTTTTT | Spacer3 Extended A-tract 5 RV | Inserting modified terminator behind spacer 3 |
| BG34287 | TCGCAAAAAACCCCGCTTCGGCGGGGTTTTTTT | Spacer3 Extended A-tract 6 FW | Inserting modified terminator behind spacer 3 |
| BG34018 | TTGCAAAAAAACCCCGCCGAAGCGGGGTTTTTT | Spacer3 Extended A-tract 6 RV | Inserting modified terminator behind spacer 3 |
| BG34019 | TCGCTCCCCGCTTCGGCGGGGATTTTTTT | Spacer3 Extended Stem1 FW | Inserting modified terminator behind spacer 3 |
| BG34020 | TTGCAAAAAAATCCCCGCCGAAGCGGGGA | Spacer3 Extended Stem1 RV | Inserting modified terminator behind spacer 3 |
| BG34021 | TCGCTTCCCCGCTTCGGCGGGGAATTTTTTT | Spacer3 Extended Stem2 FW | Inserting modified terminator behind spacer 3 |
| BG34022 | TTGCAAAAAAATTCCCCGCCGAAGCGGGGAA | Spacer3 Extended Stem2 RV | Inserting modified terminator behind spacer 3 |
| BG34023 | TCGCTTTCCCCGCTTCGGCGGGGAAATTTTTTT | Spacer3 Extended Stem3 FW | Inserting modified terminator behind spacer 3 |
| BG34024 | TTGCAAAAAAATTTCCCCGCCGAAGCGGGGAAA | Spacer3 Extended Stem3 RV | Inserting modified terminator behind spacer 3 |
| BG34025 | TCGCTTTTCCCCGCTTCGGCGGGGAAAATTTTTTT | Spacer3 Extended Stem4 FW | Inserting modified terminator behind spacer 3 |
| BG34026 | TTGCAAAAAAATTTTCCCCGCCGAAGCGGGGAAAA | Spacer3 Extended Stem4 RV | Inserting modified terminator behind spacer 3 |
| BG34027 | TCGCTTTTTCCCCGCTTCGGCGGGGAAAAATTTTTTT | Spacer3 Extended Stem5 FW | Inserting modified terminator behind spacer 3 |
| BG34028 | TTGCAAAAAAATTTTTCCCCGCCGAAGCGGGGAAAAA | Spacer3 Extended Stem5 RV | Inserting modified terminator behind spacer 3 |
| BG34029 | TCGCTTTTTTCCCCGCTTCGGCGGGGAAAAAATTTTTTT | Spacer3 Extended Stem6 FW | Inserting modified terminator behind spacer 3 |
| BG34030 | TTGCAAAAAAATTTTTTCCCCGCCGAAGCGGGGAAAAAA | Spacer3 Extended Stem6 RV | Inserting modified terminator behind spacer 3 |

| | | | |
|---|---|---|---|
| BG34031 | TCGCTTTTTTTCCCCGCTTCGGCGGGGAAAAAAATTTTTTT | Spacer3 Extended Stem7 FW | Inserting modified terminator behind spacer 3 |
| BG34032 | TTGCAAAAAAATTTTTTTCCCCGCCGAAGCGGGGAAAAAAA | Spacer3 Extended Stem7 RV | Inserting modified terminator behind spacer 3 |
| BG34033 | TCGCCCCCGCTTCGGCGGGGTTTTTT | Spacer3 Shorter U-tract 6 FW | Inserting modified terminator behind spacer 3 |
| BG34034 | TTGCAAAAAACCCCGCCGAAGCGGGG | Spacer3 Shorter U-tract 6 RV | Inserting modified terminator behind spacer 3 |
| BG34035 | TCGCCCCCGCTTCGGCGGGGTTTTT | Spacer3 Shorter U-tract 5 FW | Inserting modified terminator behind spacer 3 |
| BG34036 | TTGCAAAAACCCCGCCGAAGCGGGG | Spacer3 Shorter U-tract 5 RV | Inserting modified terminator behind spacer 3 |
| BG34037 | TCGCCCCCGCTTCGGCGGGGTTTT | Spacer3 Shorter U-tract 4 FW | Inserting modified terminator behind spacer 3 |
| BG34038 | TTGCAAAACCCCGCCGAAGCGGGG | Spacer3 Shorter U-tract 4 RV | Inserting modified terminator behind spacer 3 |
| BG34039 | TCGCCCCCGCTTCGGCGGGGTTT | Spacer3 Shorter U-tract 3 FW | Inserting modified terminator behind spacer 3 |
| BG34040 | TTGCAAACCCCGCCGAAGCGGGG | Spacer3 Shorter U-tract 3 RV | Inserting modified terminator behind spacer 3 |
| BG34041 | TCGCCCCCGCTTCGGCGGGGTT | Spacer3 Shorter U-tract 2 FW | Inserting modified terminator behind spacer 3 |
| BG34042 | TTGCAACCCCGCCGAAGCGGGG | Spacer3 Shorter U-tract 2 RV | Inserting modified terminator behind spacer 3 |
| BG34043 | TCGCCCCCGCTTCGGCGGGGT | Spacer3 Shorter U-tract 1 FW | Inserting modified terminator behind spacer 3 |
| BG34044 | TTGCACCCCGCCGAAGCGGGG | Spacer3 Shorter U-tract 1 RV | Inserting modified terminator behind spacer 3 |
| BG34045 | TCGCCCCCGCTTCGGCGGGG | Spacer3 Shorter U-tract 0 FW | Inserting modified terminator behind spacer 3 |
| BG34046 | TTGCCCCCGCCGAAGCGGGG | Spacer3 Shorter U-tract 0 RV | Inserting modified terminator behind spacer 3 |
| BG36337 | TCACGTCTCCCGAAAAGTGCCACTTGC | Readthrough_FW | Linearization backbone + RFP + 3'UTR for insertion readthrough modules by Golden Gate |

| | | | |
|---|---|---|---|
| BG36336 | TCACGTCTCATGTGCGCGGAACCCC | Readthrough_RV | Linearization backbone + RFP + 3'UTR for insertion readthrough modules by Golden Gate |
| BG36343 | ACACGTCTCTcacattctcaccaataaaaaacgcccg | sfGFP_FW | Amplification + adding GG sited to sfGFP+T0 for readthrough assay |
| BG36341 | ACTCGTCTCgTTCGggtattgtgctagctactagagaaagagg | sfGFP_RV | Amplification + adding GG sited to sfGFP+T0 for readthrough assay |
| BG36338 | TGACGTCTCCTTCGCTCGGTACCAAATCTAACTAAAAAGACGCTGAAAAGCGTCTTTTTT | L3S2P52 terminator FW | Generation of Golden Gate compatible L3S2P52 terminator fragment |
| BG36339 | TCACGTCTCACACAGGACCAAAACGAAAAAAGACGCTTTTCAGCG | L3S2P52 terminator RV | Generation of Golden Gate compatible L3S2P52 terminator fragment |
| BG37684 | TGACGTCTCCTTCGTTGCCATGTGTATGTGGGCGTACGAAGGAAGGTTTGGTATGTGGTATATTCG | MangoIII_FW | Generation of Golden Gate compatible ManoIII-(10AU) fragment |
| BG37685 | TCACGTCTCACACATTGCCATGAATGATCCCGAAGGATCATCAGAGTATGTGGGCGTACGAATATACCACATACCAAACCTTCC | MangoIII_RV | Generation of Golden Gate compatible ManoIII-(10AU) fragment |
| BG37686 | GATCCGTCTCTGCGAGGGAGCCGGCCGC | Spacer3_RV | Primer binding to Spacer 3 for Mango-III positive control |
| BG37692 | TGACGTCTCCTCGCTGCCATGTGTATGTGGGCGTACGAAGGAAGGTTTGGTATGTGGTATATTCG | MangoCTRL_FW | Primer binding toMango-III for Mango-III positive control |
| BG38703 | GCGAAAAAACCCCGCCGAAGCGGGGTTTTTTGCGCAAATAGGGGTTCCGCGCAC | Term_RV | Linearization of RFP-3'UTR-MangoIII fragments and insertion of terminator |
| BG38694 | GTAGCACCTGAAGTCAGCCCCATACG | linear-FW | Linearization of RFP-3'UTR-MangoIII fragments and insertion of terminator |
| BG36332 | GGTGGTCACTACGACGCTGA | RTddPCR_RFP_FW | Primers for RTddPCR with RFP target |
| BG36333 | CGTACTGTTCAACGATGGTGTAGT | RTddPCR_RFP_RV | Primers for RTddPCR with RFP target |
| BG36334 | cgagcgatcatcgaagtctgacc | RTddPCR_fbaA_FW | Primers for RTddPCR with FBA1 target |
| BG36335 | cctgggaaggcgttctgaact | RTddPCR_fbaA_RV | Primers for RTddPCR with FBA1 target |
| BG34791 | /phos/CCAGATCGGAAGAGCGTCGTGTAGGGAAAGAGTGTAGCGCTAGGTGTAGATCTCGGTGGTCGCCGTATCATT/phos/ | Adapter-i5_0001 | i5 barcoded adapter compatibe with Nextera Next-generation sequencing |
| BG35025 | /phos/ CCAGATCGGAAGAGCGTCGTGTAGGGAAAGAGTGTGATATCGAGTGTAGATCTCGGTGGTCGCCGTATCATT/phos/ | Adapter-i5_0002 | i5 barcoded adapter compatibe with Nextera Next-generation sequencing |
| BG35026 | /phos/CCAGATCGGAAGAGCGTCGTGTAGGGAAAGAGTGTCGCAGACGGTGTAGATCTCGGTGGTCGCCGTATCATT/phos/ | Adapter-i5_0003 | i5 barcoded adapter compatibe with Nextera Next-generation sequencing |

| | | | |
|---|---|---|---|
| BG35027 | /phos/CCAGATCGGAAGAGCGTCGTGTAGGGAAAGAGTGTTATGAGTAGTGTAGATCTCGGTGGTCGCCGTATCATT/phos/ | Adapter-i5_0004 | i5 barcoded adapter compatibe with Nextera Next-generation sequencing |
| BG35028 | /phos/CCAGATCGGAAGAGCGTCGTGTAGGGAAAGAGTGTAGGTGCGTGTGTAGATCTCGGTGGTCGCCGTATCATT/phos/ | Adapter-i5_0005 | i5 barcoded adapter compatibe with Nextera Next-generation sequencing |
| BG35029 | /phos/CCAGATCGGAAGAGCGTCGTGTAGGGAAAGAGTGTGAACATACGTGTAGATCTCGGTGGTCGCCGTATCATT/phos/ | Adapter-i5_0006 | i5 barcoded adapter compatibe with Nextera Next-generation sequencing |
| BG35030 | /phos/CCAGATCGGAAGAGCGTCGTGTAGGGAAAGAGTGTACATAGCGGTGTAGATCTCGGTGGTCGCCGTATCATT/phos/ | Adapter-i5_0007 | i5 barcoded adapter compatibe with Nextera Next-generation sequencing |
| BG35031 | /phos/CCAGATCGGAAGAGCGTCGTGTAGGGAAAGAGTGTGTGCGATAGTGTAGATCTCGGTGGTCGCCGTATCATT/phos/ | Adapter-i5_0008 | i5 barcoded adapter compatibe with Nextera Next-generation sequencing |
| BG35032 | /phos/CCAGATCGGAAGAGCGTCGTGTAGGGAAAGAGTGTCCAACAGAGTGTAGATCTCGGTGGTCGCCGTATCATT/phos/ | Adapter-i5_0009 | i5 barcoded adapter compatibe with Nextera Next-generation sequencing |
| BG35033 | /phos/CCAGATCGGAAGAGCGTCGTGTAGGGAAAGAGTGTTTGGTGAGGTGTAGATCTCGGTGGTCGCCGTATCATT/phos/ | Adapter-i5_0010 | i5 barcoded adapter compatibe with Nextera Next-generation sequencing |
| BG34793 | CAAGCAGAAGACGGCATACGAGATCCGCGGTTGTGACTGGAGTTCAGACGTGTGC | Adapter-i7_0001 | i7 barcoded adapter compatibe with Nextera Next-generation sequencing, for PCR on i7 universal adapter |
| BG35034 | CAAGCAGAAGACGGCATACGAGATTTATAACCGTGACTGGAGTTCAGACGTGTGC | Adapter-i7_0002 | i7 barcoded adapter compatibe with Nextera Next-generation sequencing, for PCR on i7 universal adapter |
| BG35035 | CAAGCAGAAGACGGCATACGAGATGGACTTGGGTGACTGGAGTTCAGACGTGTGC | Adapter-i7_0003 | i7 barcoded adapter compatibe with Nextera Next-generation sequencing, for PCR on i7 universal adapter |
| BG35036 | CAAGCAGAAGACGGCATACGAGATAAGTCCAAGTGACTGGAGTTCAGACGTGTGC | Adapter-i7_0004 | i7 barcoded adapter compatibe with Nextera Next-generation sequencing, for PCR on i7 universal adapter |
| BG35037 | CAAGCAGAAGACGGCATACGAGATATCCACTGGTGACTGGAGTTCAGACGTGTGC | Adapter-i7_0005 | i7 barcoded adapter compatibe with Nextera Next-generation sequencing, for PCR on i7 universal adapter |
| BG35038 | CAAGCAGAAGACGGCATACGAGATGCTTGTCAGTGACTGGAGTTCAGACGTGTGC | Adapter-i7_0006 | i7 barcoded adapter compatibe with Nextera Next-generation sequencing, for PCR on i7 universal adapter |

| | | | |
|---|---|---|---|
| BG35039 | CAAGCAGAAGACGGCATACGAGATCAAGCTAGGTGACTGGAGTTCAGACGTGTGC | Adapter-i7_0007 | i7 barcoded adapter compatibe with Nextera Next-generation sequencing, for PCR on i7 universal adapter |
| BG35040 | CAAGCAGAAGACGGCATACGAGATTGGATCGAGTGACTGGAGTTCAGACGTGTGC | Adapter-i7_0008 | i7 barcoded adapter compatibe with Nextera Next-generation sequencing, for PCR on i7 universal adapter |
| BG35041 | CAAGCAGAAGACGGCATACGAGATAGTTCAGGGTGACTGGAGTTCAGACGTGTGC | Adapter-i7_0009 | i7 barcoded adapter compatibe with Nextera Next-generation sequencing, for PCR on i7 universal adapter |
| BG35042 | CAAGCAGAAGACGGCATACGAGATGACCTGAAGTGACTGGAGTTCAGACGTGTGC | Adapter-i7_0010 | i7 barcoded adapter compatibe with Nextera Next-generation sequencing, for PCR on i7 universal adapter |
| BG35054 | CAAGCAGAAGACGGCATACGAGATTCTCTACTGTGACTGGAGTTCAGACGTGTGC | Adapter-i7_0011 | i7 barcoded adapter compatibe with Nextera Next-generation sequencing, for PCR on i7 universal adapter |
| BG35055 | CAAGCAGAAGACGGCATACGAGATCTCTCGTCGTGACTGGAGTTCAGACGTGTGC | Adapter-i7_0012 | i7 barcoded adapter compatibe with Nextera Next-generation sequencing, for PCR on i7 universal adapter |
| BG34792 | /phos/CCAGATCGGAAGAGCACACGTCTGAACTCCAGTCAC /phos/ | i7 universal adapter | i7 universal adapter that i7 barcoded adapters can anneal to |
| BG34794 | AATGATACGGCGACCACCG | RT primer | Primer for first-strand synthesis Term-seq |

**Table S3: Mean Pearson R and Spearman R on cross-validation and test sets.** Note that standard deviation could only be computed with cross-validation sets, as the performance of a model on the test set can only be measured once. CV: cross-validation. s.d.: standard deviation

| Model name | Pearson R (CV) | Spearman R (CV) | Pearson R test | Spearman R test |
|---|---|---|---|---|
| **Term-Seq all** | 0.45 (s.d. = 0.09) | 0.53 (s.d. = 0.05) | 0.48 | 0.53 |
| **Term-Seq E. coli** | 0.32 (s.d. = 0.10) | 0.37 (s.d. = 0.08) | 0.39 | 0.47 |
| **Term-Seq B. subtilis** | 0.40 (s.d. = 0.08) | 0.43 (s.d. = 0.07) | 0.34 | 0.45 |
| **Read-through all** | 0.79 (s.d. = 0.04) | 0.81 (s.d. = 0.05) | 0.73 | 0.76 |
| **Read-through natural** | 0.49 (s.d. = 0.06) | 0.46 (s.d. = 0.11) | 0.42 | 0.53 |
| **Read-through synthetic** | 0.90 (s.d. = 0.05) | 0.89 (s.d. = 0.05) | 0.90 | 0.88 |

**Table S4: Top 10 features per model (trained on the full dataset).** POT: point of termination. Dataset names correspond to names in the first column of Table X.

| **Term-Seq all** | **Term-Seq E. coli** | **Term-Seq B. subtilis** | **Read-through all** | **Read-through natural** | **Read-through synthetic** |
|---|---|---|---|---|---|
| Is the 3rd base of the U-tract a U? (0.077) | Is the 3rd base of the U-tract a U? (0.045) | Is the 4th base of the U-tract a U? (0.034) | Is the 7th base of the spacer a U? (0.144) | Is the 2nd base of the U-tract a U? (0.077) | Is the 8th base of the loop a C? (0.495) |
| Is the 5th base of the U-tract a U? (0.037) | Is the 9th base of the U-tract a U? (0.029) | Is the 11th base of the U-tract the POT? (0.029) | Is the 8th base of the U-tract an A? (0.097) | Is the 3rd base of the U-tract a U? (0.073) | Is the 8th base of the U-tract an A? (0.139) |
| Is the 4th base of the U-tract a U? (0.025) | Is the 6th base of the U-tract a U? (0.024) | Is the 5th base of the U-tract a U? (0.027) | Is the 2nd base of the U-tract a U? (0.054) | Is the 1st base of the U-tract a U? (0.041) | Is the 7th base of the U-tract a C? (0.038) |
| Is the 14th base of the spacer an A? (0.018) | Is the 2nd base of the U-tract a U? (0.017) | Is the 3rd base of the U-tract a U? (0.026) | Is the 4th base of the U-tract a U? (0.036) | Is the 4th base of the U-tract a U? (0.025) | Is the upstream base of the 9th base pair of the stem a G? (0.021) |
| Is the 11th base of the U-tract the POT? (0.015) | Is the 5th base of the U-tract a U? (0.017) | Is the 14th base of the spacer an A? (0.024) | Is the 3rd base of the U-tract a U? (0.021) | Is the 14th base of the loop a C? (0.019) | Is the 8th base of the U-tract a U? (0.013) |
| Is the 15th base of the spacer an A? (0.013) | Is the 8th base of the U-tract a C? (0.016) | Is the 5th base of the U-tract a C? (0.013) | Is the 8th base of the spacer an A? (0.020) | Is the 1st base of the U-tract a C? (0.018) | Is the 5th base of the U-tract a G? (0.011) |
| Is the 9th base of the U-tract a U? (0.011) | Is the 4th base of the U-tract a G? (0.016) | Is the 15th base of the loop an A? (0.012) | Is the 9th base of the U-tract a U? (0.017) | Is the 5th base of the U-tract a U? (0.014) | Is the 8th base of the spacer an A? (0.009) |
| Is the downstream base of the 2nd base pair of the stem a C? (0.010) | Is the 14th base of the U-tract a G? (0.010) | Is the 4th base pair of the stem bonded? (0.012) | Is the 5th base of the U-tract a U? (0.017) | Is the upstream base of the 2nd base pair of the stem a U? (0.014) | Is the downstream base of the 2nd base pair of the stem a G? (0.008) |
| Is the 8th base of the U-tract a C? (0.009) | Is the 4th base of the U-tract a U? (0.009) | Is the 14th base of the spacer a G? (0.010) | Is the 8th base of the U-tract a U? (0.017) | Is the 11th base of the U-tract an A? (0.012) | Is the 8th base of the spacer a C? (0.008) |
| Is the 7th base of the U-tract a U? (0.008) | Is the 12th base of the loop a U? (0.009) | Is the 5th base pair of the stem bonded? (0.009) | Is the 1st base of the U-tract a U? (0.015) | Is the 8th base of the U-tract an A? (0.011) | Is the 9th base of the loop an A? (0.008) |

#### Sequences used in this study

All plasmids contain the same backbone with a Kanamycin resistance (KanR) and a P15A origin of replication, an insulating terminator T2 and a beta-lactamase promoter *pBla*. Through modular cloning, different inserts (see sequences below), 3'UTRs (Supplementary table 1) and readthrough modules (sequences below).

The figure serves as an illustration how different sequences are combined.

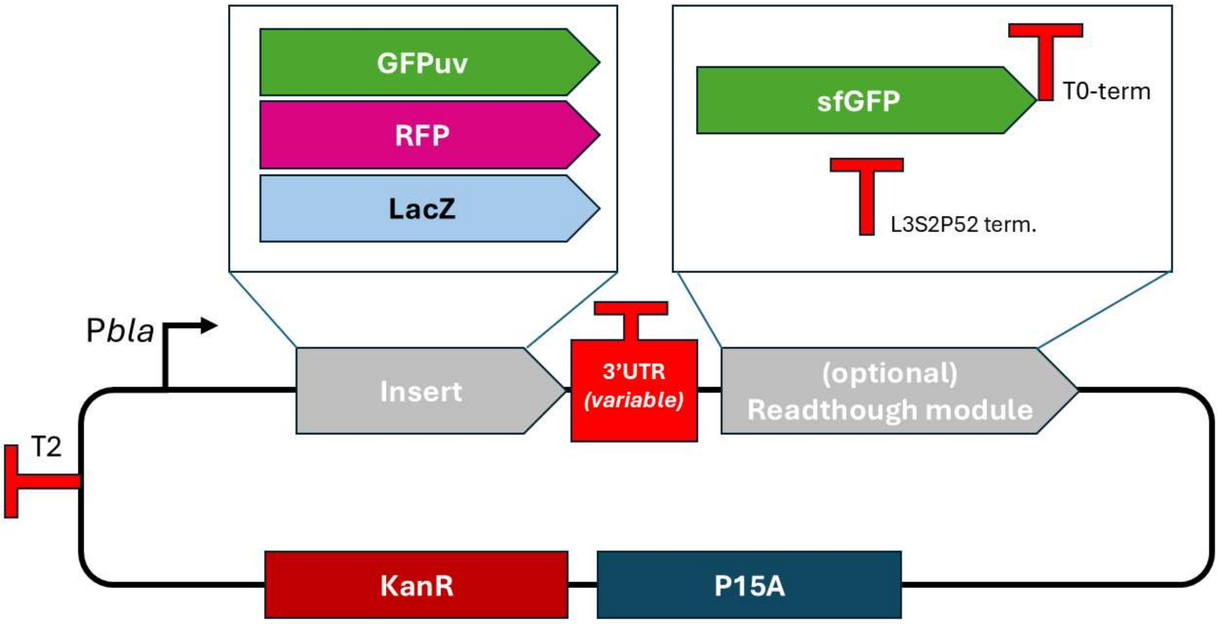


##### Backbone

Pbla is underlined.

```
  1 GCAAGTGGCA CTTTTCGGGG AAATGTGCGC GGAACCCCTA TTTGTTTATT TTTCTAAATA
 61 CATTCAAATA TGTATCCGCT CATGAATTAA TTCTTAGAAA AACTCATCGA GCATCAAATG
121 AAACTGCAAT TTATTCATAT CAGGATTATC AATACCATAT TTTTGAAAAA GCCGTTTCTG
181 TAATGAAGGA GAAAACTCAC CGAGGCAGTT CCATAGGATG GCAAGATCCT GGTATCGGTC
241 TGCGATTCCG ACTCGTCCAA CATCAATACA ACCTATTAAT TTCCCCTCGT CAAAAATAAG
301 GTTATCAAGT GAGAAATCAC CATGAGTGAC GACTGAATCC GGTGAGAATG GCAAAAGTTT
361 ATGCATTTCT TTCCAGACTT GTTCAACAGG CCAGCCATTA CGCTCGTCAT CAAAATCACT
421 CGCATCAACC AAACCGTTAT TCATTCGTGA TTGCGCCTGA CCGAGACGAA ATACGCGGTC
481 GCTGTTAAAA GGACAATTAC AAACAGGAAT CGAATGCAAC CGGCGCAGGA ACACTGCCAG
541 CGCATCAACA ATATTTTCAC CTGAATCAGG ATATTCTTCT AATACCTGGA ATGCTGTTTT
601 CCCGGGGATC GCAGTGGTGA GTAACCATGC ATCATCAGGA GTACGGATAA AATGCTTGAT
```

661 GGTCGGAAGA GGCATAAATT CCGTCAGCCA GTTTAGTCTG ACCATCTCAT CTGTAACATC
721 ATTGGCAACG CTACCTTTGC CATGTTTCAG AAACAACTCT GGCGCATCGG GCTTCCCATA
781 CAATCGATAG ATTGTCGCAC CTGATTGCCC GACATTATCG CGAGCCCATT TATACCCATA
841 TAAATCAGCA TCCATGTTGG AATTTAATCG CGGCCTAGAG CAAGACGTTT CCCGTTGAAT
901 ATGGCTCATA CTCTTCCTTT TTCAATATTA TTGAAGCATT TATCAGGGTT ATTGTCTCAT
961 GAGCGGATAC ATATTTGAAT GTATTTAGAA AAATAAACAA ATAGGCTGTC CCTCCTGTTC
1021 AGCTACTGAC GGGGTGGTGC GTAACGGCAA AAGCACCGCC GGACATCAGC GCTAGCGGAG
1081 TGTATACTGG CTTACTATGT TGGCACTGAT GAGGGTGTCA GTGAAGTGCT TCATGTGGCA
1141 GGAGAAAAAA GGCTGCACCG GTGCGTCAGC AGAATATGTG ATACAGGATA TATTCCGCTT
1201 CCTCGCTCAC TGACTCGCTA CGCTCGGTCG TTCGACTGCG GCGAGCGGAA ATGGCTTACG
1261 AACGGGGCGG AGATTTCCTG GAAGATGCCA GGAAGATACT TAACAGGGAA GTGAGAGGGC
1321 CGCGGCAAAG CCGTTTTTCC ATAGGCTCCG CCCCCCTGAC AAGCATCACG AAATCTGACG
1381 CTCAAATCAG TGGTGGCGAA ACCCGACAGG ACTATAAAGA TACCAGGCGT TTCCCCCTGG
1441 CGGCTCCCTC GTGCGCTCTC CTGTTCCTGC CTTTCGGTTT ACCGGTGTCA TTCCGCTGTT
1501 ATGGCCGCGT TTGTCTCATT CCACGCCTGA CACTCAGTTC CGGGTAGGCA GTTCGCTCCA
1561 AGCTGGACTG TATGCACGAA CCCCCCGTTC AGTCCGACCG CTGCGCCTTA TCCGGTAACT
1621 ATCGTCTTGA GTCCAACCCG GAAAGACATG CAAAAGCACC ACTGGCAGCA GCCACTGGTA
1681 ATTGATTTAG AGGAGTTAGT CTTGAAGTCA TGCGCCGGTT AAGGCTAAAC TGAAAGGACA
1741 AGTTTTGGTG ACTGCGCTCC TCCAAGCCAG TTACCTCGGT TCAAAGAGTT GGTAGCTCAG
1801 AGAACCTTCG AAAAACCGCC CTGCAAGGCG GTTTTTTCGT TTTCAGAGCA AGAGATTACG
1861 CGCAGACCAA AACGATCTCA AGAAGATCAT CTTATTAATC AGATAAAATA TTTCTAGATT
1921 TCAGTGCAAT TTATCTCTTC AAATGTAGCA CCTGAAGTCA GCCCCATACG ATATAAGTTG
1981 TAATTCGGTA CCCCGCTTCG GCGGGGTTTT TTCAAGTTCA AATATGTATC CGCTCATGAG
2041 ACAATGTGTG GGGAGACCAC AACGGTTTCC CTCTAGAAAT AATTTTGTTT AACTATAAGA
2101 AGGAGATATA CAT
[INSERT] [3’UTR] [Optional: readthrough]

#### Insert sequences

##### *GFPuv*

1 ATGAGTAAAG GAGAAGAACT TTTCACTGGA GTTGTCCCAA TTCTTGTTGA ATTAGATGGT
61 GATGTTAATG GGCACAAATT TTCTGTCAGT GGAGAGGGTG AAGGTGATGC AACATACGGA
121 AAACTTACCC TTAAATTTAT TTGCACTACT GGAAAACTAC CTGTTCCATG GCCAACACTT
181 GTCACTACTT TCTCTTATGG TGTTCAATGC TTTTCCCGTT ATCCGGATCA CATGAAACGG
241 CATGACTTTT TCAAGAGTGC CATGCCCGAA GGTTATGTAC AGGAACGCAC TATATCTTTC

301 AAAGATGACG GGAACTACAA GACGCGTGCT GAAGTCAAGT TTGAAGGTGA TACCCTTGTT
361 AATCGTATCG AGTTAAAAGG TATTGATTTT AAAGAAGATG GAAACATTCT CGGACACAAA
421 CTGGAGTACA ACTATAACTC ACACAATGTA TACATCACGG CAGACAAACA AAAGAATGGA
481 ATCAAAGCTA ACTTCAAAAT TCGCCACAAC ATTGAAGATG GATCCGTTCA ACTAGCAGAC
541 CATTATCAAC AAAATACTCC AATTGGCGAT GGCCCTGTCC TTTTACCAGA CAACCATTAC
601 CTGTCGACAC AATCTGCCCT TTCGAAAGAT CCCAACGAAA AGCGTGACCA CATGGTCCTT
661 CTTGAGTTTG TAACTGCTGC TGGGATTACA CATGGCATGG ATGAGCTCTA CAAATAA

[3’UTR][Backbone]

*RFP*

1 ATGGCTTCCT CCGAAGACGT TATCAAAGAG TTCATGCGTT TCAAAGTTCG TATGGAAGGT
61 TCCGTTAACG GTCACGAGTT CGAAATCGAA GGTGAAGGTG AAGGTCGTCC GTACGAAGGT
121 ACACAGACCG CTAAACTGAA AGTTACCAAA GGTGGCCCGC TGCCGTTCGC TTGGGACATC
181 CTGTCCCCGC AGTTCCAGTA CGGTTCCAAA GCTTACGTTA AACACCCGGC TGACATCCCG
241 GACTACCTGA AACTGTCCTT CCCGGAAGGT TTCAAATGGG AACGTGTTAT GAACTTCGAA
301 GACGGTGGTG TTGTTACCGT TACCCAGGAC TCCTCCCTGC AAGACGGTGA GTTCATCTAC
361 AAAGTTAAAC TGCGTGGTAC CAACTTCCCG TCCGACGGTC CGGTTATGCA GAAAAAAACC
421 ATGGGTTGGG AAGCTTCCAC CGAACGTATG TACCCGGAAG ACGGTGCTCT GAAAGGTGAA
481 ATCAAAATGC GTCTGAAACT GAAAGACGGT GGTCACTACG ACGCTGAAGT TAAAACCACC
541 TACATGGCTA AAAAACCGGT TCAGCTGCCG GGTGCTTACA AAACCGACAT CAAACTGGAC
601 ATCACCTCCC ACAACGAAGA CTACACCATC GTTGAACAGT ACGAACGTGC TGAAGGTCGT
661 CACTCCACCG GTGCTTAA

[3’UTR][Backbone]

*LacZ*

1 ATGACCATGA TTACGGATTC ACTGGCCGTC GTTTTACAAC GTCGTGACTG GGAAAACCCT
61 GGCGTTACCC AACTTAATCG CCTTGCAGCA CATCCCCCTT TCGCCAGCTG GCGTAATAGC
121 GAAGAGGCCC GCACCGATCG CCCTTCCCAA CAGTTGCGCA GCCTGAATGG CGAATGGCGC
181 TTTGCCTGGT TTCCGGCACC AGAAGCGGTG CCGGAAAGCT GGCTGGAGTG CGATCTTCCT
241 GAGGCCGATA CTGTCGTCGT CCCCTCAAAC TGGCAGATGC ACGGTTACGA TGCGCCCATC
301 TACACCAACG TGACCTATCC CATTACGGTC AATCCGCCGT TTGTTCCCAC GGAGAATCCG
361 ACGGGTTGTT ACTCGCTCAC ATTTAATGTT GATGAAAGCT GGCTACAGGA AGGCCAGACG
421 CGAATTATTT TTGATGGCGT TAACTCGGCG TTTCATCTGT GGTGCAACGG GCGCTGGGTC
481 GGTTACGGCC AGGACAGTCG TTTGCCGTCT GAATTTGACC TGAGCGCATT TTTACGCGCC
541 GGAGAAAACC GCCTCGCGGT GATGGTGCTG CGCTGGAGTG ACGGCAGTTA TCTGGAAGAT

601 CAGGATATGT GGCGGATGAG CGGCATTTTC CGTGACGTCT CGTTGCTGCA TAAACCGACT
661 ACACAAATCA GCGATTTCCA TGTTGCCACT CGCTTTAATG ATGATTTCAG CCGCGCTGTA
721 CTGGAGGCTG AAGTTCAGAT GTGCGGCGAG TTGCGTGACT ACCTACGGGT AACAGTTTCT
781 TTATGGCAGG GTGAAACGCA GGTCGCCAGC GGCACCGCGC CTTTCGGCGG TGAAATTATC
841 GATGAGCGTG GTGGTTATGC CGATCGCGTC ACACTACGTC TGAACGTCGA AAACCCGAAA
901 CTGTGGAGCG CCGAAATCCC GAATCTCTAT CGTGCGGTGG TTGAACTGCA CACCGCCGAC
961 GGCACGCTGA TTGAAGCAGA AGCCTGCGAT GTCGGTTTCC GCGAGGTGCG GATTGAAAAT
1021 GGTCTGCTGC TGCTGAACGG CAAGCCGTTG CTGATTCGAG GCGTTAACCG TCACGAGCAT
1081 CATCCTCTGC ATGGTCAGGT CATGGATGAG CAGACGATGG TGCAGGATAT CCTGCTGATG
1141 AAGCAGAACA ACTTTAACGC CGTGCGCTGT TCGCATTATC CGAACCATCC GCTGTGGTAC
1201 ACGCTGTGCG ACCGCTACGG CCTGTATGTG GTGGATGAAG CCAATATTGA AACCCACGGC
1261 ATGGTGCCAA TGAATCGTCT GACCGATGAT CCGCGCTGGC TACCGGCGAT GAGCGAACGC
1321 GTAACGCGAA TGGTGCAGCG CGATCGTAAT CACCCGAGTG TGATCATCTG GTCGCTGGGG
1381 AATGAATCAG GCCACGGCGC TAATCACGAC GCGCTGTATC GCTGGATCAA ATCTGTCGAT
1441 CCTTCCCGCC CGGTGCAGTA TGAAGGCGGC GGAGCCGACA CCACGGCCAC CGATATTATT
1501 TGCCCGATGT ACGCGCGCGT GGATGAAGAC CAGCCCTTCC CGGCTGTGCC GAAATGGTCC
1561 ATCAAAAAAT GGCTTTCGCT ACCTGGAGAG ACGCGCCCGC TGATCCTTTG CGAATACGCC
1621 CACGCGATGG GTAACAGTCT TGGCGGTTTC GCTAAATACT GGCAGGCGTT TCGTCAGTAT
1681 CCCCGTTTAC AGGGCGGCTT CGTCTGGGAC TGGGTGGATC AGTCGCTGAT TAAATATGAT
1741 GAAAACGGCA ACCCGTGGTC GGCTTACGGC GGTGATTTTG GCGATACGCC GAACGATCGC
1801 CAGTTCTGTA TGAACGGTCT GGTCTTTGCC GACCGCACGC CGCATCCAGC GCTGACGGAA
1861 GCAAAACACC AGCAGCAGTT TTTCCAGTTC CGTTTATCCG GGCAAACCAT CGAAGTGACC
1921 AGCGAATACC TGTTCCGTCA TAGCGATAAC GAGCTCCTGC ACTGGATGGT GGCGCTGGAT
1981 GGTAAGCCGC TGGCAAGCGG TGAAGTGCCT CTGGATGTCG CTCCACAAGG TAAACAGTTG
2041 ATTGAACTGC CTGAACTACC GCAGCCGGAG AGCGCCGGGC AACTCTGGCT CACAGTACGC
2101 GTAGTGCAAC CGAACGCGAC CGCATGGTCA GAAGCCGGGC ACATCAGCGC CTGGCAGCAG
2161 TGGCGTCTGG CGGAAAACCT CAGTGTGACG CTCCCCGCCG CGTCCCACGC CATCCCGCAT
2221 CTGACCACCA GCGAAATGGA TTTTTGCATC GAGCTGGGTA ATAAGCGTTG GCAATTTAAC
2281 CGCCAGTCAG GCTTTCTTTC ACAGATGTGG ATTGGCGATA AAAAACAACT GCTGACGCCG
2341 CTGCGCGATC AGTTCACCCG TGCACCGCTG GATAACGACA TTGGCGTAAG TGAAGCGACC
2401 CGCATTGACC CTAACGCCTG GGTCGAACGC TGGAAGGCGG CGGGCCATTA CCAGGCCGAA
2461 GCAGCGTTGT TGCAGTGCAC GGCAGATACA CTTGCTGATG CGGTGCTGAT TACGACCGCT
2521 CACGCGTGGC AGCATCAGGG GAAAACCTTA TTTATCAGCC GGAAAACCTA CCGGATTGAT
2581 GGTAGTGGTC AAATGGCGAT TACCGTTGAT GTTGAAGTGG CGAGCGATAC ACCGCATCCG

2641 GCGCGGATTG GCCTGAACTG CCAGCTGGCG CAGGTAGCAG AGCGGGTAAA CTGGCTCGGA

2701 TTAGGGCCGC AAGAAAACTA TCCCGACCGC CTTACTGCCG CCTGTTTTGA CCGCTGGGAT

2761 CTGCCATTGT CAGACATGTA TACCCCGTAC GTCTTCCCGA GCGAAAACGG TCTGCGCTGC

2821 GGGACGCGCG AATTGAATTA TGGCCCACAC CAGTGGCGCG GCGACTTCCA GTTCAACATC

2881 AGCCGCTACA GTCAACAGCA ACTGATGGAA ACCAGCCATC GCCATCTGCT GCACGCGGAA

2941 GAAGGCACAT GGCTGAATAT CGACGGTTTC CACATGGGGA TTGGTGGCGA CGACTCCTGG

3001 AGCCCGTCAG TATCGGCGGA ATTCCAGCTG AGCGCCGGTC GCTACCATTA CCAGTTGGTC

3061 TGGTGTCAAA AATAA

[3’UTR][Backbone]

#### Readthrough parts

##### *eGFP + T0t*

eGFP is indicated in green, the T0 terminator is underlined.

[Insert][3’UTR]

1 GCAAGTGGCA CTTTTCGGGT ATTGTGCTAG CTACTAGAGA AAGAGGAGAA ATACTAGATG

61 CGTAAAGGCG AAGAACTGTT CACGGGCGTA GTTCCGATTC TGGTCGAGCT GGACGGCGAT

121 GTGAACGGTC ATAAGTTTAG CGTTCGCGGT GAAGGTGAGG GCGACGCGAC CAACGGCAAA

181 CTGACCCTGA AGTTCATCTG CACCACCGGT AAACTGCCGG TGCCTTGGCC GACCTTGGTG

241 ACGACGTTGA CGTATGGCGT GCAGTGTTTT GCGCGTTATC CGGACCACAT GAAACAACAC

301 GATTTCTTCA AATCTGCGAT GCCGGAGGGT TACGTCCAGG AGCGTACCAT TTCCTTCAAG

361 GATGATGGCT ACTACAAAAC TCGCGCAGAG GTTAAGTTTG AAGGTGACAC GCTGGTCAAT

421 CGTATCGAAT TGAAGGGTAT CGACTTTAAA GAGGATGGTA ACATTCTGGG CCATAAACTG

481 GAGTATAACT TCAACAGCCA TAATGTTTAC ATTACGGCAG ACAAGCAAAA GAACGGCATC

541 AAGGCCAATT TCAAGATTCG CCACAATGTT GAGGACGGTA GCGTCCAACT GGCCGACCAT

601 TACCAGCAGA ACACCCCAAT TGGTGACGGT CCGGTTTTGC TGCCGGATAA TCACTATCTG

661 AGCACCCAAA GCGTGCTGAG CAAAGATCCG AACGAAAAAC GTGATCACAT GGTCCTGCTG

721 GAATTTGTGA CCGCTGCGGG CATCACCCAC GGTATGGATG AACTGTACAA ATAATAATAC

781 TAGTAGCGGC CGCTGCAGTC CGGCAAAAAA GGGCAAGGTC TTGGACTCCT GTTGATAGAT

841 CCAGTAATGA CCTCAGAACT CCATCTGGAT TTGTTCAGAA CGCTCGGTTG CCGCCGGGCG

901 TTTTTTATTG GTGAGA

[Backbone]

##### *L3S2P52 terminator*

The L3S2P52 terminator is underlined

[Insert] [3’UTR]

1 GCAAGTGGCA CTTTTCGCTC GGTACCAAAT CTAACTAAAA AGACGCTGAA AAGCGTCTTT

61 TTTCGTTTTG GTCC

[Backbone]

##### *Mango-III(10AU) constructs*

Mango-III(10AU) in vitro transcription was performed on a linear template containing p*bla*-RFP-3’UTR-Mango-III(10AU)-stabilizing terminator. In the (+) control, the 3’UTR was omitted to ensure constitutive Mango-III transcription.

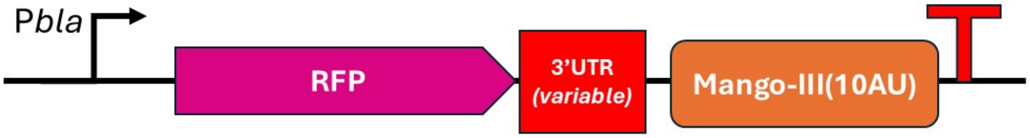


MangoIII(10AU) is highlighted, the terminator underlined.

[pbla][RFP][3’UTR]

1 GCAAGTGGCA CTTTTCGTTG CCATGTGTAT GTGGGCGTAC GAAGGAAGGT TTGGTATGTG

61 GTATATTCGT ACGCCCACAT ACTCTGATGA TCCTTCGGGA TCATTCATGG CAATGTGCGC

121 GGAACCCCTA TTTGCGCAAA AAACCCCGCT TCGGCGGGGT TTTTTCGC
